# Latent plasticity of the human pancreas across development, health, and disease

**DOI:** 10.1101/2025.10.01.679230

**Authors:** Elisabetta Mereu, Diego Balboa, Johannes Liebig, Aitor Gonzalez-Herrero, Anna Martinez-Casals, Mariya Mardamshina, Fanny Mollandin, Felix Schicktanz, Luca Tosti, Valerie Vandenbempt, Dana Avrahami, Edgar Bernardo, Frida Björklund, Robert Lorenz Chua, Marten Engelse, Javier García-Hurtado, Nathalie Groen, Maaike Hanegraaf, Pablo Iañez, Katharina Jechow, Björn Konukiewitz, Christian Lawerenz, Domenica Marchese, Mauro J. Muraro, Silvia Pellegrini, Valeria Sordi, Alexander Sudy, Ulrike Taron, Foo Wei Ten, Timo Trefzer, Sven Twardziok, Maarten van Agen, Françoise Carlotti, Eelco de Koning, Jorge Ferrer, Benjamin Glaser, Holger Heyn, Emma Lundberg, Lorenzo Piemonti, Katja Steiger, Alexander van Oudenaarden, Wilko Weichert, Christian Conrad, Roland Eils

## Abstract

The pancreas plays a central role in major human diseases, yet our understanding of its cellular diversity and plasticity remains incomplete. Here, we present a single-cell multiomics atlas of the human pancreas, profiling over four million cells and nuclei from 57 donors across fetal development, adult homeostasis, and type 2 diabetes (T2D). Integrating sc/snRNA-seq, snATAC-seq, VASA-seq, spatial transcriptomics (Xenium), and multiplexed proteomics (CODEX), we resolve gene expression, chromatin accessibility, and spatial organization at high resolution. We identify transcriptionally plastic centroacinar-like cells (pCACs) in adults with fetal-like features, delineate endocrine and exocrine lineage trajectories during development, and uncover HNF1A-defined beta cell epigenetic states. In T2D, we observe shifts in beta cell subtypes and altered regulatory programs. Glucose perturbation of healthy islets reveals cell-type-specific adaptation and stress responses. This atlas provides a foundational framework to understand pancreas biology and the role of cellular plasticity in regeneration and disease.

---

The pancreas is a vital organ with both exocrine and endocrine functions, essential for proper digestive processes and glucose regulation in the body. The stability of its functions is sustained by limited cell turnover, primarily through self-duplication of pancreatic cells, a process that has been demonstrated in mice and raises the possibility of a similar mechanism in humans (1–4). The identity of pancreatic cells is regulated by changes in the expression of specific genes and transcription factors (5), which are established during embryonic development and maturation and appear to be conserved between mice and humans (6, 7). This dynamic gene expression allows the pancreas to adapt to changes in its functions and preserve homeostasis throughout the lifespan. Experimental studies have shown that adult pancreatic acinar cells can, under certain conditions, undergo epigenomic remodeling and adopt progenitor-like features. For instance, forced expression of Pdx1, MafA, and Ngn3 can reprogram acinar cells into insulin-producing beta-like cells (8), while injury-induced Notch signaling can drive dedifferentiation toward a progenitor-like state (9). Such plasticity might even predispose to neoplastic evolution in the presence of oncogenic drivers (10). Although these findings support the concept of acinar plasticity, the extent and physiological relevance of such processes remain incompletely understood. In this regard, it is crucial to systematically delineate the cellular and molecular landscape of the developing, homeostatic, and diseased human pancreas. Toward this goal, efforts to build a more complete human cell atlas of the pancreas benefit from integrating transcriptomic and epigenetic profiling with spatial information on its heterogeneous cytoarchitecture, as well as relevant anatomical and physiological context, to advance our understanding of the cellular heterogeneity and dynamics of this organ. Single-cell genomics has been applied to investigate the cellular heterogeneity of the human pancreas, particularly within the islets of Langerhans (11–14), but these studies have faced important limitations in accurately defining the full spectrum of heterogeneous cell types. Technical challenges include i) the high enzymatic activity of the pancreas, which makes the isolation of exocrine cells difficult, ii) variability in experimental techniques, resulting in a lack of marker consensus between individual datasets (15), iii) lack of adjacent cytoarchitecture spatial information. In addition to these limitations, previous studies on the cellular heterogeneity of the human pancreas have been restricted to single-cell or single-nucleus RNA sequencing (sc/nRNA-seq), which provides valuable information on the transcriptomic identity of individual cells but does not capture their epigenetic states or spatial context. As epigenomic mechanisms are essential for maintaining cell identity (7, 16, 17), regulating plasticity, and mediating responses to stress and injury (10), integrating epigenomic profiling with transcriptomic and spatial approaches facilitates deeper insights into pancreatic cell differentiation, adaptation, and dysfunction. Such integrative maps ultimately facilitate the development of new strategies for treating pancreatic diseases such as diabetes, pancreatitis, and pancreatic cancer.

In this study, we combined advanced single-cell sequencing techniques, together with spatial transcriptomics (Xenium) and proteomics (CODEX), to create a single-cell multiomics reference atlas of the human pancreas (Fig. 1), including early prenatal life, adulthood, and Type 2 diabetic (T2D) samples, consisting of 57 patient donors. To generate this resource, we processed adult pancreatic tissue from four anatomically distinct regions: head, PPY-rich head, body, and tail. Notably, while many donors displayed histologically normal pancreatic tissue, pathologists also identified features such as fibrosis, lipomatosis, and pancreatic intraepithelial neoplasias (PanINs) in a subset of samples, presenting a view of pancreatic cellular states across physiological and early pathological contexts. Additionally, analysis of human embryonic pancreas samples from week 13 to 17 post-conception (wpc) for single-cell transcriptomics and from 10 to 18 wpc for proteomics allowed us to assess cell fate determination and differentiation during development, and to compare these data with adult pancreas data.

**Figure 1.**
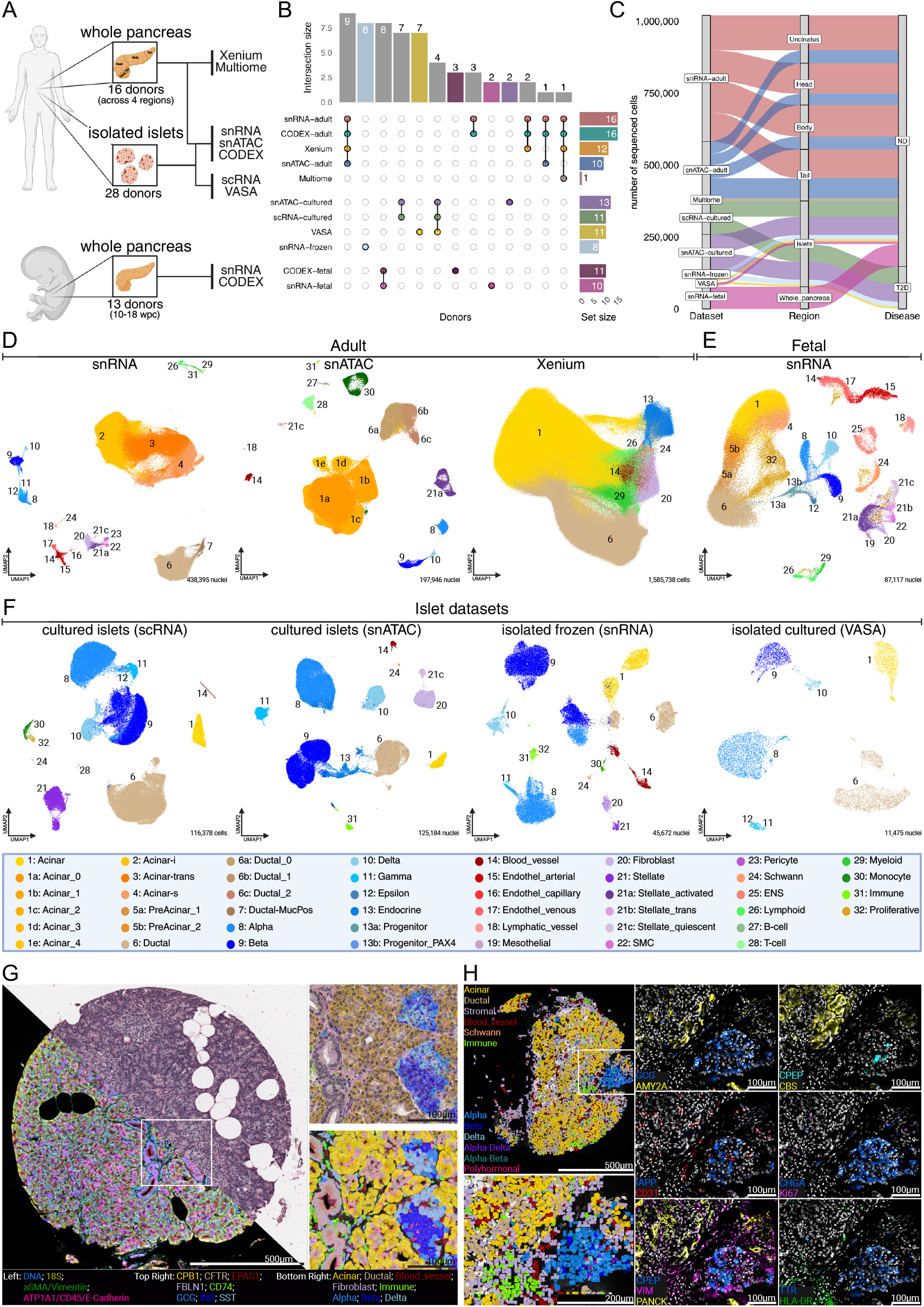
Overview of ESPACE Human Pancreas Cell Atlas. **A**. Summary on the number of included donors and used technologies. **B**. Upsetplot showing how many donors are within each method and how many are shared across different methods. **C**. Sankey plot showing the distribution of cells/ nulcei from all scRNA/snRNA experiments across datasets, pancreatic regions and disease. **D**. UMAP representation of the adult datasets including the snRNA dataset (left), snATAC (middle), xenium (right). Colors and numbers are indicating cell identity listed in the table below. **E**. UMAP representation of the fetal snRNA dataset. Colors and numbers are indicating cell identity listed in the table below. **F**. UMAP representation of the islet datasets including the scRNA dataset (left), snATAC (middle left), snRNA (middle right) and VASA-seq (right). Colors and numbers are indicating cell identity listed in the table below. **G**. Visualization of Xenium data. The main panel (left) shows a representative TMA core half-stained with H&E and half with immunofluorescence (IF). The white square marks the zoomed-in region displayed on the right: transcript counts overlaid on the H&E image (top right) and cell type labels on the IF image (bottom right). Colors correspond to transcripts or cell types, as indicated in the legend below. (Visualization generated with 10x Genomics Xenium Explorer 3.2.0) **H**. Visualization of CODEX data. The main panel (top left) shows a pancreas section with cell type labels. The white square marks a zoomed-in region, shown below and to the right: cell type labels overlaid on the nuclei image (bottom left), and selected protein stainings displayed over the nuclei image (panels on the right). Label and protein stain colors are indicated in the images. This figure was created using BioRender.com

This multiomic resource exemplifies the following key insights:

1. dentification of rare centroacinar, enriched for acinar-to-ductal metaplasia (ADM) and multipotent progenitor markers.
2. Identification of very rare subpopulations, such as novel ionocyte-like cells within the MUC+ ductal cells, and epsilon cells within the endocrine compartment.
3. dentification of islet endocrine cell heterogeneity, including two epigenetic beta cell states (HNF1A_high and HNF1A_low), with a decreased proportion of HNF1A_high in T2D.
4. dentification of consensus dysregulated markers in T2D beta cells, showing altered expression of beta cell transcription factors, including reduced expression of HNF1A target genes.
5. D lineation of *NEUROG3*+ endocrine progenitors differentiation trajectory into major endocrine cell types.

## Results

### ESPACE – the Expression and Spatial Analysis Pancreas Atlas in numbers

The ESPACE atlas provides a multimodal, high-resolution map of the human pancreas across developmental stages, anatomical regions, and disease states. We integrated data from 154 samples, including adult whole pancreas (64 samples from 16 donors across four anatomical regions), isolated islets (80 samples profiled using four distinct assay types from 28 donors), and fetal pancreas (13 donors from gestational weeks 10–18). We employed ten complementary modalities, encompassing transcriptomic (snRNA-seq, scRNA-seq, VASA-seq), epigenomic (snATAC-seq, 10x Multiome), and spatial platforms (Xenium spatial transcriptomics and CODEX high-plex immunofluorescence), with overlapping donors across *multiple datasets (Supp. Table* 1). Together, these approaches captured over 4 million cells and nuclei, enabling robust cross-platform validation of spatial, proteomic, and transcriptional signatures in both adult and developing pancreatic tissue.

### Molecular definition of adult human pancreatic cell types using single-nuclei transcriptomics and chromatin accessibility

To comprehensively dissect the cellular heterogeneity of the adult human pancreas, we employed droplet-based transcriptomics assays snRNA-seq and single-nuclei ATAC-seq (snATAC-seq) for chromatin profiling of a broad range of tissue samples collected from four pancreas regions, namely uncinate process (U), head (H), body (B) and tail (T) (Fig. 1A-C, Supp. Fig. 1A-D, Supp. Table 1).

To ensure adequate representation of acinar and ductal cells, we relied on protocol specifics (18), which generates a suspension of intact nuclei from tissue biopsies, using citric acid, inhibiting the autolytic activity of enzymes and thus drastically improves mRNA quality especially from epithelial cells.

Using snRNA-seq, we first aimed to identify pancreatic cell populations and characterize them by their specific gene markers. To guarantee the high quality of cells and facilitate cell type identification, we implemented stringent quality control procedures (see Methods), including the exclusion of doublets and the removal of ambient RNA, resulting in a reduction of background noise and an improvement in data quality. After integrating the samples and harmonizing gene expression (see Methods), we analyzed 438,395 high-quality nuclei across 64 samples from 16 healthy donors (Fig. 1D). Using a combination of previously established (13, 15) gene markers, we first identified the major pancreatic cell types (Supp. Fig. 2A, Supp. Fig. 1E-F, Supp. Table 2). Next, we conducted a comprehensive analysis of the pancreatic cell compartments, including acinar (Supp. Fig. 2B), ductal (Supp. Fig. 2C), and endocrine cells (Supp. Fig. 2D), to uncover 21 major cell types and 34 additional cell states that contribute to understanding pancreas cell heterogeneity. This indepth examination revealed the high level of complexity and granularity within our pancreatic cell atlas, emphasizing its exceptional sensitivity to capture even the most infrequent cell populations, exemplified by the endocrine epsilon cells (Supp. Fig. 1F), which were represented by only 70 cells in our dataset (0.015%) and the 21 ionocyte-like cells (Supp. note 1) within the ductal compartment (0.0064%) (Supp. Fig. 4).

Although acinar (*RBPJL, PRSS1*) and ductal cells (*CFTR, MUC5B*) comprise almost 90% of all analyzed cells (Supp. Fig. 2G), followed by the endocrine cells with less than 2% (I*NS, GCG, SST, PPY*), we uncovered a diverse cellular landscape within the pancreatic tissue, including stellate cells (*ACTA2, PDZRN4*), Schwann cells (*SCN7A, CDH19*), fibroblasts (*COL6A3, COL4A1*), blood vessels, and endothelial cells (*FLT1, VWF, CDH5*), as well as immune cells, both lymphoid and myeloid (*CD3E, CD163*), which were consistently detected in all the patient donors (Supp. Fig. 2G-H).

Our findings confirmed the existence of previously described acinar cell subtypes (13), with secreting Acinar (Acinar-s) cells expressing *AMY2A, CLPS*, and *CELA2B* and idling Acinar (Acinar-i) cells characterized by the expression of *INSR, FOXP2, RBPJL* and *MAP3K5* (Supp. Fig. 2A). Furthermore, we identified a distinct cluster of transitioning cells (referred to as Acinar-trans in Fig. 1D), which were marked by the expression of *RBPJL, LRIG1* and *REG3G* (Supp. Fig. 2A), clearly representing an intermediate state between the Acinar-i and Acinar-s subtypes (Supp. Fig. 2E). The central cluster, Acinar_7, was particularly interesting, as it contained a distinct subset of cells (Supp. Fig. 2B), exhibiting a hybrid acinar–ductal phenotype, characterized by high expression of ductal markers such as *CFTR, BICC1* and *SLC4A4*, reminiscent of centroacinar cells (CACs). Similarly, within the ductal population, we identified previously described main groups (13), including the conventional Ductal (*CFTR, SLC4A4, BICC1, PDE3A, PKHD1*) and Ductal-Muc+ (*KRT19, MUC5B, MUC1*) along with a distinct subpopulation, Ductal_4, which expressed both ductal and acinar markers (Supp. Fig. 2C, 2F), suggesting plasticity within the exocrine compartment.

### Chromatin accessibility profiling and integration with snRNA-seq data

To further characterize the molecular profiles of these pancreatic cell groups, we analyzed the chromatin accessibility of 197,946 high-quality nuclei from 10 donors previously analyzed in snRNA-seq (Fig. 1B-C). Unsupervised clustering identified 19 diverse cell groups consistently across donors (Fig. 1D, Supp. Fig. 3A), whose identities were explored in two ways. First, we transformed the binary snATAC-seq data into gene activity (GA) counts, quantifying gene scores by integrating chromatin accessibility signals from promoters and gene body regions and creating a new gene-by-cell matrix from the snATAC-seq data. We then annotated the main populations based on cell-type specific markers already identified in the snRNA-seq counterpart (Supp. Fig. 3B). While the main pancreatic cell populations were readily identified, distinguishing between acinar and ductal states was more challenging, as their gene activity scores derived by snATAC-seq data showed strong similarities. Nonetheless, the Acinar_4 cluster in this modality, exhibited a clear correspondence in expression pattern to Acinar_7 cluster in the snRNA-seq data, characterized by co-expression of acinar and ductal markers (Supp. Fig. 3B-C). Secondly, we leveraged an additional single-cell Multiome dataset (Fig. 1B-C, Supp. Fig. 3D) to better link the snRNA-seq and snATAC-seq profiles in pancreatic populations and to generate a unified embedding of both modalities (see Methods).

In this integrated space, the harmonization of both modalities (see Methods) yielded strong concordance for the major cell populations, and we observed consistent expression patterns of key acinar and ductal markers (Fig. 2A-B, Supp. Fig. 3E). To refine this integration and infer the most likely correspondences between transcriptional and chromatin-accessible cell states, we trained a deep neural network–based model using scOMM (19), using the Multiome data as reference. In this way, the model learned the joint transcriptional and chromatin-accessibility profiles of distinct pancreatic cell types and was then used to map adult snATAC-seq profiles onto this reference space. On a held-out validation set, scOMM achieved 95% accuracy, leveraging the matched RNA and ATAC modalities to reliably infer cell identities. While direct validation is not possible for standalone snATAC-seq datasets, similar predictive accuracy is expected given the shared signal structure. Mapping the adult snATAC-seq data onto this multi-omic reference enabled robust identification of major pancreatic cell types (Supp. Fig. 3E) and revealed a comparable degree of heterogeneity within acinar and ductal populations, consistent with the snRNA-seq data (Fig. 2C). At the same time, the chromatin accessibility profiles uncovered additional complexity: several snATAC-seq clusters, particularly within the acinar and ductal compartments, exhibited only partial correspondence to snRNA-seq-defined populations. This suggests the presence of intermediate or transitional cell states not fully captured by transcriptional profiles alone. Also with this approach, the Acinar_4 cluster from the snATAC-seq data showed the strongest match with the ductal phenotype (Fig. 2C, Supp. Fig. 3F), further supporting that a subset of these nuclei mirrors the transcriptional profile of Acinar_7 within the Acinar-trans group identified in the snRNA-seq analysis.

**Figure 2.**
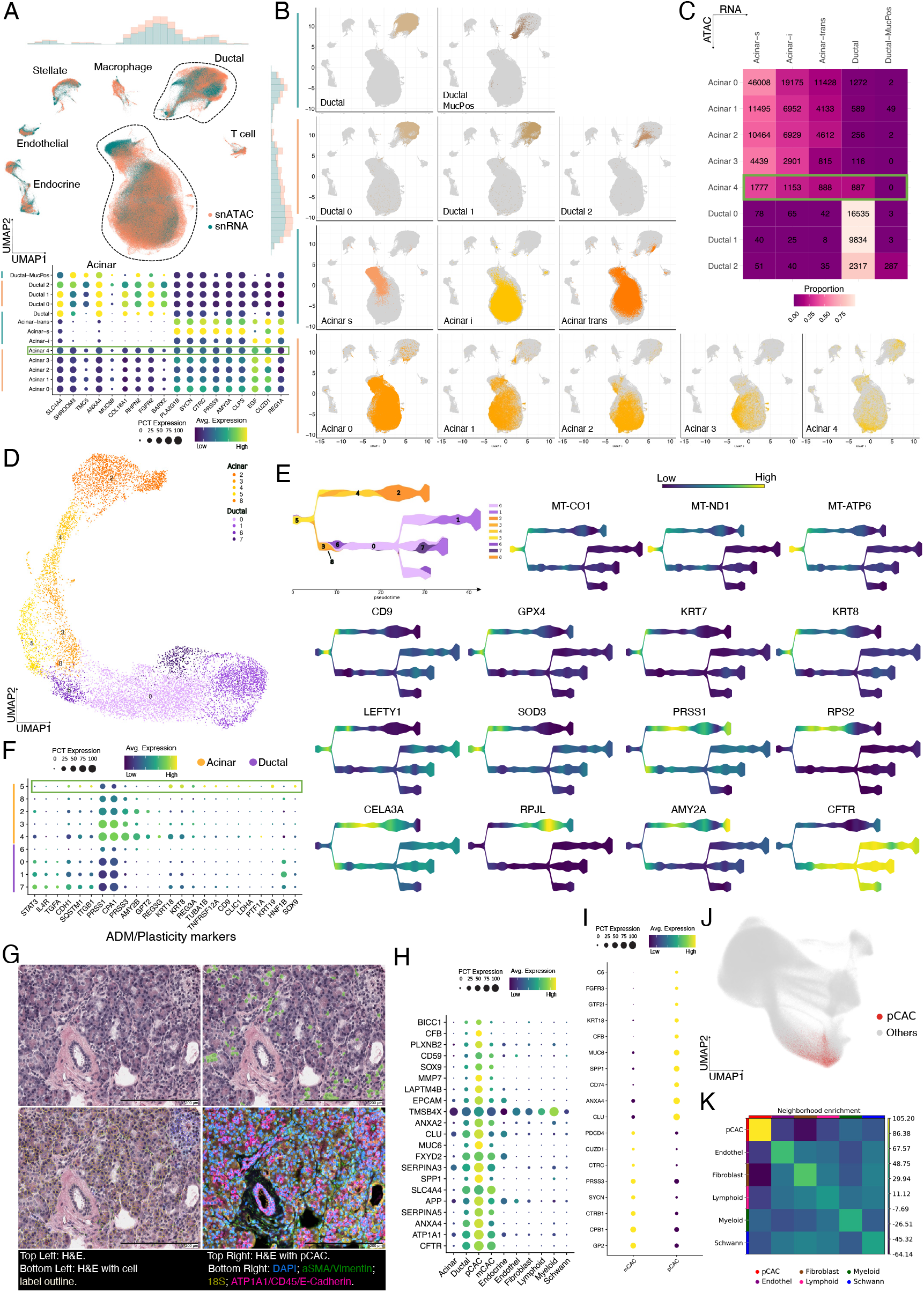
Transcriptional and Epigenetic Heterogeneity of the Human Adult Pancreas. **A**. UMAP visualization of 636,341 cells integrated from snRNA-seq and snATAC-seq datasets, using multiome data as a bridge. Points are colored by the originating technology. Marginal plots display the distribution of cells along the x- and y-axes for each technology. The dot plot highlights the expression of known acinar and ductal markers, based on deep annotation derived from the individual analysis of each dataset. **B**. UMAP showing the location of acinar and ductal subclusters identified by each technology. **C**. Heatmap displaying the results of transferring snRNA-derived labels (reference) to snATAC data (query) using DeepSCore, with colors representing cell proportions. **D**.UMAP reduction of 12,396 cells identified as acinar or ductal with highly plastic profiles in the snRNA data. Acinar cells exhibiting ductal-like gene expression and ductal cells showing acinar-like gene expression were merged and re-clustered for further analysis. **E**. Trajectory inference performed with STREAM for the 12,396 cells described in panel C. Highlighted are acinar-like and ductal-like lineages in orange and purple, respectively. Clusters shown in panel C are annotated, and gene expression patterns for acinar/ductal identity and plasticity markers are visualized. **F**. Heatmap of ADM/plasticity marker expression (curated from literature) across the clusters identified in panel C. The original cell type assignments for these clusters are also indicated. **G**. TME staining of healthy pancreas. The images were extracted from the same TME core under different staining conducted for the Xenium analysis. (Visualization generated with 10x Genomics Xenium Explorer 3.2.0). **H**. Expression levels of the pCAC signature genes in pCAC and broad cell types in the Xenium sample. **I**. pCAC signature expression level comparison between the cells identified as pCAC by the joint module score approach vs mCACs. **J**. UMAP reduction of the Xenium RNA assay. pCACs have been highlighted in red. **K**. Neighborhood enrichment analysis of the different cell types identified in the Xenium sample.

### High-resolution mapping of acinar and ductal cells uncovers rare plastic centroacinar cells in the adult human pancreas

Intrigued by the presence of hybrid acinar–ductal states, we focused our analysis on cluster 7 from the acinar population and a subset of cluster 4 from the ductal population in the snRNA-seq dataset (Supp. Fig. 2B–C), as these nuclei exhibited mixed transcriptional profiles suggestive of a transitional identity. Similar acinar-to-ductal plasticity has been described in disease contexts (e.g., acinar-to-ductal metaplasia, ADM), typically in mouse models (1, 20), but has not previously been characterized at this resolution in healthy human pancreas tissue. We therefore reanalyzed these nuclei together, revealing that, rather than forming two distinct groups, they converged into a shared root cluster (cluster 5) with a less differentiated phenotype (Fig. 2D). To further investigate this differentiation process, we performed pseudotime analysis using STREAM (21), enabling the reconstruction of complex branching trajectories from both single-nucleus transcriptomic and chromatin data (Supp. note 2 and 3). This analysis revealed that cells progressively diverged toward either a more acinar-like (clusters 2 and 4) or ductal-like identity (clusters 0, 1, 6, and 7), forming a continuum originating in cluster 5 (Fig. 2E). The resulting trajectories illustrated this branched transition, beginning with early activation of mitochondrial genes (22) (*MT-CO1, MT-ND1, MT-ATP6*); markers of cellular adaptation and plasticity such as *CD9* and *GPX4* (23, 24); metabolic regulators including *SOD3*; and cytoskeletal genes *KRT7* and *KRT8*, previously associated with ADM and pancreatic remodeling (22, 25, 26). Later stages were marked by expression of mature acinar (*PRSS1, AMY2A*) and ductal (*CFTR*) lineage genes.

Additionally, the co-expression of acinar and ductal markers in cluster 5, together with high levels of ADM- and plasticity-associated genes such as *KRT19, SOX9*, and *ITGB1* (27–29), further supports that cluster 5 represents a transcriptionally plastic cell population resembling centroacinar cells. *SOX9* has been described as being expressed in a subset of ductal epithelial and centroacinar cells in the human adult pancreas (30, 31), which we confirmed by performing immunohistochemistry staining on tissue cores from human adult pancreas. Our findings demonstrated that *SOX9* was expressed in both ductal and centroacinar cells, showing a positivity rate of 95%. This rate was determined by the presence of at least one positive cell of each cell type per core, based on a total of 77 tissue cores analyzed (Supp. Fig. 5A). Our STREAM analysis also identified *ANXA13* as a potential marker of this plastic population (Supp. Fig. 5B). This gene was previously reported of its higher expression in cholangiocellular carcinoma cells relative to pancreatic ductal adenocarcinoma cells (32). Notably, our results showed *ANXA13* expression in ductal and centroacinar cells within the screened tissue cores, with 91.2% positivity among a total of n = 80 (Supp. Fig. 5B). Given that *ANXA13* has not been previously associated with centroacinar cells, this highlights it as a potential novel marker of transitional ductal–acinar states in pCAC. We therefore refer to these cells as plastic centroacinar-like cells (pCACs). In contrast, *TGFA, IL4R*, and *STAT3* (Fig. 2F) are predominantly expressed in the opposite ductal branch, suggesting roles in immune modulation, while *REG3G* and *REG3A*, more enriched in acinar groups, are associated with regeneration and acinar identity. The transcription factor *HNF1B* also shows high expression in ductal cells, underscoring its relevance for ductal identity and function (Fig. 2F). Collectively, this expression landscape highlights the functional plasticity of transitional cells within this trajectory and their potential to adapt and shift phenotypic identity in the adult human pancreas.

### In situ validation of the plastic centroacinar-like cells (pCACs) using Xenium spatial transcriptomics

To further validate the existence and spatial localization of this rare, plastic subpopulation, we performed in situ transcriptomic profiling using high-plex spatial in-situ transcriptomics (Xenium) on pancreatic tissue (Fig. 1D, G). We selected a subset of samples from the same donors previously analyzed by snRNA-seq and snATAC-seq, specifically analyzing seven Tissue MicroArray (TMA) blocks from 14 individuals (Fig. 1B, Supp. Table 1). We utilized the pre-designed Xenium 5k panel (see Methods), supplemented with 100 custom probes (Supp. Table 3) targeting transcripts of particular interest, selected either as top differentially expressed genes in our single-cell analysis or as genes with relevant differentiation patterns that were absent from the core panel. These included 13 acinar-specific markers, 11 ductal markers, 26 genes associated with the pCAC population as well as genes for endocrine identity and other rare cell types.

To identify the pCAC within the spatial data, we applied a module scoring approach (see Methods) based on the pCAC gene signature derived from our snRNA-seq data (Supp. Table 4). The resulting in situ projection predominantly highlighted centroacinar cells, but also included a subset of acinar and ductal cells. We anticipated that technical differences between the snRNA-seq and the spatial transcriptomics platforms could contribute to discrepancies in cell identification. To refine this classification, we leveraged pathologist-annotated centroacinar cells across the TMA sections. Based on these expert annotations, we generated a second gene module score reflecting morphology-defined centroacinar cells (mCAC), which produced a highly specific spatial pattern consistent with their expected anatomical location and morphology. Yet, the snRNA-seq data indicated that the pCAC population does not represent the full spectrum of centroacinar cells, but rather a specific subset distinguished by elevated transcriptional plasticity. We therefore focused on the subset of centroacinar-like cells identified by both the single-nuclei pCAC signature and the pathologist-derived annotations. This overlap enabled us to derive a refined and robust in situ signature for those we identified as the pCACs, which comprised the 3.12% of all acinar and ductal cells in the adult snRNA dataset (Fig. 2G) vs the 1.43% found in the Xenium data. Like in snRNA-seq data, when compared to all other cell types in the spatial transcriptomics data using differential gene expression analysis (Supp. Table 5), pCACs showed high co-expression of markers such as *CFTR, SLC4A4, BICC1*, and *SOX9*, consistent with a ductal and centroacinar identity (Fig. 2H). In contrast, mCACs exhibited a more acinar-like transcriptional profile, with top differentially expressed genes including *CPB1, CTRB1, PRSS3, CTRC, SYCN*, and *CUZD1* (Fig. 2I). Notably, the pCACs also expressed a distinct set of genes associated with plasticity, stress response, and differentiation, such as *CLU, ANXA4, KRT18*, and *SPP1*, highlighting their transcriptional divergence and potential progenitor-like state.

Next, we systematically mapped the spatial distribution of pCACs (Fig. 2J) and interrogated their surrounding microenvironment. We performed neighborhood enrichment analysis (see Methods), which revealed that pCACs were consistently found in close proximity to myeloid cells (Fig. 2K), predominantly macrophages, beyond the expected proximity to acinar and ductal cells. This pattern was robust across all TMA cores and donors (Supp. Fig. 5C). While other stromal populations, such as lymphoid, fibroblasts, and endothelial cells, were occasionally nearby, their proximity varied depending on the specific core or sample. Interestingly, quantitative correlation analysis showed that the presence of myeloid cells had the strongest association with pCAC abundance (Pearson r > 0.8), whether measured across donors, across TMAs, or within individual cores.

### Transcriptional heterogeneity of adult pancreatic endocrine islet cells

The cellular diversity of pancreatic islets has been the subject of extensive study in recent years, revealing the existence of multiple, non-overlapping axes of transcriptional heterogeneity among islet endocrine cells (14, 33–39). This complex diversity in islet cell identity or state is thought to play a critical role in fine-tuning glucose homeostasis in health and disease (40). However, it remains uncertain which findings reflect biological differences or are instead influenced by sample processing or platform-specific artefacts.

To systematically explore the extent of this heterogeneity, we compared four complementary single-cell RNA-seq datasets that we generated using different technologies: 1) snRNA-seq of islet endocrine cells from frozen adult pancreas sections, 2) scRNA-seq of isolated cultured islets, 3) snRNA-seq of isolated frozen islets and 4) single-cell full-length mRNA VASA-seq of isolated cultured islets (Fig. 1D, 1F, 3A). This comparison across diverse approaches, using single nuclei and whole cells, employing 3’ biased and full-length mRNA sequencing, on isolated cultured islet cells or nuclei from frozen tissues that have not undergone pancreas digestion, purification, culture and dissociation processes, provides a multi-faceted view of transcriptional diversity and informs about potential artifacts and biases inherent to the different processing and sequencing methods (39, 41)(Supp. Fig. 7). Furthermore, we leveraged the spatial transcriptomics (Xenium) and proteomics (CODEX) datasets to orthogonally validate the existence of endocrine subpopulations.

**Figure 3.**
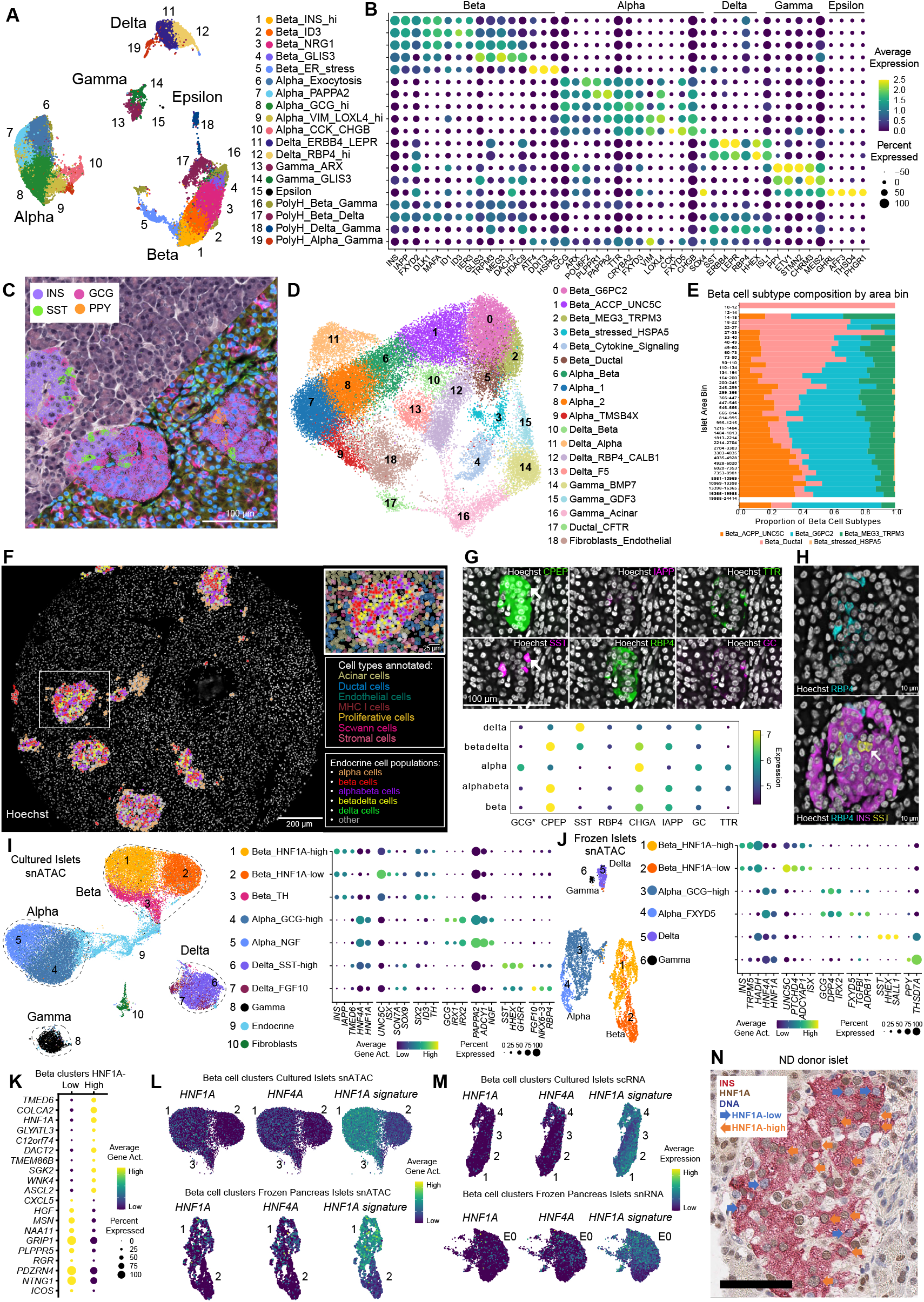
Heterogeneity of adult pancreas islet endocrine cells. **A**. UMAP projection of endocrine cells from scRNA-seq of isolated cultured islets (27,476 cells) colored by cell type subpopulations. **B**. Dotplot indicating the expression of characteristic cell type-subpopulation gene markers. **C**. Visualization of Xenium spatial transcriptomics data. A representative TMA core containing pancreatic islet structures, half-stained with hematoxilin-eosin and half with immunofluorescence markers (same markers as Fig. 1G). Transcript counts for the main endocrine hormones are overlaid on the composite image, as indicated in the legend on the top. (Visualization generated with 10x Genomics Xenium Explorer 3.2.0). **D**. UMAP projection of endocrine cells detected in the Xenium spatial transcriptomics data (40,714 endocrine cells) colored by cell type subpopulations. **E**. Distribution of beta cell subtypes detected in the Xenium data across endocrine clusters of different size. Endocrine clusters were binned by area in groups of 10 to 24,500 μm2 . **F**. Visualization of CODEX data focusing on islet cell populations. An illustration of the spatial compartmentalization of annotated islet cell populations within a single core of pancreatic tissue. The inset in the top right corner highlights a magnified view of a representative islet, showcasing all annotated cell populations. Label and protein stain colors are indicated in the images. **G**. Examples of identified endocrine cell populations with classical and hybrid states. Top row: Representative immunostaining images showing the expression of key protein markers: C-peptide (CPEP), transthyretin (TTR), amylin (IAPP), retinol-binding protein 4 (RBP4), and somatostatin (SST) in pancreatic tissue sections. Arrows indicate hybrid cell states co-expressing CPEP (beta cell–associated) and SST (delta cell–associated). Bottom row: A dot plot depicting protein expression levels of specific endocrine markers utilized for differentiating the main endocrine cell populations, including alpha, beta, delta, alphabeta, and betadelta cells. **H**. Tissue core from non-disease pancreas tail region showing heterogeneous expression of signature pancreatic islet protein markers. Arrow indicates an example of a SST immunoreactive positive cell that is RBP4 negative immunoreactive. C-PEP (magenta), RBP4 (cyan), SST (yellow). **I**. UMAP projection of endocrine cells in the cultured islets snATAC-seq dataset, colored by cell type subpopulations, and dotplot depicting the gene body chromatin accessibility (i.e. gene activity) of characteristic cell type-subpopulation markers. **J**. UMAP projection of endocrine cells in the endocrine subset of frozen adult pancreas snRNA-seq dataset, colored by cell type subpopulations, and dotplot depicting the gene activity of cell type-subpopulation markers. **K**. Dotplot depicting the gene activity of markers for beta cell subpopulations high and low in HNF1A activity. **L**. Feature plot representing the gene activity of HNF1A, HNF4A and HNF1A signature genes in cultured and frozen beta cell subpopulations from snATAC-seq datasets. **M**. Feature plot representing the gene expression of HNF1A, HNF4A and HNF1A signature genes in cultured and frozen beta cell subpopulations from sc/snRNA-seq datasets. **N**. Immunohistochemistry for HNF1A and INS protein in islets of an adult pancreas from a non-diabetes donor.

We first focused our analysis on the five major human islet endocrine cell types (alpha, beta, gamma, delta, and epsilon cells), that were consistently detected across all datasets. We also observed subpopulations of endocrine cells with a polyhormonal identity (expressing more than one canonical hormone of the major endocrine groups: INS, GCG, SST, PPY) (Supp. Fig. 8B, Supp. Table 6). To identify robust marker genes of the endocrine populations, we performed a comprehensive analysis throughout the different datasets (see Methods), comparing the genes differentially expressed across these cell types resulting in a consensus geneset panel for each population. (Supp. Fig. 8B, Supp. Table 7).

Notably, snRNA-seq of frozen sections exhibited a reduced cross-contamination of hormone transcripts, with less off-target detection of *INS, GCG, SST*. This suggests a diminished capture of leaked mRNA molecules, a common artifact in the human islet single cell RNA-seq, which was more pronounced in the other 3 datasets (Supp. Fig. 8B). In contrast, in the datasets that used isolated islets, we observed an increased expression of stress-response genes, which can reflect the impact of pancreas tissue dissociation, islet isolation and culture (e.g. intermediate early genes *FOS, FOSB, JUND;* ER-stress genes *ATF4, DDIT3, HSPA5*) (39) (Supp. Fig. 8B).

A detailed analysis of the isolated cultured islets scRNA-seq dataset (27,476 endocrine cells) unveiled various subpopulations of endocrine cell types that heterogeneously expressed genes associated with endocrine cell functions (Fig. 3A-B, Supp. Table 8). Beta cells presented five distinct subpopulations with an expression gradient of beta cell canonical markers, specifically *INS* and *IAPP* (Fig. 3B). The INS_high cluster 1 displayed higher expression of *INS* and mature beta cell marker *MAFA*, together with the highest expression levels of *IAPP, FXYD2, RBP4* and *CDKN1C*, indicating a beta cell population highly active in insulin biosynthesis (34, 39) (Fig. 3B). Cluster 2 presented increased expression of transcription factors *ID1* and *ID3*, a heterogeneity observation previously described in other studies (37). Cluster 3 presented an intermediate phenotype with intermediate levels of *NRG1* and *TRPM3*, a calcium channel involved in beta cell function (42). Cluster 4 displayed lower INS levels together with increased expression of GLIS3, an important transcription factor for beta cell development and function (43), *HDAC9* and *ZNF385D* (44). Finally, we identified a subcluster of stressed beta cells (cluster 5) expressing endoplasmic reticulum stress markers *ATF4, DDIT3*, and *HSPA5* (Fig. 3B, Supp. Table 8).

Alpha cells also presented heterogeneity in their gene expression across detected subpopulations. These included clusters with varying levels of *GCG*, and other canonical alpha cell markers like *TTR* and *CRYBA2* (Fig. 3B): cluster 6, with increased expression of genes involved in exocytosis, including *POU6F2, DCC*, and *NPAS3*; cluster 7 with elevated *PAPPA2, G6PC2, ASPH*, and genes involved in microtubule formation; cluster 8 with high levels of *LOXL4* and *VIM* and, a phenotype that has been associated with dysfunctional alpha cells (45), and cluster 9 with expression of *CCK* and *CHGB*. Furthermore, delta cells presented two clear subclusters, one marked with lower *SST* and increased *ERBB4* and *LEPR* expression (cluster 11), and one with increased *SST, HHEX, ISL1* and *RBP4* expression (cluster 12) (Supp. Table 8).

Next, we explored the presence of these endocrine cell subpopulations in the complementary islet datasets (Supp. Fig. 7), revealing distinct subpopulations for beta, alpha and delta cells, in all datasets, with clear heterogeneity in their gene expression patterns and exhibiting similar markers as those seen in cultured islets (Supp. Fig. 8C-F, Supp. Table 8).

The strategy used for pancreas frozen sections sampling enabled the investigation of regional differences in the composition of the endocrine cell compartment (Supp. Fig. 8G). Sections derived from the uncinate process and head of the pancreas exhibited an increased percentage of gamma cells (∼25%), consistent with previous histological observations (46). Additionally, our analyses revealed gene markers associated with specific pancreas regions for the major endocrine cell types (Supp. Fig. 8H).

### Spatial transcriptomic and proteomic investigation of adult pancreatic endocrine cells

We investigated islet endocrine cell heterogeneity in the spatial transcriptomics Xenium dataset (Fig. 3C), identifying a total of 40,714 endocrine cells, with distinct endocrine cell types and subpopulations (Fig. 3D, Supp. Table 6). For example, analysis of the spatial transcriptomic data identified different beta cell subpopulations with increased expression of *INS* and *ACPP* (cluster 1); genes involved in beta cell maturation and function: *G6PC2, MAFA, NTN1* (cluster 0); *MEG3, NTNG2, TRPM3* and MLXIPL (cluster 2). Also, this analysis pinpointed the presence of smaller beta cell subpopulations with elevated expression of endoplasmic reticulum (ER) stress markers *HSPA5* (cluster 3); cytokine signaling genes (cluster 4); and pancreatic ductal cell marker genes *CFTR* and *SERPINA3* (cluster 5).

We then explored the spatial distribution of these endocrine cell type subpopulations in the dataset. Unsupervised cell neighbourhood analysis was performed on the endocrine cells, identifying endocrine cell clusters that varied in size over a wide range (10 to 25,000 μm2 in area), from individual endocrine cells to well-defined islets (Fig. 3E, Supp. Table 9). Intriguingly, we identified numerous individual cells expressing endocrine hormone transcripts (*INS, GCG, SST; PPY, GHRL*) outside islets structures, scattered through the pancreas parenchyma and within ductal structures (Supp. Fig. 9). Furthermore, the distribution of the beta cell subpopulations across the endocrine clusters of different sizes was not homogeneous, with beta cell cluster 5 (Beta_Ductal) being more abundant in smaller endocrine clusters (10-500 μm2 in area), while beta cell clusters 0 and 1 mostly composing the bigger endocrine clusters/islets (2,000-20,000 μm2 in area). Thus, Xenium spatial transcriptomics enables the investigation of islet endocrine cell heterogeneity, spatial location and cell neighborhood composition, which is precluded with other higher-resolution single cell transcriptomic approaches where position information is lost due to tissue dissociation. Moreover, these results indicate the presence of endocrine cell populations of smaller size outside the well-defined and characterized islets structures, both scattered in the parenchyma and in ductal structures, but with a particularly distinct transcriptional profile (Supp. Table 9). These endocrine cell populations, which have been previously described using endocrine hormone immunostaining (47), have a hitherto unknown function in pancreas endocrine physiology. They could potentially represent a local source of hormone for pancreas parenchyma or specialized function to meet different metabolic demands (48), evidence of endocrine regenerative potential in the adult pancreas (49) or a subpopulation at higher risk of autoimmune recognition and destruction (50).

To gain deeper insight into the proteomic complexity of pancreatic endocrine cell populations, we applied highly multiplexed single-cell imaging to pancreatic tissue sections from the same donors (Fig. 3F). This enabled us to examine protein expression in endocrine cells across different histological regions while preserving spatial context. Following unbiased cell segmentation, quantification, and clustering (Supp. Fig. 10A-C), we annotated a total of 65,548 islet cells using an eight-marker antibody panel: CHGA, GCG, GC, and TTR (alpha cell-associated); CPEP and IAPP (beta cell-associated); and RBP4 and SST (delta cell-associated). This imaging-based approach provided a complementary view of islet cell heterogeneity and confirmed the presence of several transcriptionally defined subpopulations.

Notably, marker expression patterns across alpha, beta, and delta cells demonstrated considerable complexity, including cells with co-expression of multiple markers, potentially reflecting transitional or polyhormonal states (Fig. 3G). While classical beta cell population expresses both CPEP and IAPP markers (Suppl. Fig. 10B), other beta cell populations, which are still defined by CPEP positivity, exhibited heterogeneous expression of RBP4 (Suppl. Fig. 10C). This variation is consistent with previous studies showing that *RBP4*-negative beta cells display enhanced electrophysiological function compared to their RBP4-positive counterparts (39), suggesting that this proteomic variation may correspond to functional differences.

A further example of intrapopulation heterogeneity was observed in delta cells co-expressing CPEP, IAPP, and varying levels of SST (Fig. 3G, top row), mirroring transcriptional subtypes identified in the Xenium dataset. Another example consistent with transcriptomic findings, delta cells were found to be either RBP4-positive or RBP4-negative (Fig. 3H). These proteomic results reinforce the existence of transcriptionally defined endocrine subtypes and demonstrate that this cellular heterogeneity is preserved at the protein level and within the spatial architecture of the pancreas.

Proteomic heterogeneity and complex expression patterns were not limited to beta and delta cells but extended to every identified population, as demonstrated in Fig. 3G (bottom row), with markers such as TTR and RBP4 exhibiting lower overall intensities and GCG in annotated hybrid states excluded to avoid bleed-through artifacts. These findings reinforce the existence of intermediate endocrine cell states and suggest the presence of potentially transitional or hybrid polyhormonal phenotypes, consistent with observations made using multiplexed deep visual proteomics, which uncovered rare hybrid subpopulations with mixed alpha-, beta-, and delta-like signatures within native pancreatic islets (51). These hybrid phenotype cells were not randomly distributed; spatial analysis revealed that beta cells frequently neighbored other hybrid beta cells expressing markers of delta or alpha identity, suggesting possible spatially restricted microenvironments that support or reflect this phenotypic plasticity (Supp. Fig. 10D). In contrast, more canonical alpha and delta cell populations were spatially more distant from more mixed polyhormonal cell states (Supp. Fig. 10E).

Together, these findings illustrate the utility of integrating spatial transcriptomics and multiplexed imaging proteomics to comprehensively map the phenotypic diversity of islet endocrine cells. This dual-modality approach enables the identification and localization of rare or transitional endocrine states that may play critical roles in endocrine function, regeneration, or disease susceptibility within the native pancreatic tissue environment.

### Chromatin accessibility profiling of human islet endocrine cells reveals heterogeneous HNF1A activity within the beta cell population

We took advantage of snATAC-seq generated on the same isolated culture islet cells to investigate the level of heterogeneity in the endocrine cell populations at the chromatin accessibility level (Fig. 3I). We identified subpopulations of beta, alpha and delta cell types with distinct chromatin accessibility profiles, in line with the previous transcriptomic and proteomic observations (Supp. Table 10). Noteworthy, we also identified an additional level of heterogeneity in beta cells, related to HNF1A high and low transcription factor activity (Fig. 3I). Comparison of these two distinct beta cell clusters revealed different levels of chromatin accessibility at the gene body of transcription factor gene *HNF1A*, and its direct target genes (e.g. *TMED6, SGK2, A1CF, HNF4A*) (Fig. 3K, Supp. Table 10)(52).

Similar beta cell clusters with a *HNF1A* high and low chromatin accessibility program were also detected in the snATAC-seq from frozen pancreas sections, confirming the presence of these heterogeneity axis in non-cultured, minimally perturbed, adult human islet beta cells (Fig. 3J, Supp. Table 10). We selected the top 25 genes with differential accessibility (i.e. gene activity) between these two beta cell populations in cultured and frozen beta cells (Fig. 3K, Supp. Table 10) and utilized them to investigate in the snATAC and scRNA datasets the presence of cells with enhanced *HNF1A* transcriptional program (Fig. L-M). Furthermore, we conducted immunohistochemistry for HNF1A and INS on adult pancreas sections and observed that ∼30% of INS+ beta cells presented lower levels of HNF1A protein in their nuclei, validating the heterogeneous expression of HNF1A in human beta cells also at the protein level (Fig. 3N). Thus, we validated the presence of subpopulations of beta cells with increased and decreased *HNF1A* gene signature at the protein, transcript and transcription factor activity level, indicating that HNF1A-high and low programs in beta cells represent an additional axis of beta cell heterogeneity (53, 54).

### Regulatory landscape of islets from individuals with type 2 diabetes

Utilising single-cell genomics to investigate human islets in type 2 diabetes (T2D) offers a transformative platform for unraveling the molecular mechanisms underlying this intricate metabolic disorder. Here, we investigated the transcriptional and chromatin accessibility changes between non diabetic (ND) and T2D exhibited by human islets obtained from donors combining scRNA-seq and snATAC-seq data. Additionally, we explored the impact of culturing in different glucose concentrations on the regulatory dynamics of endocrine cells and compared these effects with alterations induced by long-term in vivo hyperglycemia characteristic of T2D.

Initially, we integrated scRNA-seq data from isolated cultured islets obtained from both ND and T2D (Fig. 4A, Supp. Fig. 11A-B). Leveraging scOMM to compare the heterogeneity in endocrine cell populations observed in the ND dataset to that of the T2D dataset, we identified similar populations in both datasets (Supp. Fig. 11C). Subsequently, we performed differential cell type abundance analysis in the integrated dataset using MILO (55), which assigns cells to partially overlapping neighborhoods on a k-nearest neighbor graph. This unveiled various cell neighborhoods particularly enriched in ND and T2D (Fig. 4B, Supp. Fig. 11D).

Beta cells in ND neighborhoods presented increased expression of *IAPP, RBP4, PCSK1* and *FXYD2*, together with ribosomal protein genes *RPS26, RPS29*, suggesting a cell subpopulation with enhanced insulin biosynthesis (56). T2D beta cell neighborhoods presented increased expression of *SPP1, DGKB* and *TFF3*, genes previously linked with beta cell dysfunction and development (57– 59) (Fig. 4C, Supp. Table 11). Alpha cells in ND neighborhoods showed higher expression of mitochondrial genes involved in oxidative phosphorylation (*MT-ND3, MT-CYB, MT-CO1*), while the T2D alpha cells presented increased expression of *IGFBP2, B2M* and *CD44* (Fig. 4C).

To further delineate genes that are consistently dysregulated in beta cells from T2D individuals, we compared the differentially expressed genes across three different datasets: 1) cRNA-seq of isolated cultured islets (5 ND and 6 T2D donors); 2) snRNA-seq of isolated frozen islets (4 ND and 4 T2D donors); and 3) VASA-seq of isolated cultured islets (6 ND and 5 T2D donors). Despite major differences in tissue processing and sequencing technologies, we identified 87 genes that were downregulated on the T2D beta cells common to the 3 datasets, including transcription factors (*SOX6, ELF1, ELF3* and *HIF1A*), direct HNF1A targets (*A1CF, HNF4A, TMED6*), regulators of signaling pathways (*MAP3K8, RICTOR*) and cilia-related genes (*CFAP70, DNAI1*) (Fig. 4D).

In the intersection of T2D beta cells upregulated genes (64 genes), we found transcription factors related to beta cell maturation (*MAFA, SIX3* and *ONECUT2*) (59), suggesting a compensatory response to improve beta cell function. Also, we observed an upregulation of genes associated with islet immune response (*GAD1, GAD2, HLA-A*), netrin receptors (*UNC5C, UNC5B, DSCAM*), previously linked to beta cell survival and insulin secretion (60), and signaling pathways (*P2RY1, AKT2, PLK2, WNT4, PRICKLE2, CDKN1A, GADD45A*) (Fig. 4F, Supp. Table 12). The overlap at the level of biological pathways in which these genes are involved was considerable: cilium and microtubule-based movement, membrane trafficking, regulation of organelle organization, cytoskeleton, protein localization or cell projection were among the most enriched biological pathways in the genes that are downregulated in T2D beta cells across datasets. Conversely, regulation of growth, response to nutrients and stress, signaling by GTPases and SRP-dependent cotranslational protein targeting to membranes were the biological pathways enriched in T2D beta cell upregulated genes. These parallel analyses highlight the common dysregulated pathways across datasets.

**Figure 4.**
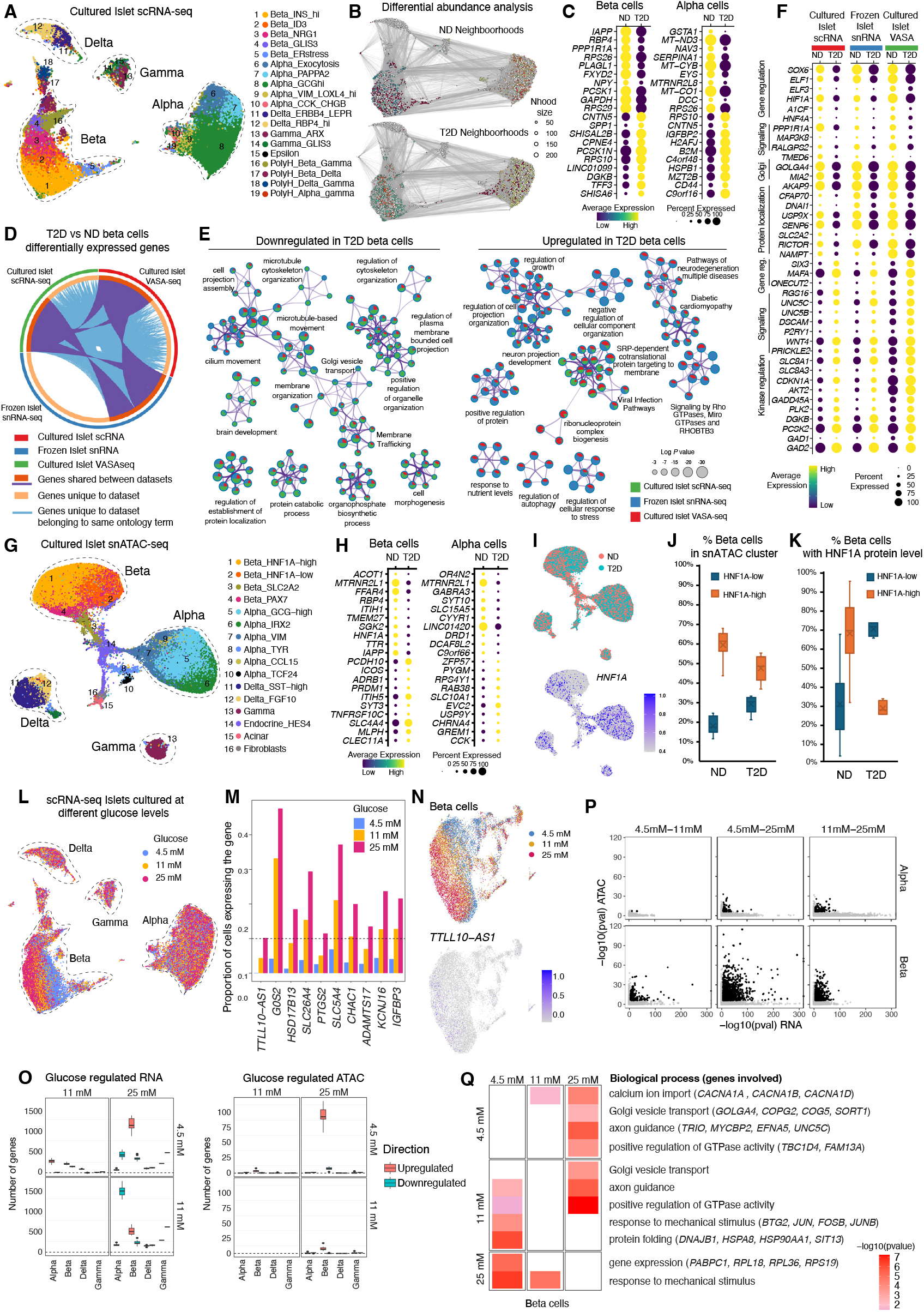
Regulatory landscape of human islets from individuals with type 2 diabetes. **A**. UMAP projection of endocrine cells from cultured islets scRNA-seq integrating T2D and ND donors colored by cell type subpopulations. **B**. Differential cell abundance analysis of integrated T2D and ND scRNA-seq datasets, depicting neighbourhoods specific for T2D and ND. **C**. Dotplot representing the expression of genes differentially regulated in beta and alpha from ND and T2D donors. **D**. Differentially expressed genes in T2D beta cells that are common across the three datasets, with genes shared between datasets labelled in orange and purple curves, genes unique for a dataset labelled in beige and genes sharing ontology term enrichment labelled in blue curves. **E**. Metascape networks of enriched ontology terms in T2D vs ND beta cells downregulated and upregulated gene sets from the three different datasets. **F**. Dotplot representing the expression of T2D beta cells differentially regulated genes that are common to the three datasets. **G**. UMAP projection of endocrine cells from cultured islets snATAC-seq integrating T2D and ND donors colored by cell type subpopulations. **H**. Dotplot representing the expression of gene bodies that present differential chromatin accessibility in beta and alpha cells from ND and T2D donors. **I**. UMAP projection of endocrine cells from cultured islets snATAC-seq integrating T2D and ND donors colored by disease status (ND or T2D, top) and HNF1A gene body chromatin accessibility level (bottom). **J**. Proportion of beta cells in HNF1A-high or HNF1A-low snATAC-seq cluster in samples from ND and T2D donors. **K**. Proportion of beta cells (INS+) presenting high or low levels of HNF1A protein immunoreactivity in adult pancreas sections from ND and T2D donors. **L**. UMAP projection of endocrine cells from islets cultured at different levels of glucose. **M**.Proportion of beta cells expressing glucose-induced genes at different glucose concentrations. **N**. UMAP projection of beta cells clustering coloured by culture glucose concentration. Expression of glucose-induced gene TTLL10-AS1 in cultured beta cells. **O**. Number of glucose regulated genes in different islet cell types and glucose concentrations at RNA and ATAC level. **P**. Significance relationship between glucose-regulated genes at RNA and ATAC level in alpha and beta cells. **Q**. Biological processes enriched among the gene sets regulated by glucose in beta cells.

We also explored the scATAC dataset of T2D and ND human islets and observed a reduced proportion of beta cells presenting the HNF1A high program in T2D donors, aligning with the downregulation of HNF1A target observed in T2D scRNA-seq datasets (Fig. 4G-K).

### Islets cultured in different levels of glucose reveal cell-type specific transcriptional programs in non-diabetic donors

Human islet cells are exposed to a wide range of glucose concentrations in health and disease. This creates a need to understand single cell expression at different glucose states. We investigated the response of endocrine cells to different glucose concentrations using scRNA-seq and snATAC-seq on islets from non-diabetic donors cultured for at least 72 hours in 4.5 mM, 11 mM and 25 mM glucose, corresponding to fasting, mildly elevated glycemia, and severe hyperglycemic levels, respectively. We functionally characterized them using dynamic glucose stimulated insulin secretion in a perifusion assay, observing that culture in 11 or 25 mM glucose significantly reduced insulin secretion capacity in a glucose-concentration dependent manner (Suppl. Fig. 12A-F). Additionally, total insulin content was unchanged, indicating that insulin production is conserved between preparations and culture conditions (Suppl. Fig. 12G).

Culture of pancreatic islets in different glucose concentrations induced condition-specific transcriptional profiles (Fig. 4L-N). We investigated cell type-specific responses to glucose variation, observing different patterns of glucose response between cell types and modalities (Fig. 4O). Alpha cells and beta cells showed increased transcriptional responses to glucose variation than delta cells and gamma cells. Using snATAC-seq, we similarly observed an increased gene activity response in beta cells compared to other cell types (Fig. 4O). Altogether, this suggests cell type-specific patterns of responses to increasing glucose concentrations.

In alpha cells, we observed a majority of downregulated genes induced by 25 mM glucose compared to 4.5 mM or 11 mM (2,837 genes). Interestingly, an opposite phenomenon was observed in beta cells, where a majority of genes induced by 25 mM of glucose are upregulated compared to 4.5 mM or 11 mM (4,272) genes (Supp. Table 13). Similar to the scRNA-seq dataset, most of the gene activity variation in beta cells occurred relatively to 25 mM of glucose (Fig. 4O). The biological relevance of identified RNA glucose-dependent genes was further explored in each cell type and pairwise comparison (Supp. Table 13, Supp. Fig. 12H), where an enrichment of genes related to glucose homeostasis, insulin secretion and stress response was identified.

We then explored the similarities between scRNA-seq and snATAC-seq signals focusing on alpha cells and beta cells, which have a higher number of genes identified as differentially expressed or active. We first investigated the significance relationship between the two modalities (Fig. 4P). We observed a partial recapitulation of scRNA-seq signal in beta cells in snATAC-seq, with an overlap of 30.5% of genes below p-value threshold. Next, we investigated whether these common genes showed the same pattern of log2FC between scRNA-seq and snATAC-seq in beta cells (Supp. Fig. 12I). All pairwise tests together, 92% of genes shared the same direction of log2FC between the two modalities, indicating consistent glucose-dependent signal in between scRNA-seq and snATAC-seq. We used this robust set of gene to perform gene set enrichment analysis, and identified key pathways enriched, such as calcium ion import (GO:0070509) at relatively higher concentration of glucose (11 mM vs 4.5 mM, 25 mM vs 4.5 mM), Golgi vesicle transport (GO:0048193) induced at 25 mM of glucose, or response to mechanical stimulus (GO:0009612) induced at 4.5 mM of glucose (Fig 4Q)(see also Supp. Note 4).

### Endocrine lineage specification and NEUROG3 dynamics in fetal pancreas

Human pancreas development remains poorly understood due to limited access to fetal tissue, but single-cell and spatial omics technologies are beginning to overcome this barrier. Here, we performed snRNA-seq (frozen material) and high-plex spatial proteomic imaging (Formalin-Fixed Paraffin-Embedded - FFPE sections) from dissected fetal pancreases, ranging from 10 to 18 weeks post conception (wpc) (Fig. 5, Supp. Fig. 13-16, Supp. Table 14).

After ambient RNA correction and stringent quality control (QC) (see Methods), this human fetal pancreas snRNA-seq dataset comprised 87,117 high quality nuclei and consist of 44 transcriptionally distinct annotated populations, including acinar, ductal, endocrine, mesenchymal, endothelial, neuronal, and immune cells. Rare and transitional populations, such as pre-acinar and *NEUROG3+* endocrine progenitors, were identified alongside canonical alpha, beta, delta, and epsilon endocrine subsets (Fig. 1E, Supp. Fig. 13B-D, Supp. Fig. 17)(see also Supp. Note 5). Complementary spatial proteomics from 224,713 FFPE-embedded cells confirmed the presence of these populations across developmental stages (Fig. 5A-B, Supp. Fig. 13-16) (see also Supp. Note 6).

**Figure 5.**
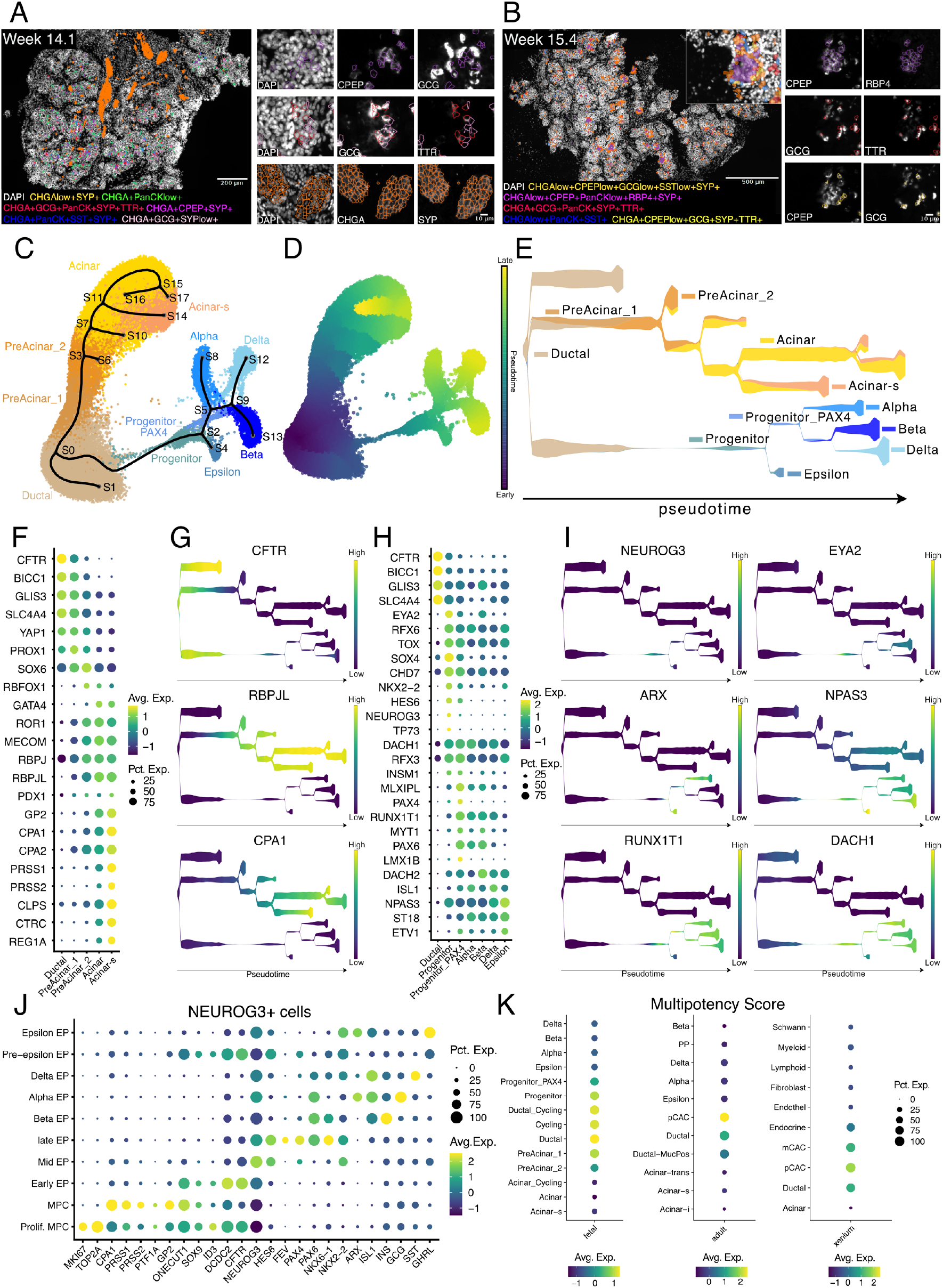
Transcriptional heterogeneity of human fetal pancreas. **A-B**. Multiplexed imaging of endocrine populations during human pancreas development at 14.1 (A) and 15.4 (B) weeks post conception (wpc), accompanied by close-up images. **C-E**. STREAM analysis of the fetal epithelial subset without cycling cells. C. UMAP plot of epithelial subset showing the trajectory inference (branches) calculated by STREAM. **D**. Visualization of thepseudotime on the same UMAP. **E**. Stream plot visualizing cell density along different trajectories over pseudotime. Plot is colored by cell types. **F**. DotPlot showing the expression of genes/transcription factors in the fetal dataset from the ductal to the acinar compartment. **G**. Stream plots visualizing CFTR, RBPJL and CPA1 expression over pseudotime for exocrine differentiation identified by STREAM analysis. **H**. DotPlot showing the expression of genes/ transcription factors in the fetal dataset from the ductal to the endocrine compartment. **I**. Stream plots visualizing NEUROG3 expression over pseudotime as well as other significant transcription factors for endocrine differentiation identified by STREAM analysis. **J**. Dotplot of NEUROG3 expressing cells showing different markers of multipotency and cell identity. **K**. Dotplot showing the multipotency module score across the fetal, adult and xenium datasets.

### Developmental trajectories of human fetal pancreas

To infer the developmental trajectories in fetal pancreas, we applied the pseudotime analysis tool STREAM to our dataset (21), identifying and reconstructing three main branching cellular trajectories (Fig. 5C - E): i) A short ductal differentiation trajectory, likely representing the development of “trunk progenitors” into more mature ductal cell identity; ii) A ductal to acinar differentiation trajectory, characterized by cells expressing both ductal (*CFTR, BICC1, GLIS3*) and acinar markers (*RBPJ, RBPJL, MECOM*) at the beginning of the pseudotime, with diminishing ductal marker expression and increasing acinar marker expression along the trajectory (Fig. 5F-G). iii) An endocrine differentiation trajectory, starting at the transition from ductal cells to endocrine cells with a cell population expressing *NEUROG3* (Fig 5H-I, Supp. Fig. 17), a key transcription factor for the initiation of endocrine development (60, 61). The next population of endocrine progenitors in the trajectory expressed known transcription factors important for endocrine cell formation and lineage allocation, such as *RFX6, PAX4, ARX* and *RUNX1T1* (62–64). Pseudotime analysis pinpointed other transcription factors that could potentially play a role in endocrine cell development, with unknown function in this context (*EYA2, NPAS3, DACH1, DACH2*) (Fig. 5I, Supp. Fig. 13E). The endocrine progenitor trajectory branches into developing epsilon cells (*GHRL, GHRLOS*) and a branch transcriptionally similar to endocrine progenitors, but with reduced *NEUROG3* and increased *PAX4* expression (‘Progenitor_PAX4’ cluster). This branch bifurcated first into the alpha cell cluster (*GCG, TTR*), which also has some *PPY*-expressing cells (likely some early gamma cells) (Supp. Fig. 13D), and then into beta (*INS, MAFA*) and delta cell branches (*SST, RBP4*), reconstructing the trajectories for all the pancreatic islet endocrine cell types.

The expression of *NEUROG3* during fetal pancreas development is critical for the differentiation of endocrine cells (60, 65). We observed the highest *NEUROG3* expression in the arising endocrine cells but also at a lower proportion in the clusters annotated as acinar and ductal cells. Cells expressing *NEUROG3* were selected and reclustered, identifying 10 different subpopulations (Fig. 5J, Supp. Fig. 17A-E). Previous studies have identified distinct subclusters of endocrine progenitor (EP) cells with a varying expression of *NEUROG3* (66, 67). We identified Early EP cells with expression of ductal markers *DCDC2, CTFR, SOX9, ID3* and high expression of *ONECUT1*, Mid EP cells which presented the highest expression of *NEUROG3*, and Late EP cells, that expressed *FEV, PAX4, PAX6, NKX2-2* and *NKX6-1*. Early, mid, and late EP cells expressed reduced levels of endocrine hormone genes. Within the *NEUROG3*-expressing subset, we also identified distinct clusters with clear endocrine cell type expression profiles. Pre-epsilon and Epsilon EP cells expressed *GHRL* and *NKX2-2* and were transcriptionally close to Mid EP, while Beta EP (*INS+*). On the other hand Alpha EP (*GCG+*), and Delta EP (*SST+*) cells were closer to Late EP cells (Supp. Fig. 17C-E), indicating their developmental lineage. Multipotent progenitor cells (MPCs) expressed markers such as *GP2, PTF1A, ONECUT1*, and acinar-specific genes *CPA1, PRSS1*, and *PRRS2*. MPCs have been detected in the developing human fetal pancreas expressing *CPA1* and *GP2* (8, 68), however, the presence of *NEUROG3* in these cells has not been previously reported. A subset of MPCs presented high expression of cell cycle marker genes *MKI67* and *TOP2A*, indicating their active proliferation status. Additional cell cycle analysis further highlighted the proliferative nature of these MPCs, with decreasing S and G2M phase gene signature towards the more differentiated EP cell types (Supp. Fig. 17D). *NEUROG3* expression has been suggested as a sign of cell cycle exit and unipotent endocrine precursor cells (69). However, we observed a lower but detectable expression of *NEUROG3* and *PTF1A* in multipotent and proliferative cells, when compared to fetal acinar cells and Mid EP cells (Supp. Fig. 17E). These results highlight the presence of *NEUROG3*-low MPC during the endocrine cell expansion/ development. We examined the expression of genes associated with the Notch signaling pathway, important for pancreatic and endocrine development (70, 71) (Supp. Fig. 17F). We observed high expression of *DLL1, YAP1, HES1*, and JAG1 in MPCs and early EP, with a consistent decrease in mid EP and the specified endocrine cell EP populations. This decrease coincides with the increased expression of *NEUROG3* as previously reported (72). Furthermore, the pre-epsilon cells showed a higher expression of NOTCH signaling markers, suggesting their earlier endocrine progenitor state. Finally, we observed preferential expression of MPCs markers in ductal and pCAC populations in adult pancreas snRNA-seq and Xenium datasets (Supp. Fig. 17G-I).

### Comparison of transcriptional programs in adult and fetal pancreatic cells

This ESPACE project enables the rare opportunity to comprehensively compare fetal and adult pancreas snRNA-seq datasets generated with the same method and in the same lab, minimizing technical batch effects. We subset both datasets into the three main compartments of the pancreas (acinar, ductal and endocrine), integrated them and calculated differentially expressed and conserved genes between adult and fetal datasets (see Methods) (Supp Fig. 18, Supp. Table 15).

In the endocrine subset fetal alpha, delta and epsilon cells form trajectories extending toward their adult counterparts, rather than merging into fully integrated clusters, suggesting ongoing development (Supp. Fig. 18A). In contrast, fetal and adult beta cells form distinctly separated clusters (Supp. Fig. 18A&L), indicating a more pronounced transcriptional divergence. Although postnatal beta cell maturation is well established (73, 74) and supported by recent comparative transcriptomic studies (75), our data resolve this transition with higher resolution at the single-cell level, revealing distinct clustering and transcriptional programs. While UMAP visualizations are qualitative, the observed structure is supported by differential gene expression and gene ontology analyses. INS is the most conserved gene between fetal and adult beta cells, whereas other shared genes such as *PLAGL1* and *ROBO2* are involved in broader cellular functions like proliferation and migration (Supp. Fig. 18B). GO analysis of the top 100 differentially expressed genes highlights the functional divergence: fetal beta cells are enriched for developmental terms, while adult beta cells show enrichment in metabolic and stimulus response pathways (Supp. Fig. 19D). These patterns are reflected in compartment-specific markers, such as *SYT16* in adult cells, linked to vesicle trafficking (76), and the transcription factor *PLAGL1* in fetal cells (66) (Supp. Fig. 18B). In general, the fetal endocrine population shows elevated expression of genes typically associated with neuronal development (Supp. Table 15), consistent with developmental parallels between endocrine and neural lineages. (For further GO comparisons see Supp. Fig. 19A-D.)

Integration of the ductal compartments revealed comparable patterns: despite both fetal and adult ductal cells expressing canonical markers (*CFTR, SLC4A4*), they form distinct clusters (Supp. Fig. 18C). GO analysis of the fetal population highlighted terms associated with axon guidance, morphogenesis, and projection development—primarily driven by genes such as *SLIT2, ROBO1/2, EPHA7*, and *SEMA3C*. This finding is supported by previous work showing that the SLIT–ROBO signaling axis regulates epithelial identity and branching morphogenesis in the developing pancreas (77). *ROBO1/2* are expressed in early pancreatic progenitors and are required for their maintenance and expansion, while SLIT ligands are expressed more in the surrounding mesenchyme (77). In contrast, adult ductal cells show enrichment in secretion and signal transduction pathways, reflecting their mature functional role (Supp. Fig. 19E).

Fetal and adult acinar cells displayed marked transcriptional differences, even between mature acinar cells within the fetal subset and those in the adult pancreas (Supp. Fig. 18E&F). Differential expression analysis revealed that adult acinar cells are enriched for digestive enzyme transcripts such as *PNLIP* and *AMY2A*, whereas fetal acinar cells show higher expression of lncRNAs (*MEG3, MEG8)* and genes involved in tissue remodeling (*ELMO1, HPSE2*). The most differentially expressed gene, *DLK1*, is a component of the Notch signaling pathway, which plays a critical role in exocrine pancreatic development (78). Additionally, *DLK1*, along with *MEG3* and *MEG8*, is part of the imprinted *DLK1-DIO3* locus, which is active during development (79). Conserved markers between fetal and adult acinar cells include both general acinar identity genes (*GP2, RBPJL*) and early digestive enzymes (*CEL, CELA3A*). Interestingly, the dominant adult enzymes (*PNLIP, AMY2A*) show minimal to no expression in the fetal subset, consistent with previous findings (13, 80) (Supp. Fig. 18F). GO term enrichment based on the top 100 DEGs further supports these observations: adult cells are enriched for terms related to pancreatic secretion, digestion, and protein regulation, while fetal cells show enrichment for terms related to growth, tissue organization, and only later for secretion, which is consistent with the fact that nutrient absorption by the gut becomes relevant only after birth (Supp. Fig. 19F).

To connect these developmental insights with our findings in the adult pancreas, we revisited the plastic centroacinar cells (pCACs) that we previously identified in the adult dataset (Fig. 2G-K). Based on the integration of fetal and adult pancreas, we observed that pCACs share transcriptional features with fetal acinar-precursor populations, particularly those marked as PreAcinar_1, suggesting a conserved or reactivated gene program. To investigate this further, we curated a set of multipotency-associated genes from published fetal pancreas studies (81–83) and computed a module score across all annotated cell types (see Methods). Among adult cells, pCACs displayed the highest multipotency scores, with expression levels comparable to the fetal progenitor-like populations (Fig. 5K). To validate this observation in an independent spatial dataset, we applied the same module score to the Xenium adult pancreas data and again found that pCACs had the highest multipotency score among all cell types (Fig. 5K), reinforcing their unique transcriptional identity and suggesting persistence of developmental programs.

## Discussion

The human pancreas is a highly heterogeneous and functionally complex organ, and its plasticity is both a critical feature of homeostatic maintenance and a contributor to disease vulnerability. In this work, we present the ESPACE atlas, a multimodal, high-resolution reference encompassing over four million cells and nuclei from 57 individuals spanning fetal development, adult homeostasis, and T2D. A key strength of ESPACE lies in its integration of diverse single-cell technologies. By integrating single-cell transcriptomics, chromatin accessibility, spatial transcriptomics, and highly multiplexed proteomics, we offer a unified framework for understanding pancreas biology in unprecedented detail.

We leveraged this unique dataset to uncover rare but transcriptionally distinct plastic centroacinar-like cells (pCACs), which occupy an intermediate state along the acinar-to-ductal axis. Identified in snRNA-seq and snATAC-seq, these cells co-express acinar and ductal markers and display elevated expression of stress- and plasticity-associated genes. Spatial transcriptomics revealed that these cells are consistently located near myeloid, and also lymphoid, immune cells, suggesting an association with reactive or inflammatory environments. Proteomic validation confirmed their centroacinar identity via SOX9 and identified ANXA13 as a novel marker.

Notably, integration with fetal pancreas data revealed that pCACs share transcriptional features with fetal acinar progenitors, suggesting that these adult cells retain or reactivate a developmental gene program associated with cellular plasticity. Altogether, these findings suggest that the adult human pancreas retains a transcriptionally plastic niche potentially primed for regeneration or metaplasia, an observation with significant implications for pancreatic cancer initiation and repair processes.

In the endocrine compartment, multimodal profiling revealed substantial heterogeneity across all five major islet cell types. Notably, beta cells exhibited epigenetic divergence at the *HNF1A* locus, giving rise to *HNF1A*-high and HNF1A-low chromatin accessibility states, a novel axis of regulatory heterogeneity that was validated across technologies and sample sources. Proteomic staining analyses further confirmed subtype-specific marker expression at the protein level.

The vast majority of pancreatic endocrine cell omics study isolated islets or sorted cells, thus introducing at least 2 potential confounders, the stress applied to isolate and culture the islets and the inability to analyse endocrine cells outside well-developed islets. Our spatial transcriptomic analysis addressed this limitation and uncovered endocrine-like cells situated outside conventional islet structures, often in close proximity to pancreatic ducts. These cells expressed a unique set of genes associated with progenitor-like or plastic states (e.g., *CD44, CD24, CD9, CXCL12, H19, TGFBR2*), suggesting potential roles in regeneration, lineage flexibility, or stress response (84). Interestingly, the proximity of these cells to ductal regions aligns with prior evidence implicating the ductal compartment as a reservoir of facultative progenitors in the pancreas (31, 85). While functional validation is needed, their transcriptomic signature hints at a transitional identity distinct from both classical endocrine and exocrine lineages.

Our profiling of fetal pancreas provided a high-resolution view of human pancreatic development. Pseudotime analyses reconstructed the three major lineage trajectories, acinar, ductal, and endocrine, and identified transcription factors, such as *EYA2, NPAS3, DACH1, DACH2*, which are not yet described in the context of endocrine lineage commitment. The fine-grained dissection of *NEUROG3+* cell states revealed intermediate populations poised for commitment to all endocrine lineages and expands our understanding of human islet ontogeny. Comparison to adult data revealed that fetal and adult cell types, while sharing classical cell type identity markers, remain transcriptionally distinct. In the endocrine compartment, alpha, delta, and epsilon cells exhibit continuum-like transitions toward their mature states, whereas fetal and adult beta cells remain transcriptionally separated, highlighting the postnatal maturation of beta cells.

From a disease perspective, our atlas provides new insights into T2D-associated alterations. In T2D donors, we observed changes in beta cell subpopulation abundance, with a notable reduction in the *HNF1A*-high epigenetic state. Transcriptionally, T2D islets displayed both compensatory (e.g., increased *MAFA, SIX3, ONECUT2*) and dysfunctional (e.g., downregulated HNF1A targets) programs, suggesting concurrent adaptation and failure. Exposure of ND islets to varying glucose concentrations recapitulated some of these patterns and revealed cell-type-specific transcriptional and epigenetic responses, particularly pronounced in beta cells. These data underscore the dynamic regulatory landscape of islet cells and suggest that metabolic context can influence cell identity and function in a manner akin to chronic disease. In conclusion, the ESPACE Human Pancreas Atlas represents a foundational resource for the biomedical community. It enables comprehensive exploration of pancreas biology across time, space, and disease. As we have demonstrated, by bridging molecular, spatial, and developmental layers, this atlas provides the tools needed to interrogate fundamental questions in pancreatic function, regeneration, and pathology. Importantly, the integrative framework we established here can serve as a blueprint for future organ atlases, helping to illuminate human biology with single-cell precision.

### Study Limitations

While the ESPACE Human Pancreas Atlas provides a comprehensive and multimodal characterization of pancreatic cell states across development, homeostasis, and disease, several limitations should be acknowledged. First, although we included samples from 57 individuals and spanned diverse anatomical regions and disease contexts, the availability of matched data across all modalities (e.g., spatial, transcriptomic, epigenetic) for every donor was limited, potentially constraining some integrative analyses. Fetal tissues from different time points were divided for processing in transcriptomics and proteomics analyses, which may influence the output results of each technology (86). Second, we demonstrate that enzymatic dissociation and in vitro culturing of islets, although controlled, introduce stress-related transcriptional artifacts, particularly in endocrine cells. While we addressed this by including frozen nuclei-based assays, certain dynamic cellular states may still be under- or over-represented. Third, snATAC-seq remains lower in resolution than RNA-based modalities, limiting fine-grained identification of rare regulatory and transitional states, especially important during plastic transformation. Additionally, as chromatin profiling was not performed using multimodal (RNA+ATAC) assays, clear matching between transcriptional and epigenetic states in both exocrine and endocrine populations remains limited, and inferred correspondences rely on cross-modal label transfer and deep-learning–based integration.

While single-cell and single-nucleus RNA sequencing are powerful tools for profiling complex tissues, they are subject to technical limitations. In our fetal pancreas datasets, we observed a marked underrepresentation of mesenchymal cells in snRNA-seq compared to spatial proteomics (CODEX). This discrepancy likely arises from cell-type-specific sensitivity to nuclear isolation protocols, as mesenchymal and stromal populations are particularly vulnerable to mechanical and chemical stress during tissue processing, leading to their selective loss. Comparable biases in cell type recovery have been documented in studies comparing dissociative methods and spatial approaches (87, 88). This example illustrates the importance of multiomics integration in enabling a holistic approach; in our study, we combined sequencing technologies with spatial proteomics to achieve better sample representation.

Spatial transcriptomics, while preserving native tissue architecture and avoiding dissociation bias, also faces important technical challenges. One of the most critical is accurate cell segmentation, especially when attempting cell-level comparisons with sc/snRNA-seq data. Despite recent advances, current segmentation algorithms, even those that incorporate protein-based markers such as in Xenium Prime, remain imperfect. As a result, transcript-to-cell assignments may be inaccurate, and these misassignments can distort cell identity, introduce artifacts, and hinder downstream analysis and cross-platform integration. This challenge is amplified by the inherently sparse nature of transcriptomic data in spatial platforms compared to single-cell RNA-seq. While segmentation-free methods, such as pixel- or spot-based analyses (89, 90), can reduce segmentation bias and capture continuous spatial gradients, they typically lack single-cell resolution and are less compatible with integrated single-cell atlases. In our study, we adopted a segmentation-based approach to facilitate direct cell-level comparison between spatial transcriptomics and matched single-nucleus RNA-seq datasets. This strategy enables cell type annotation, label transfer, and atlas-level integration that remain essential for interpreting fetal and adult pancreas at single-cell resolution, despite segmentation-related limitations.

Cell segmentation applied to tissue remains a challenge in the context of spatial proteomics as well, primarily due to the heterogeneity of cell types with varying shapes and sizes. Fetal pancreas samples showed additional complexity due to high cell density, which hindered precision in defining the boundaries of individual cells. In our dataset, we introduced additional settings to enhance the acquisition of single-cell information from densely packed cells by tuning the definition of nuclei, accepting less spherical cells, and allowing for closer nuclei. Although we considered it critical to include different time points from fetal tissues on the same microscope slide to reduce batch-to-batch effects during sample preparation, this approach posed the challenge of establishing experimental conditions for samples with varying expression levels of certain proteins.

In addition to segmentation, spatial transcriptomics and proteomics are also limited by tissue sectioning. Although thin sections aim to capture a single-cell layer, in practice, they often intersect multiple vertical layers. As spatial data is typically visualized as a two-dimensional projection, transcripts from vertically stacked cells can be conflated, creating hybrid profiles that do not represent true biological cells. For instance, transcripts from the apical region of an alpha cell and the basal region of an adjacent beta cell may be merged, resulting in ambiguous or artifactual identities, particularly in tightly packed glandular structures such as pancreatic islets. These challenges complicate precise celltype annotation and downstream biological interpretation. Finally, while our cross-sectional dataset enables powerful inference of developmental and disease-related trajectories, direct lineage tracing or in vivo functional validation was beyond the scope of this study. Future longitudinal or perturbation-based studies will be needed to validate candidate regulators and lineage pathways identified here.

## Data availability

The sequencing data is deposited on EGA (Accession Number: XXX). An interactive visualisation for exploration and investigation of the data can be found on XXXX website

### Funding

This work was funded by the European Commission (EUHorizon2020 874710) and supported by additional funding sources as follows. E.M. was supported by the Ramón y Cajal fellowship RYC2021-032359-I, funded by the Spanish Ministry of Science, and by the Catalan Agency for Management of University and Research Grants (AGAUR, 2021 SGR 01586). D.B. was supported by the EMBO Long-Term Fellowship (ALT295-2019); the 2022 EFSD Rising Star Programme; the Ramón y Cajal program of the Ministerio de Ciencia e Innovación (RYC2021-033131-I); the Early Career Researcher grant Sigrid Jusélius foundation (@003701165704@); the Academy Research Fellowship 2024 Research Council of Finland (361593) and the Excellence Emerging Investigator Grant Novo Nordisk Foundation (NNF24OC0089232).

## Acknowledgments

We acknowledge the Spatial Proteomics Facility at the Royal Institute of Technology, funded by the Science for Life Laboratory and the National Microscopy Infrastructure, NMI (VR-RFI 2016-00968) for performing the PhenoCycler-Fusion experiments on the fetal pancreas. Human pancreatic islets were provided by the NIDDK-funded Integrated Islet Distribution Program (IIDP) (RRID:SCR_014387) at City of Hope, NIH Grant # 2UC4DK098085. We acknowledge the Scientific Computing of the IT Division at the Charité - Universitätsmedizin Berlin for providing computational resources that have contributed to the research results reported in this paper. This work was supported by the de.NBI Cloud within the German Network for Bioinformatics Infrastructure (de.NBI) and ELIXIR-DE (Forschungszentrum Jülich and W-de.NBI-001, W-de.NBI-004, W-de.NBI-008, W-de.NBI-010, W-de. NBI-013, W-de.NBI-014, W-de.NBI-016, W-de.NBI-022). We thank the Sequencing Facility of the Max Planck Institute for Molecular Genetics (MPIMG) for performing the sequencing runs of the adult and fetal pancreas samples. We thank Charité – Universitätsmedizin Berlin’s Research Data Management (Forschungsdatenmanagement) service for providing and supporting the OMERO platform. We are grateful to Prof. Dr. med. Wilko Weichert, who sadly passed away during the preparation of this manuscript, for his support and contributions.

## Competing interests

H.H. is co-founder and Chief Scientific Officer of Omniscope, a Scientific Advisory Board member at Nanostring and Mirxes, a consultant for Moderna and Singularity, and has received honoraria from Genentech. M.M. is co-founder and CEO at Single Cell Discoveries. A.v.O. is an Advisory Board member at Single Cell Discoveries.

## Author contributions

Conceptualization: E.M., D.B., L.T., F.C., E.d.K., J.F., B.G., H.H., E.L., L.P., K.S., A.v.O., W.W., C.C., R.E.; Software: J.L., S.T.; Formal Analysis: E.M., D.B., J.L., A.G., A.M.C., Ma.Ma., F.M., F.S., L.T., V.V., E.B., N.G., K.J., A.S., F.W.T., T.T.; Investigation: E.M., D.B., J.L., A.G., A.M.C., Ma.Ma., F.S., L.T., E.B., F.B., R.L.C., M.E., J.G., M.H., K.J., A.S., T.T., M.v.A.; Resources: E.M., D.B., F.C., E.d.K., J.F., B.G., H.H., E.L., L.P., K.S., A.v.O., W.W., C.C., R.E.; Data Curation: E.M., D.B., J.L., A.G., A.M.C., U.T., S.T.; Visualization: E.M., D.B., J.L., A.G., A.M.C., Ma.Ma.; Supervision: E.M., D.B., F.C., E.d.K., J.F., B.G., H.H., E.L., L.P., K.S., A.v.O., W.W., C.C., R.E.; Writing – Original Draft: E.M., D.B., J.L., A.G., A.M.C., Ma.Ma., V.V.; Writing – Review & Editing: L.T., J.F., B.G., A.v.O., C.C., R.E.; Project Administration: C.C., R.E.; Funding Acquisition: F.C., E.d.K., J.F., B.G., H.H., E.L., L.P., K.S., A.v.O., W.W., C.C., R.E.

## Materials and Methods

### Sample procurement of adult pancreas tissue

Pancreatic tissue procurement from adult organ donors was performed at Leiden University Medical Center (Leiden, Netherlands). The tissue acquisition was approved by the local clinical ethic committee and followed their guidelines. The samples were obtained in a standardized manner from four distinct locations (pancreas head, body, tail, processus uncinatus). For this purpose, three slices along the sagittal axis were taken from each organ, divided into one slice through the head of the pancreas, including the processus uncinatus, and one slice each through the central region of the pancreatic body and the pancreatic tail. The first slice from the head region was further divided into a cranial part hereinafter referred to as head and a caudal part representing the processus uncinatus. The grossing and sampling procedure was performed as fast as possible to ensure optimal tissue quality. The tissue from each individual location was further subdivided and depending on further downstream analysis either snap frozen with liquid nitrogen or fixated in neutral buffered formalin 10% and shipped to the corresponding consortium partners. Additionally, tissue microarrays (TMA) from the formalin fixated material of the organ donors were constructed. Of 17 organ donors only one was excluded completely from the here described analysis due to the detection of major pathological changes, resulting in 16 organ donors with high quality tissue samples.

Furthermore, from archived material from the Institute of Pathology of Technical University of Munich (Munich, Germany) we generated TMA blocks containing pancreas tissue with microscopic normal architecture on morphological level. These tissue samples derived from transplant cases or surgical specimens in which inconspicuous pancreas tissue could be identified distant from the pathological lesion. TMA generation and distribution was approved by the local clinical ethic committee (2022-207-S-NP). A complete list of samples is presented in Supplementary Table 1.

### Culture of primary human islets

Human pancreatic islets were obtained from organ donors through three sources: the Diabetes Research Institute at the IRCCS San Raffaele Scientific Institute (Milan, Italy), Leiden University Medical Center (Leiden, Netherlands) and Prodo Laboratories Inc. (Aliso Viejo, CA, USA). Islet isolation procedures were conducted in accordance with local ethical guidelines and approvals, specifically by the Human Research Ethics Boards at each respective site. The use of anonymized donor tissue was approved by the Clinical Research Ethics Committee of Hospital Parc de Salut Mar. In all cases, informed consent for research use of pancreatic tissue was provided by the donors’ families. Islets were shipped to Centre for Genomic Regulation (Barcelona, Spain) in CRML-1066 transport medium and, upon arrival, cultured at 37°C in a humidified incubator with 5% CO_2_. The islets were maintained in glucose-free RPMI 1640 medium supplemented with 10% fetal calf serum, 100 U/mL penicillin, 100 U/mL streptomycin, and 4.5 mM, 11 mM or 25 mM glucose, for a period of 72 to 96 hours prior to downstream processing.

### In vitro human islet glucose stimulated insulin secretion

Dynamic glucose-stimulated insulin secretion was assessed using a custom-designed perifusion system. The setup included an Ismatec 8-channel peristaltic pump (Model ISM931A, Ismatec), equipped with 2-stop, 0.38 mm Tygon tubing (Fisher Scientific), which directed the flow through a Biorep PERI-CHAMBER perifusion chamber. The outflow was routed via 1.016 mm Tygon tubing (Fisher Scientific) to a Biorep PERI-NOZZLE to ensure uniform drop collection. A total of 50 human islets were suspended in Krebs-Ringer Buffer maintained at 37°C. Perifusion was carried out at a constant flow rate of 250 μL/min, using buffer containing varying glucose concentrations (low glucose 3 mM, high glucose 15 mM) and additional secretagogues (10 nM Exendin 4, 30 mM KCl). Effluent samples were manually collected every 5 minutes into a 96 deep-well plate. At the conclusion of the perifusion, the islets were collected for total insulin content analysis. The islets were first sonicated in water for 20 seconds, then transferred to acidic ethanol (1.5% HCl in 100% ethanol). Insulin concentrations in the 5-minute effluent fractions and total insulin content were quantified using the Cisbio HTRF High Range Insulin Assay Kit (Cisbio 62IN1PEG), according to the manufacturer’s protocol. Measurements were performed in 96-well plates and read using a Tecan SPARK 10M multimode plate reader.

### Single cell/nuclei preparation

A rice-grain-sized piece of frozen tissue was cut with a scalpel from the frozen samples. For the fetal samples the whole piece was used. To isolate the nuclei we used the citric acid nuclei isolation protocol published by Tosti et al (18). In short, the frozen tissue piece was given into 1mL of pre-cooled S25 buffer inside a glass douncer tissue grinder (DWK Life Sciences, CAT. 357358). The tissue was crushed using a loose pestle and incubated for 5 min on ice. After repeating that step the tissue was homogenized with five more strokes of the loose pestle followed by five more strokes with the tight pestle. The suspension was transferred through a 35 µm strainer into a 1.5 mL Eppendorf tube and the strainer was washed with an additional 250 µL S25 buffer. The nuclei suspension was centrifuged at 500 g for 5 min at 4 ºC. Afterwards the supernatant was carefully removed and resuspended in 500 µL S25 buffer. After checking the nuclei number and quality the suspension was centrifuged as before or, if a lot of debris was visible, a density centrifugation was done instead. For this 500 µL of S88 buffer is carefully added to the bottom of the Eppendorf tube and centrifuged at 1000 g for 10 min at 4 ºC. Following removing the supernatant, resuspending in 300 mL S25 buffer and checking the number of nuclei, the suspension was centrifuged at 500 g for 5 min at 4 ºC. The supernatant was removed as much as possible and the nuclei were resuspended in the appropriate amount of resuspension buffer for the desired concentration. The nuclei suspension was diluted to load 10,000 nuclei per sample onto the 10x Genomics Chromium controller. After Gel Bead-in-Emulsion (GEM) generation and the RT-PCR, the GEMs were stored at 4 ºC until all adult samples were at this point.

For cultured human islets, approximately 2000 islets were harvested into 15 mL Falcon tubes, pelleted by centrifugation (200 rcf, 5 min), and washed with ice-cold phosphate-buffered saline (PBS). After removing the PBS, islets were enzymatically dissociated using a 1:1 mixture of TrypLE (Life Technologies, #12563029) and 0.05% Trypsin-EDTA (ThermoFisher, #25300054). The dissociation was carried out for 10 minutes at 37°C in a shaking water bath, with gentle pipette mixing every 3–5 minutes to facilitate breakdown into single cells. To stop the enzymatic reaction, an ice-cold solution of 5% fetal bovine serum (FBS) in PBS was added. The resulting cell suspension was filtered through a 33 μm strainer to remove clumps and extracellular DNA. Cells were then centrifuged at 250 rcf for 3 minutes, resuspended in ice-cold 0.04% bovine serum albumin (BSA) in PBS, and stained with Trypan Blue for viability assessment. Single-cell dissociation efficiency was confirmed microscopically, and cells were manually counted using a hemocytometer. Following dissociation and quality control, the single-cell suspension was aliquoted into separate tubes for scRNA-seq and scATAC-seq. For scATAC-seq, nuclei were isolated following 10x Genomics Nuclei Isolation for Single Cell ATAC sequencing protocol (CG000169). Briefly, single-cell suspension was resuspended in lysis buffer (10 mM Tris-HCl (pH 7.4), 10 mM NaCl, 3 mM MgCl2, 0.1%, Nonidet P 40 Substitute (Sigma 74385), 0.1% Tween-20 and 0.01% Digitonin, 1% BSA), and incubated on ice for 5 min. Lysis was stopped by adding 1 mL of chilled wash buffer (10 mM Tris-HCl (pH 7.4), 10 mM NaCl, 3 mM MgCl2, 0.1% Tween-20, 1% BSA). Nuclei were centrifuged 500 g for 5 min at 4ºC and resuspended in Diluted Nuclei Buffer.

### Single cell/nuclei library preparation and sequencing

Fetal: From each fetal sample 10,000 nuclei were loaded onto two lanes of a 10x Genomics Chip. The libraries were generated with the Chromium Next GEM Single Cell 3’Kit v3.1 (PN-1000268) following the corresponding protocol (CG000315).

Adult: Library generation was performed with the Chromium Next GEM Single Cell 3’ Kit v3.1 (PN-1000268) following the corresponding protocol (CG000315) was adapted to be automated with the Biomek i7 (Beckman Coulter, Life Sciences). After library generation, few libraries underwent an additional manual clean-up before they were sent out for sequencing.

Cultured islets: From each cultured islet sample, 7,000 nuclei were loaded onto a 10x Genomics Chromium controller channel, following manufacturers’ instructions. Library generation was performed with the Chromium Next GEM Single Cell 3’ Kit v3.1 (PN-1000268) following the corresponding protocol (CG000315) for scRNA-seq, and with Chromium Next GEM Single Cell ATAC Reagent Kits v1.1 (protocol CG000204) for scATAC-seq.

All libraries were sequenced on an Illumina NovaSeq600 (S4 flow cell, paired-end).

### Single cell/nuclei data pre-processing

The fastq-files of the sequenced samples were loaded into the One Touch Pipeline (OTP), which is an automation platform managing Next-Generation Sequencing (NGS) data and calling bioinformatic pipelines for processing these data (91). The used pipeline consisted of cellranger (v6.1.1) and the alignment to the GRCh38 genome.

### Single cell/nuclei adult and fetal pancreas snRNA ambient RNA removal

To reduce ambient RNA contamination, we applied CellBender (v0.2.2) (92) to all samples. For each sample, the parameters --expected-cells and --total-droplets-included were manually set. The number of expected cells was estimated from the Cell Ranger web summary, while the number of included droplets was determined from the knee plot in the Cell Ranger output, ensuring that cell-containing as well as a sufficient number of empty droplets were included for accurate background modeling. The model was trained for 150 epochs with a false positive rate (FPR) of 0.01. All downstream analyses were performed using the CellBender count matrix.

### Single cell/nuclei adult and fetal pancreas snRNA processing

Initial data analysis was performed using R (v4.1.2) and Seurat (v4.4.0) (93). Cells with fewer than 1,000 UMIs and fewer than 200 genes (for adult samples) and 300 genes (for fetal samples) were excluded as low-quality cells. Upper thresholds for number of genes per cell were determined individually based on their gene expression distributions, with an average cutoff around 6,000 genes per cell. Data were then normalized, log1p transformed, the top 2,000 variable genes were identified, scaled, and principal component analysis (PCA) was performed following standard Seurat workflows.

### Single nuclei doublet detection and removal in adult snRNA

Potential doublets were identified in each sample separately using two tools: DoubletFinder (v. 2.0.3) (94) and Scrublet (v. 0.2.3) (95), both run with standard parameters as described in their respective vignettes. Only cells that were identified by both tools as doublets were removed from the dataset.

### Single cell/nuclei integration

Fetal and adult samples were processed separately following the same integration workflow. For each dataset, individual samples were merged using Seurat’s merge() function. The merged objects were normalized, log1p transformed and the top 2,000 variable genes were identified. Data were then scaled and principal component analysis (PCA) was performed. Batch correction was applied using Harmony (v1.2.0) (96), using sample identity as the integration variable. Dimensionality reduction was performed with UMAP based on the Harmony-corrected components. Cell–cell neighborhood graphs were constructed, and clustering was performed across a range of resolutions (0.1 to 1.5) to identify transcriptionally distinct populations.

### Single cell/nuclei annotation of adult and fetal snRNA data

Cell type annotation was performed in three iterative levels of increasing resolution. Initially, a reference-based approach was used to guide broad classification. Specifically, label transfer using scOMM (19) was applied using the pancreas dataset (13) to generate preliminary cell type predictions. These initial labels were refined by identifying differentially expressed genes (DEGs) per Leiden cluster and manually assigning broad cell type categories (annotation level 1: epithelial, mesenchymal, immune) based on canonical markers. Each broad category was then subset into a separate object and reprocessed independently, including normalization, identification of variable genes, scaling, dimensionality reduction, clustering, and DEG analysis. This enabled finer classification into specific cell types (annotation level 2). Subsequently, each identified cell type was again subsetted, reprocessed, and analyzed to resolve cell states or subtypes using DEGs and known markers from the literature (annotation level 3).

Throughout this iterative process, additional doublets not identified during initial filtering were detected and removed. After completion of all three annotation levels, the fully annotated subsets were merged back into a single object for downstream analysis.

### snATAC adult whole pancreas processing

R (version 4.1.2) and Signac (v. 1.5.0) (97) were used for the initial snATAC data analysis. Aggregated samples were annotated using the hg38 annotation extracted from EnsDb.Hsapiens.v86 package (v. 2.99.0) annotation with the UCSC annotation style. Initial normalization and dimensionality reduction were performed according to the standard Signac pipeline. After an initial exploration, low quality cells were discarded with more relaxed parameters than the original pipeline (TSS enrichment >2 and % in reads >25). Then, the dimensional reduction was recalculated and Harmony (v. 1.0) was run over dimension 2 to 40 to correct sampling batch effect and clusters were calculated using the SLM algorithm available in the Seurat package (v. 4.1.0) (93) at default resolution value. The gene activity matrix was also calculated using the GeneActivity function available on Signac., with a 2000 bp extension of the upstream region.

### snATAC annotation of adult whole pancreas data

Based on the gene activity data, manual and automatic annotation was employed to assign biological identity to the clusters. Manual cell types were assigned according to expression of the markers described in Tosti et al. (13). Azimuth (v. 0.5.0) and the pancreatic AzimuthDB were used to contrast the performed manual annotation.

### snATAC doublet detection and removal in whole adult pancreas data

After the initial clustering, AMULET (v. 1.1) (98) as used to detect potential doublet cells and unwanted cells were removed. Furthermore, Acinar cluster 3 was removed since it presented a high proportion of doublets and accumulation of high-count cells, hinting at a technical cluster. Similarly, Acinar cluster 4 was also removed since only two cells remained.

### snATAC peak calling in adult whole pancreas

MACS2 (v. 2.2.7.1) (99) was used to call the peaks per each single identified acinar and ductal sub-state as well as the rest of annotated cell types individually. Afterwards, the different peak callings were performed, the results were merged into a single combined object. After the new set of peaks was calculated, the snATAC was rebuilt based on these peaks and the initial analysis was repeated till the cell types were newly annotated and the gene activity matrix recalculated step. Nevertheless, an acinar cluster had to be filtered out in the quality control step since it presented a low FRiP.

### snATAC and scRNA integration of adult whole pancreas data

To integrate the scRNA-seq and snATAC-seq data from whole adult pancreas samples the bridge integration available in the Seurat documentation was conducted. The multiome sample served as the bridge between both adult datasets. The first 30 dimensions were used for the integration (avoiding the first dimension in the case of snATAC data since it is heavily correlated to sequencing depth). Donors were considered as a batch variable for the integration. snATAC and scRNA label transferring between the adult whole pancreas datasets. ScoMM (19) was the software selected to transfer labels between both objects. To better bridge the gap between both modalities, an intermediate step was added by employing the multiomic data. For the RNA to ATAC label transfer, the RNA to multiome model was limited to the common marker genes identified in the scRNA. The model was trained with the following parameters: a single hidden node layer (1000 nodes), 30 epochs, a batch size of 32, relu activation, 0.001 learning rate, dropouts enabled and patience value of 20. Afterwards, the multiome to ATAC model was trained, common accessible peaks identified in the multiome following the RNA label transfer. The parameters for training this model were the following:two hidden node layers (1500 and 500 nodes), 30 epochs, a batch size of 32, a dropout rate of 0.7, relu activation, batch normalization and a learning rate of 0.001. For the snRNA to multiome transfer, a 70-30 split between training and validation was used while a 50-50 ratio was employed in the multiome to snATAC transfer.

### snATAC to scRNA label transfer from the adult non-diabetic endocrine sample

A direct label transfer between the snATAC gene activity matrix and the scRNA data was performed for the adult non-diabetic endocrine samples using ScoMM (19). In this case, three separate models were first trained and later stacked to bridge the gap between accessibility and transcriptomic data. Models were trained with default parameters except the dropout rate, set to 0.3, and the prediction threshold, lowered to 0.2. The selected models had a single layer of hidden nodes with 1300, 1400 and 2000 nodes. To stack these models into a single one, first a model fitting was required, where a validation split of 0.2 and a patience of 5 over 50 epochs was considered. Then, a stacked model with a single layer (1900 hidden nodes) was trained. A dropout percentage of 0.3 was considered and l2 was set to 0.01.

### Converting objects between Seurat and Scanpy

Most analyses were carried inside the Seurat and Signac environments. Nevertheless, some analyses were carried out with Python exclusive packages, such as STREAM. The object conversion between environments was done employing two main packages: sceasy (v. 0.0.7) to convert from Seurat to H5AD and SeuratDisk (v. 0.0.0.9015) to convert from H5AD to H5Seurat.

### snATAC trajectory analysis for the adult whole pancreas data

For the trajectory analysis two groups of cells were subset from the main whole pancreas adult snATAC object: Acinar 4 subcluster and Ductal cell simultaneously expressing CPA1, PRSS1 and CELA3A, three known acinar TFs. The Seurat object containing these cells was then rescaled and the gene matrix was recalculated. The best dimensional reduction was obtained considering the first 15 UMAP components calculated over 20 PCA dimensions corrected with harmony. Afterwards, the clustering was performed with the Leiden algorithm and a 0.5 resolution. The object was then transformed to H5AD format and trajectories were inferred with the STREAM package (v. 1.1) (21). Based on the dimensional reduction calculated with Seurat, 10 initial clusters were set for the elastic principal graph, which was optimized with the following parameters: epg_alpha=0.01, epg_mu=0.05, epg_lambda=0.01. Afterwards, branching was recalculated by adding 65 nodes to the elastic principal graph. Finally., the elastic principal graph branches were collapsed based on the number of edges (epg_collapse_ mode=‘EdgesNumber’, epg_collapse_par=2).

#### snATAC and snRNA adult pancreatic network

To build the snRNA and snATAC network a subsample of 60k cells was generated based on the snATAC data. Additionally, in order to balance the proportion of cells, the number of acinar cells selected was equated to the number of ductal cells. After subsetting the snATAC data, both snATAC and snRNA objects were transformed to python objects. Once data was transformed, SCENIC+ (v. 1.0) (100) was employed to generate the adult pancreatic network and downstream analysis. Pipeline was run according to the available tutorial for unpaired data modality. Five cells were considered for each pseudo-cell built by the software and the downstream tutorials were used as a guide for further analysis.

#### Multiome sample processing, merging and annotation

An individual Seurat object was built for each of the two samples. Similar to the adult whole pancreas scATAC data, the hg38 annotation from the EnsDb.Hsapiens.v86 package was used for annotation, with the UCSC annotation style. However, only the peaks previously identified in the whole adult pancreas were kept to maintain consistency between datasets. Quality controls were performed according to the standard Signac pipeline and cells with less than 50000 ATAC counts, less than 25000 RNA counts and a TSS enrichment higher than two were kept. Once both objects were evaluated, they were merged into a single object, and after additional QC evaluations, cells with lower than 300 ATAC counts and 1000 RNA counts were discarded. Data was then normalized using SCTransform for the scRNA information and a PCA and UMAP (40 dimensions) was built. For the scRNA data, a UMAP reduction was built with dimension 2 to 40 of LSI reduction. Finally, a multimodal nearest neighbor matrix was used to generate a multimodal UMAP visualization and cells were clustered at 0.8 resolution and the gene activity matrix was calculated.

### Pseudotime alignment between snATAC and snRNA data

Pseudotime values extracted from the trajectory analysis with STREAM (21) were added to the corresponding snRNA/snATAC object. From the snRNA object, cells in the S0 to S1, S1 to S2 and S2 to S3 were selected. From the snATAC data, cells prior to the branching event in S3 were selected regardless of the branch they belonged to. After the cells were selected, pseudotime values were scaled between 0 and 1. Expression matrices were limited to the unique top 50 branch markers per each of the branches described before, sorted by log2FC and q-value. Finally, global alignment was performed by running cellAlign v.0.1.0 according to the default parameters.

### Multiome cell type annotation

Manual and automatic cell annotation were employed. For the manual annotation, the set of markers reported in the Tosti et al. (13) paper allowed us to define the different cell types. Automatic cell annotation through DeepSCore was performed to revise the manual annotation. The dataset published together with the Tosti et al. (13) paper served as a reference for label transferring. Similarly, our own adult whole pancreas scATAC dataset was employed as a reference for the label transferring of the scATAC modality.

### Multiome doublet identification

Three different software were used for doublet detection: AMULET for the scATAC data and Scrublet (v. 0.2.3) (95) and DoubletFinder (v. 2.0.3) (94) for the sn/scRNA. Each software was run per sample and the different predictions evaluated. The predictions from AMULET and Scrublet were kept for filtering out double positive cells for further analysis.

### Consensus endocrine marker identification across sn/ scRNA-seq datasets

Consensus markers were calculated for endocrine cells based on the methodology described by Van Gurp et al. (15). First, pairwise DEA analysis was performed for all the endocrine cells in the different datasets. Afterwards, for each individual comparison, the results across the datasets were integrated, which were then further integrated by target cell type in order to procure the final list of DEGs per cell type across datasets.

### MILO analysis

We applied the MiloR R package (v1.6.0) (55) to perform differential abundance (DA) testing on endocrine cell population from cultured human islets dataset. A k-nearest neighbor graph was computed on harmonized PCA embeddings, and Milo neighborhoods were constructed using buildNhoodGraph(). We tested for differential abundance between type 2 diabetes (T2D) and non-diabetic (ND) donors using testNhoods() and patient-level metadata. Spatial FDR correction was applied, and neighborhoods with FDR < 10% were retained.

To group spatially adjacent neighborhoods with similar DA signals, we used groupNhoods() with max.lfc.delta = 3. To assign DA groups back to single cells, we used the Milo@ nhoods matrix to determine which cells belonged to which neighborhoods. For each cell, we identified all associated neighborhoods, retrieved their corresponding Nhood-Groups, and recorded the groups in a binary matrix.

To assign cells to disease-associated groups, we examined all neighborhoods each cell belonged to and determined whether they were part of T2D- or ND-associated Nhood-Groups (ND: groups 1, 4, 6, 11, 12, 13, 16; T2D: groups 2, 5, 8, 10, 15). Cells with neighborhoods assigned to both sets were labeled as “mixed” in a new Milo_T2D_ND_clean column. Only cells consistently associated with either T2D or ND were used for differential gene expression analysis. We then used Seurat’s FindMarkers() function to compare gene expression between T2D- and ND-associated alpha, beta and delta cells (based on Milo_T2D_ND_clean), using Wilcoxon rank-sum tests with default filtering (logfc.threshold = 0.0, min.pct = 0.1, only.pos = FALSE).

### Xenium in situ sample preparation and sequencing

For the experiment the pre-designed Xenium Prime 5K Human Pan Tissue & Pathways Panel (PN-1000671) as well as additional 100 custom probes (PN1000766, Supp. Table 3) were used. Tissue sectioning and sample preprocessing were performed according to the manufacturer’s protocol.

Briefly, 5-μm FFPE sections were mounted on Xenium slides and dried to optimize adherence. Subsequently, the samples went through a 3-day protocol (CG000760, Rev C) including multiple steps such as priming hybridization, RNAse treatment, polishing, probe hybridization, ligation, amplification and applying cell segmentation staining reagents. The pre-processed samples were analyzed using the Xenium Analyzer with the instrument software version 3.1.0.0 and analysis version 3.1.0.4. Subsequently, Hematoxylin & Eosin (H&E) staining of the tissue sections was performed according to the manufacturer’s protocol for post-Xenium stainings.

### Xenium data processing

The analysis was performed in Python (v3.12.8) using scanpy (v1.10.4) (101), squidpy (v1.6.2) (102), and spatialdata (v0.3.0) (103), with numpy (v1.26.4) (104) and pandas (v2.2.3) for data handling. Visualizations were generated using seaborn (v0.13.2) (105) and matplotlib (v3.10.0) (106). To facilitate labelling of individual cores, the images from the Xenium experiment were uploaded to Omero, a web server based image repository, where a region of interest (ROI) around each core on each TMA was annotated and uniquely labeled. After loading the Xenium data using spatialdata the ROI coordinates were imported using ezomero (v3.1.0) (107), to assign each cell to their corresponding TMA core. For QC all TMAs were merged after ensuring unique cell IDs. Cell identification was based on the Xenium segmentation output and cells with less than 10 detected transcripts as well as transcripts expressed in less than 5 cells were removed. The samples were normalized to the median of total counts followed by log1p transformation and dimensional reduction. Fifty principal components were used for integration with HarmonyPy (v0.0.10), followed up by nearest neighbour construction and UMAP visualization. In the end the object with all TMAs consisted of 1,585,738 cells.

### Xenium H&E image alignment

To achieve accurate alignment between H&E-stained images of the corresponding DAPI-stained sections, we implemented a multi-step registration pipeline using scikit-image (v0.25.2) (108). First, downscaled images for both modalities were used for feature detection with Scale-Invariant Feature Transform (SIFT) algorithm (maximum ratio of 0.8) followed by robust model estimation using the RANSAC algorithm (residual threshold of 2 pixels and 1000 maximum trials) to estimate an affine transformation between the images. This transformation was then adapted and applied to the full-resolution H&E image. To further refine the alignment, ROIs were manually selected across the images, and nuclei within these ROIs were segmented using StarDist (v0.9.1) (109–111). SIFT was then used to extract additional common features from the segmented nuclei and a final affine transformation was estimated using RANSAC. The aligned H&E images were saved in the OME-TIFF format.

### Xenium cell type assignment

**S**imilar to the single-cell data analysis, this was an iterative process. First, Leiden clusters were computed, followed by the identification of DEGs. These DEGs were used to identify marker genes reported in Tosti et al. and our single-cell dataset, allowing us to broadly annotate the cells into Acinar, Ductal, Endocrine, Mesenchymal, and Immune categories (annotation_l1). Next, cells sharing the same annotation were subset. Each subset was normalized, log1p-transformed, subjected to dimensionality reduction (PCA), integrated using HarmonyPy (v0.0.10), followed by nearest-neighbors computation, Leiden clustering, and DEG analysis. Using the DEGs a finer annotation of each subset was performed (annotation_l2). The endocrine compartment received an even finer annotation based on their DEGs, which is reflected in ‘annotation_l2.5’.

The new labels were transferred into the object consisting of all TMAs and using Sopa (v2.0.0) (112) exported into a format compatible for visualization with 10x Genomics Xenium Explorer 3.

### Xenium pCAC identification

Using the H&E stained images a pathologist manually annotated in three TMAs a total of ∼200 cells that based on location and phenotype were centroacinar cells. Annotations of the centroacinar cells were imported (geojson v3.2.0) and their coordinates were transformed to match the spatial transcriptomics data (see Xenium H&E image alignment.) Cells overlapping with the annotations were labelled as ‘centroacinar cells’. The final transformed annotations were integrated into the AnnData object and the 258 cells were visualized using Xenium Explorer. Due to imperfect cell segmentation, we manually selected intact cells in Xenium Explorer and annotated them using their cell IDs in the AnnData object as ‘CAC’ in the ‘annotation_l2_cac’ meta.data column. This reduced our 258 detected cells to 198 intact cells. This label was used to compute DEGs, from which we retained genes with a positive log fold change and an adjusted p-value < 0.05. These genes were used to compute a robust module score using Scanpy’s score_genes() function, which was run 100 times with different random seeds, and cells within the top 5% of scores across all iterations were selected. The same approach was applied to DEGs from cluster 5 from the scRNA ‘centro-acinar’ object (see Fig. 2D), but before the robust module score was calculated the genes were filtered to retain only genes detected in the Xenium experiment. Finally, the intersection of the top 5% scoring cells from both robust module scores was used to classify plastic centroacinar cells (pCAC).

#### Xenium ionocyte identification

For identification of the ionocyte population in the Xenium data we also used the robust module score approach utilizing Scanpy’s score_genes() function. As input for the score the genes identified for the ionocyte population (also see Supp. Fig. 3D) were used, namely: HEPACAM2, FOXI1, KIT, TFCP2L1, PLCG2, CLNK, DMRT2, ATP6V1B1, ATP6V1B1, ATP6V1C2, KCNMA1, KRT19 and CFTR. Only the top 0.1% cells were classified as potential ionocytes and visualized in the Xenium dataset (also see Supp. Fig. 3E).

### Xenium multipotency score

The multipotency score is calculated using the AddModuleScore() function from Seurat. As input multipotency genes that were expressed in the snRNA cluster 5 (see Fig. 2D) were used, namley: PDX1, NKX6-1, SOX9, HNF1B, FOXA2, RHOV, MNX1, INSM1, ST18, PAX6, JAG2, DLL1, HEY1, HES1, NTRK2, SLC4A4, TM4SF1, KRT19, LYZ, SPP1, H19, FXYD2, MKI67, TOP2A, PCNA, DLK1, SHISA2, SOX4, TFF1, KLF5, CLU and AGR2 (78-80).

### Xenium islet composition analysis

The xenium AnnData object was subset to only include endocrine cells. To identify spatially distinct islet structures, we computed spatial neighborhoods using squidpy’s spatial_neighbors() with radius=20 µm, grouping cells by TMA using the library_key parameter. A spatial connectivity graph was constructed and used to identify spatial clusters via connected component analysis (scipy.sparse.csgraph.connected_components) so that each cell was assigned to a spatial cluster.

For each spatial cluster, summary statistics were computed including total cell area, nucleus area, x/y coordinates, and cell count. Clusters with a total islet area below 1000 µm^2^ were defined as “small islets,” while the remaining were considered “large islets.” Differential gene expression between small and large islet-associated cells was performed using Scanpy’s rank_genes_groups() with the Wilcoxon ranksum test, comparing the small islet group against the large islet group.

### CODEX Immunohistochemistry staining & slide scanning

Sections (4 µm thick) of a TMA block containing 102 cores from adult pancreas FFPE archived material underwent deparaffinization in xylene, followed by hydration in graded alcohols and blocking for endogenous peroxidase in 0.3% hydrogen peroxide diluted in 95% ethanol. The next step was Heat-Induced Epitope Retrieval (HIER) treatment, using pH 6 EDTA buffer (Lab Vision, Freemont, CA) in a decloaking chamber (Biocare Medical, Walnut Creek, CA) for 4 min at 125 °C, after which the samples were allowed to cool to 90°C. The automated immunohistochemistry was performed as previously described (113) using anti-SOX9 antibody (Atlas Antibodies, Cat. No. AMAb90795, dilution 1:1500) and anti-ANXA13 antibody (The Human Protein Atlas, Cat. No. HPA019650, dilution 1:1000). The stained slides were scanned using a Digital Slide Scanner Axioscan 7 (ZEISS), equipped with a Hitachi 203 camera at 20x with resolution of 0.22µm/pixel. The images were manually annotated by two different experts.

### CODEX Indirect immunofluorescence staining & antibody conjugation

Prior to in-house conjugation of antibodies with a unique oligonucleotide sequence for multiplexing experiments, the specificity of the antibodies was assessed by performing single indirect immunofluorescence staining (114) on single pancreatic tissue sections from a FFPE block. The block was sectioned to a thickness of 5 µm, mounted on positively charged microscope slides (VWR, Epredia, Cat. No. 76406-502), and baked at 55 °C for 25 min. Each tissue section was deparaffinized by immersing it in HistoChoice (Sigma, Cat. No. H2779), for 5 min twice, and then rehydrated through a series of ethanol solutions (100% ethanol twice, followed by 90%, 70%, 50%, 30%, ddH2O twice; each step lasted 5 min). The next step was Heat-Induced Epitope Retrieval (HIER) treatment, using pH 6 EDTA buffer (Akoya Biosciences, Cat. No. AR900250ML) in a pressure cooker (Bio SB TintoRetriever, Cat. No. BSB-7087) at a temperature of 114–121 °C, high pressure, for 20 min. Every tissue sample was equilibrated at room temperature, followed by two 2-minute incubations in ddH2O. To minimize tissue auto-fluorescence, each tissue section underwent a photobleaching protocol where the slide was placed in a 50 ml falcon containing 25 ml 1x PBS (Gibco, Cat. No. 18912-014), 4.5 ml 30% H2O2 (Sigma, Cat. No. 216763) and 0.8 ml 1M NaOH (Sigma, Cat. No. S5881). The section in solution was exposed to two LED lamps (GLIME, Light Therapy Lamp UV Free 32000), positioned on either side and incubated for 45 minutes. A new photobleaching solution was prepared and replaced for an additional 45-minute incubation under the LED lamps, followed by four washes with 1x PBS washes, each step for 2 min. Each primary antibody tested was diluted in antibody diluent containing 0.3 % Triton (Sigma Aldrich, T8787) 1x PBS pH 7.4. The diluted antibody was then applied to the tissue section to be incubated overnight at 4°C. The next day, the material was washed with TBS-T three times, with each wash lasting 15 minutes. To prevent nonspecific binding, the tissue was incubated for 30 min in TNB buffer containing 0.1 M Tris-HCl/0.15 M NaCl and 0.5% blocking reagent pH 7.5 (Akoya, SKU FP1020). Depending on the species reactivity, the secondary antibodies used were goat-anti-rabbit IgG (H+L) Highly Cross-Adsorbed secondary Antibody, Alexa Fluor™ 647 (ThermoFisher, Cat. No. A-21245) or goat-anti-mouse IgG (H+L) Highly Cross-Adsorbed secondary Antibody, Alexa Fluor™ 555 (ThermoFisher, Cat. No. A-21424). The secondary antibody was diluted 1:800 in TNB containing nuclear staining Hoechst (ThermoFisher, Cat. No. H3570) and added to the tissue sections for a 90-minute incubation at room temperature. Finally, the stained material was mounted with Fluoromount-G (Thermo Fisher Scientific, 00-4958-02) on a coverslip for image acquisition.

Each single tissue section was image acquired using an inverted DMi8 microscope (Leica Microsystems, Mannheim, Germany), equipped with a digital CMOS camera (ORCA-Flash 4.0 V3, Hamamatsu), employing a 20x/0.75 NA dry objective with a resolution of 0.325µm/pixel.

The antibody was selected for oligonucleotide conjugation based on the following criteria: 1) The staining pattern was as expected (using The Human Protein Atlas database and antibody datasheets as references); 2) The signal-to-noise ratio was higher than 3. The custom antibody conjugations were carried out in accordance with the CODEX protocol from Akoya, User Manual Rev. C.

### CODEX non-disease adult and fetal highly multiplexed staining

The highly multiplexed panels include antibodies pre-conjugated by Akoya Biosciences and in-house conjugated antibodies (Supp. Table 1: adult_antibody_panel_CODEX, fetal_antibody_panel_CODEX). The multiplex antibody staining protocol used for the adult and fetal pancreas tissues was the same as that described by (115), in accordance with the CODEX protocol from Akoya, User Manual Rev. C, with the exception of the glass format due to the instrument version: the former version is referred to as CODEX, while the newer version is referred to as PhenoCycler Fusion; for simplicity, we will refer to it as CODEX. Tissue MicroArray (TMA) adult sections were placed on glass coverslips 22x22mm (Marienfeld, Cat. No. 102052) coated with 0.1% poly-L-lysine (Sigma, Cat. No. P8920): 184 tissue cores divided into 9 coverslip (Supp. Table 1: TMA_adult_CODEX). Whereas the FFPE fetal materials were sectioned and mounted on positively charged microscope slides 75×25 mm (VWR, Epredia, Cat. No. 76406-502): 11 tissues divided into 2 different slides (Supp. Table 1: TMA_fetal_CODEX). Each FFPE tissue section underwent sample preparation as detailed in the previous section. After the material was treated with the photobleaching protocol, each glass was then transferred to a new 6-well plate (for coverslip format)/ Coplin Jar (for microscope slide format) containing Hydration Buffer (Akoya Biosciences, Cat. No. 7000017) and incubating for 2 min, repeating this step twice. Following this, the glass was placed in a reservoir with a Staining Buffer (Akoya Biosciences, Cat. No. 7000017) and equilibrated for 30 min. During this time, the antibody cocktail solution was prepared by diluting all the oligonucleotide-barcoded antibodies in Staining Buffer containing N, J, G, and S blockers (Akoya Biosciences, Cat. No. 7000017). Once prepared, the mixed antibodies were applied to the tissues for an overnight incubation at 4 °C. The following day, the antibody cocktail solution was removed from the tissue by washing with Staining Buffer, followed by three different consecutive post-fixation steps with washes between them: 1) Incubation in 1.6% paraformaldehyde (ThermoFisher, Cat No. 043368) diluted in Storage Buffer (Akoya Biosciences, Cat No. 7000017) for 10 min incubation at room temperature; ncubation in ice-cold methanol (Sigma, Cat No. 322415)for 5 min at 4 °C; 3) Incubation in Fixative Solution (Akoya Biosciences, Cat No. 7000017) diluted at 1:50 in 1x PBS. Each glass was then stored in a Storage Buffer until the image acquisition step. Per each experiment, a 96-well reporter plate was prepared, containing a fluorescent dye conjugated to an oligonucleotide sequence complementary to one specific antibody barcode. Unique reporters were added in groups of up to three spectrally different dyes, including AF488, Atto550, and Cy5 (Akoya Biosciences), along with nuclear staining in each single well.

### CODEX Adult - image acquisition & computational image processing

Image acquisition was performed on the CODEX instrument (currently the PhenoCycler-Fusion system, Akoya Biosciences, Marlborough, MA), connected to an inverted DMi8 microscope (Leica Microsystems, Mannheim, Germany). Images were acquired with a digital CMOS camera (ORCA-Flash 4.0 V3, Hamamatsu), employing a 20x/0.75 NA dry objective with a resolution of 0.325 µm/pixel, Z-steps = 5 of z = 1.5 μm. The light source used was a SOLA-SM-II. The CODEX Instrument Manager (version 1.30.0.12) was utilized to control CODEX and integrate the microscope control software. The generated 16-bit .tiff images were processed using CODEX® Processor application (version 1.7.0.6), which performed image alignment across cycles, tile stitching, background subtraction, deconvolution, extended depth of field, and shading correction.

Every single tissue core from each TMA image/section (TMA n = 9) was cropped, and the images were cell segmented separately using our in-house developed toolkit PIPEX, as described in the previous section. Single cells were cell segmented based on Hoechst nuclear staining (nuclei_diameter=20px, nuclei_expansion=20px); this segmentation was refined using the CDH1 (E-cadherin) membrane marker (membrane_diameter=25px). Direct calculations of segmented cell data (mean pixel intensities, markers binarization) were generated from each tissue core and saved in a Comma-Separated Value (CSV) file for downstream analysis, where additional metadata was included.

### CODEX Adult - data cleaning, clustering & cell annotation

Data cleaning, clustering, cell annotation, and analysis were primarily conducted using R version 4.2.2 (RStudio) and Python version 3.12 (Jupyter Notebook and PyCharm), along with Perseus software (version 2.0.10.0). We combined datasets from nine tissue microarray (TMA) slides, resulting in a comprehensive dataset of 1,850,689 segmented cells. This dataset underwent filtration to remove technical artifacts, yielding a refined total of 1,756,482 cells. The filtration process involved excluding cells based on size, specifically removing those in the first and 100th percentiles, followed by filtering out the top 1% based on Hoechst, CTNNB1, and CDH1 intensity.

Next, we applied the log1p transformation to the data and conducted batch correction using the ComBat method from the “sva” R package. We then performed quantile normalization using the “preprocessCore” R package to ensure data harmonization.

For cell type annotation, we utilized Leiden clustering, which categorized the cells into distinct types as follows: acinar (n=846,840), ductal (n=303,139), endothelial (n=142,586), endocrine (n=65,548), MHC I+ (n=77,806), proliferative (n=24,266), Schwann (n=5,198), stromal (n=256,633), and other (n=38,419). To identify subpopulations of endocrine cells, we isolated the endocrine dataset and employed automated Otsu thresholding for annotation using the scikit-image Python library (version 0.22.0). This analysis revealed distinct subpopulations of endocrine cells, including alpha (n=16,883), beta (n=12,856), delta (n=495), alphabeta (n=3,783), betadelta (n=1,373), polyhormonal (n=2,014), and “other” (n=28,144). The “other” category included cells neighboring GCG+ cells, which were excluded from analysis due to technical staining artifacts that led to signal bleed-through.

For cell clustering and unsupervised analysis, we utilized the “scanpy,” “squidpy,” and “numpy” packages in Python (101, 102, 104), with visualizations, including a Sankey plot, generated using the Plotly library. For spatial analysis, we calculated Euclidean distances between annotated cell populations using Perseus software, and we visualized the results using the “matplotlib” and “seaborn” libraries in Python (105, 106). Additionally, the expression of endocrine markers was illustrated in a dot plot format using the “matplotlib.pyplot” library.

### CODEX Fetal - image acquisition & computational image processing

Image acquisition was performed on the PhenoImager Fusion 2.2.0 instrument (Akoya Biosciences, Marlborough, MA) where the three fluorescent oligo reporters were applied to the tissue in iterative imaging cycles. The images were captured using a widefield microscope with an Olympus UCPlanFL 20x/0.70 NA dry objective with a resolution of 0.51 µm/pixel. Whole-slide images were automatically aligned using DAPI as a reference, and the background was subtracted to be assembled into the final 8-bit QPTIFFs image files.

Every tissue from each QPTIFFs image file (n = 2) was cropped and converted into a separate .tiff for each marker channel. These images were cell segmented using our in-house developed toolkit PIPEX (116). The subsequent strategy was followed to be able to obtain single-cell quantification: 1) Nuclei mask: Stardist ML (111) model was applied on the DAPI images (nuclei_diameter=10px), selection nuclear expansion (nuclei_expansion=20px) and additional settings to be able to obtain single cell information in these tissues with tightly packed cells (nuclei_definition=0.1, nuclei_closeness=0.3; 2) Membrane mask: a watershed algorithm was run using CDH1 (E-cadherin) images (membrane_diameter=15, membrane_compactness=0.7) to obtain the membrane boundary shape information. The nuclei mask expansions were cut to fit the membrane boundaries calculated in the membrane mask. Direct calculations of segmented cell data (mean pixel intensities, markers binarization) were generated from each tissue and saved in a Comma-Separated Value (CSV) file for downstream analysis, where additional metadata was included.

CODEX Fetal - data cleaning, clustering & cell annotation The data cleaning step was performed using Python (v. 3.10) in the PyCharm IDE (v. 2024.2.5). Per each single tissue, a sanity filtering step was performed using cell size to discard segmented cells that were too small or too large; and DAPI intensities. Visual inspection of each image per marker, including the cell segmentation mask, facilitated the removal of staining artifacts. It also allowed for exclusion of regions that were presumably not part of the pancreas tissue based on expected canonical protein expressions. The cleaned data were input into PIPEX software for data normalization (Min-Max), following the merge of all the CSV files for batch correction (ComBat) based on the slide number. The combined datasets from the 11 tissue sections included 224,713 cells. For further analysis in PIPEX, k-means (k = 10) clustering using ACTA2, CD31, CD45, CHGA, C-Pep, E-cadherin, GCG, HLA-ABC, HLA-DR, Pan-Cytokeratin, RBP4, SST, SYP, TTR, VIM as markers, was performed separately for each week and cell type annotations were assigned based on ranked protein expression and visual image inspection following annotation_l1 categories. Following the annotations, clusters corresponding to the same cell type were combined. A more in-depth analysis of the endocrine population included 18,741 cells, each cluster annotated with an endocrine marker was merged and then re-clustered using k-means using CHGA, C-Pep, GCG, Pan-Cytokeratin, RBP4, SST, SYP, TTR as markers. Supplementary data contain whole tissue images with the corresponding cell annotations. The Min-Max normalized data were used to create violin plots (PanCytokertain and CHGA), and the binarized data from Ki67 expression were used to generate a horizontal bar plot categorized by different ages.

