## Supplementary figures for "Latent plasticity of the human pancreas across development, health, and disease"

### Supplementary Fig 1.

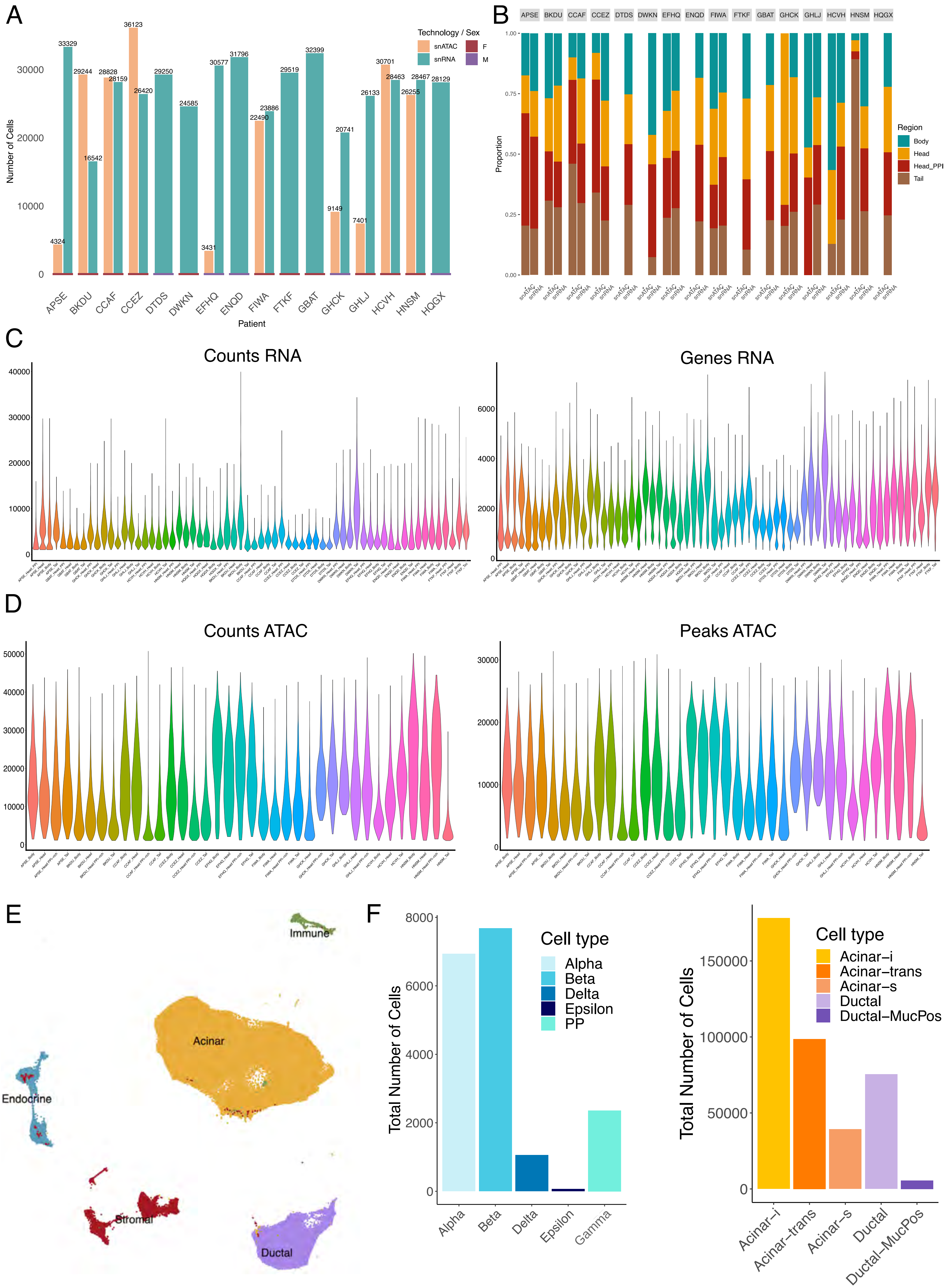

***Supplementary Fig. 1.***

**A.** Breakdown of the snATAC and snRNA adult datasets according to donor and sex of the donor.

**B.** Breakdown of the number of cells per donor according to the sex of the donor and the cellular region of origin, respectively.

**C.** Quality control metrics for the reads sequenced per donor split by region of origin in the snRNA adult dataset.

**D.** Quality control metrics for the reads sequenced per donor split by region of origin in the snATAC adult dataset.

**E.** UMAP showing the annotated broad cell types in the adult snRNA dataset.

**F.** Available number of cells in the exocrine and endocrine compartment, respectively, at a deep annotation level.

Supplementary Figure 2

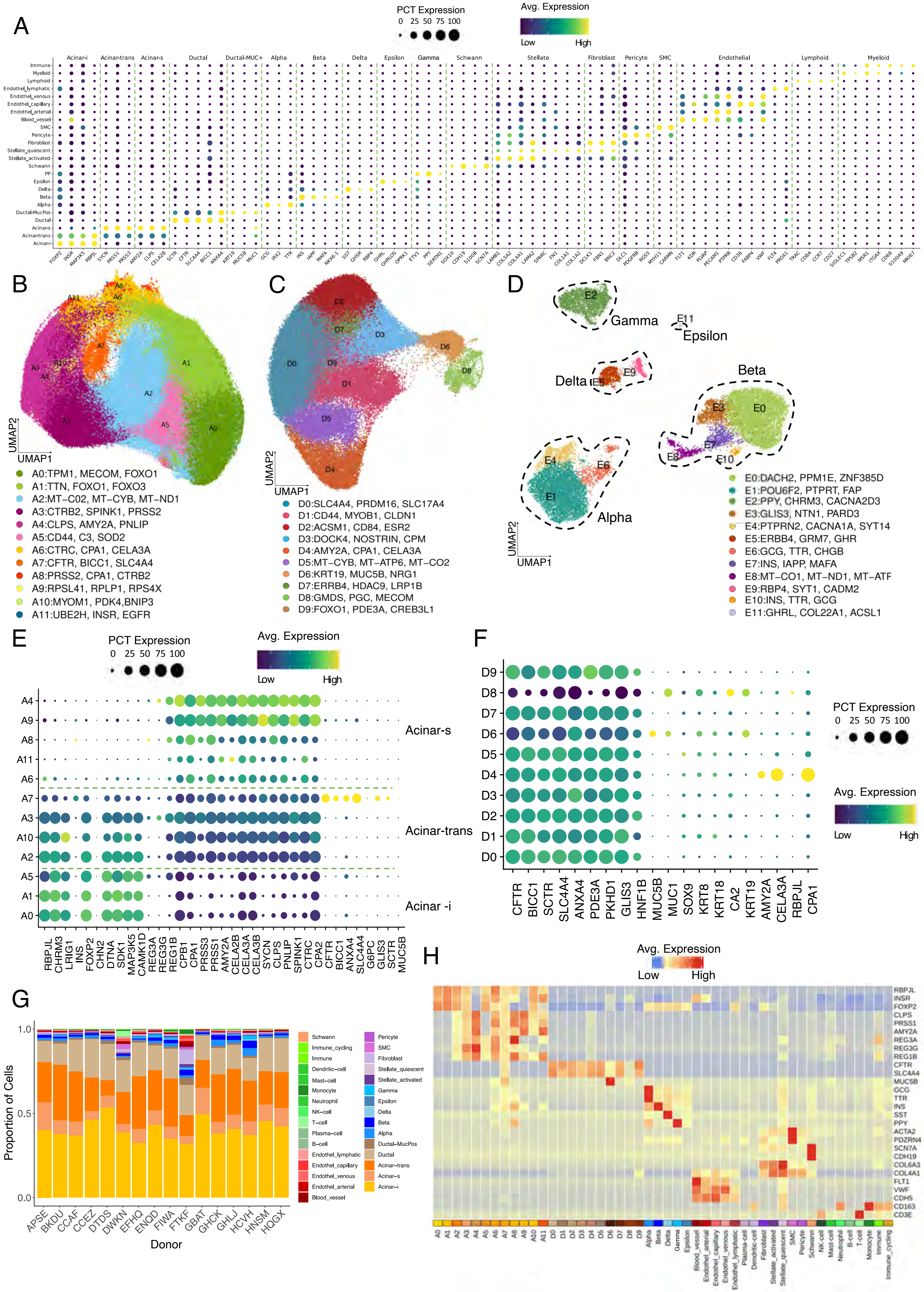

***Supplementary Fig. 2.***

**A.** Dot-plot of the markers selected for the cell type annotation process.

**B-D.** UMAP representations of 316,117 acinar cells (B), 81,047 ductal cells (C), and 18,143

endocrine cells (D) from the complete adult dataset. Unsupervised clustering analysis is displayed, with clusters annotated based on specific gene markers. The top cluster-specific markers are listed in the legend.

**E.** Dot plot showing the expression of specific acinar gene markers used to define three main acinar states: Acinar-i, Acinar-s, and Acinar-trans, as referenced in Fig. 1D. Ductal markers have been included to reflect the plastic state of A7.

**F.** Dot plot showing the expression of specific ductal markers and acinar markers reflecting the plasticity of D4.

**G.** Summary of the cell types annotated per donor at a deep annotation level in the snRNA adult dataset.

**H.** Heatmap of the main cell type markers expression for the different cell types and states (for acinar and ductal cells mainly) identified in the adult snRNA sample.

Supplementary Figure 3

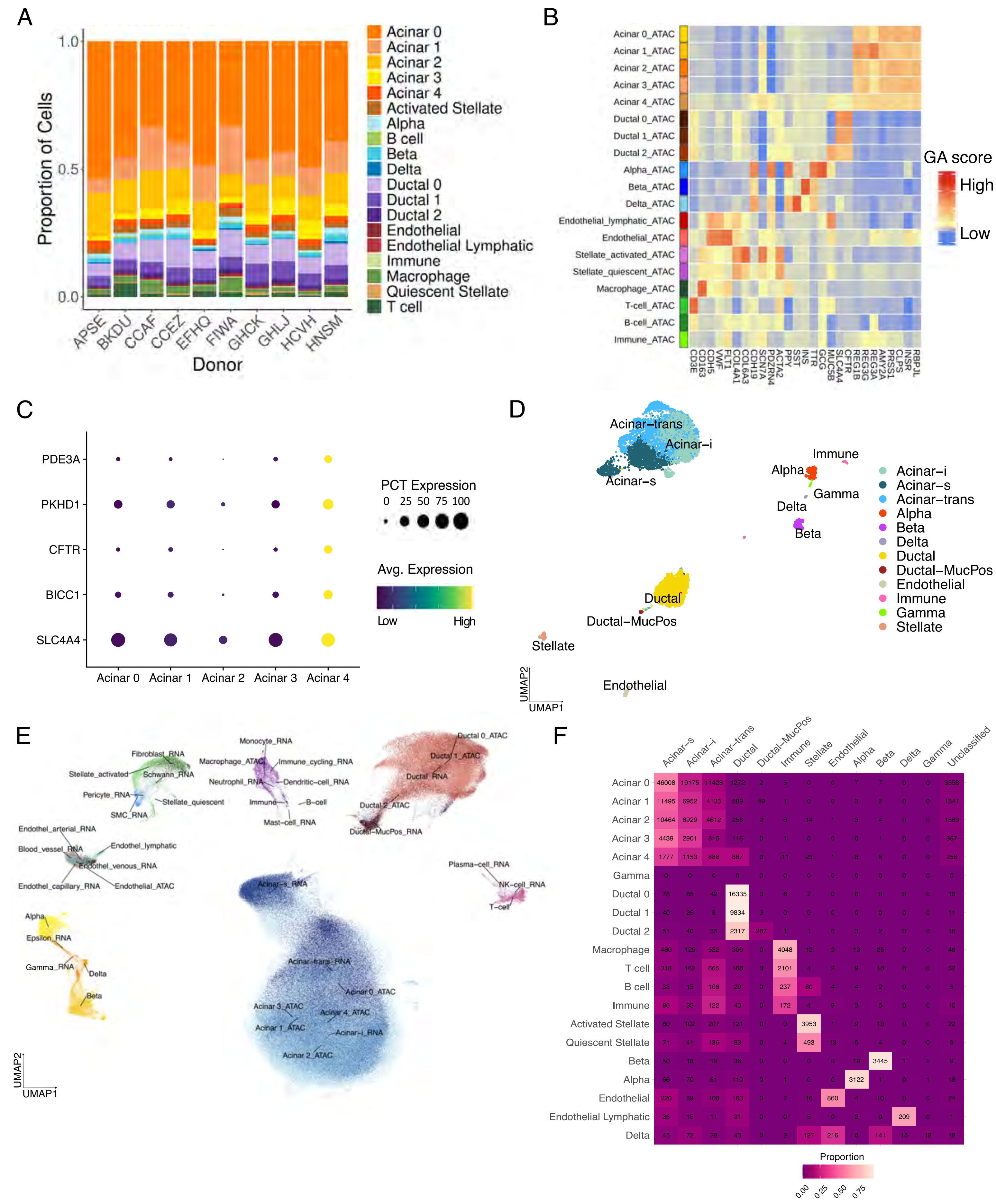

***Supplementary Fig. 3.***

**A.** Abundance of the different identified cell types and substates per snATAC adult donors.

**B.** Heatmap of the gene activity expression values for the canonical markers of the identified cell types in snATAC adult data.

**C.** Dot plot highlighting the relative expression of specific ductal gene markers, with higher

expression observed in the Acinar 4 cluster.

**D.** UMAP of the single-nuclei multiomics dataset with cell-type annotations aligned to the reference snRNA-seq of the adult pancreas in Fig. 2A.

**E.** UMAP of 636,341 cells integrated from snRNA-seq and snATAC-seq datasets, using multiome data as a bridge. Colors represent cell types or states across both datasets.

**F.** Correlation between snRNA-seq and snATAC-seq cell clusters.

Supplementary Figure 4

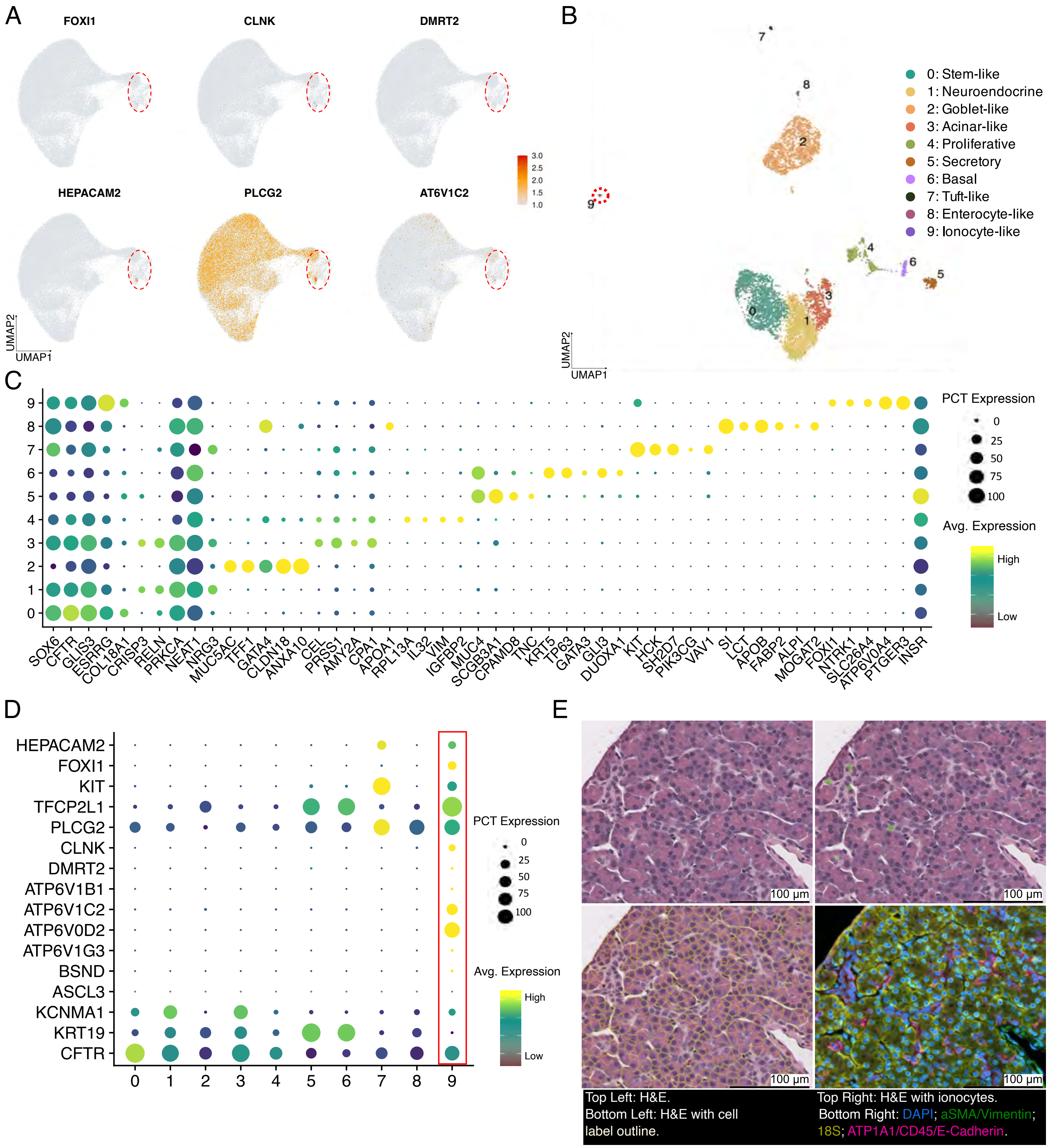

***Supplementary Fig. 4.***

**A.** UMAP of the ductal subset showing the gene expression of specific ionocyte gene markers.

**B.** UMAP of 5570 ductal cells obtained from the unsupervised clustering of the heterogeneous Ductal 6 and Ductal 8 substates in Supp. Fig. 2C, identifying a tuft-like pancreatic cell subset (cluster 7) and an ionocyte cell subset (cluster 9, highlighted).

**C.** Dotplot of identified marker genes for the clusters depicted in panel B.

**D.** Dotplot showing the relative gene expression of ionocyte-like markers in the clusters identified in panel B.

**E.** TME image of an adult snRNA core. Ionocytes have been highlighted in green in the corresponding panels. (Visualization generated with 10x Genomics Xenium Explorer 3.2.0)

### Supplementary Figure 5

A

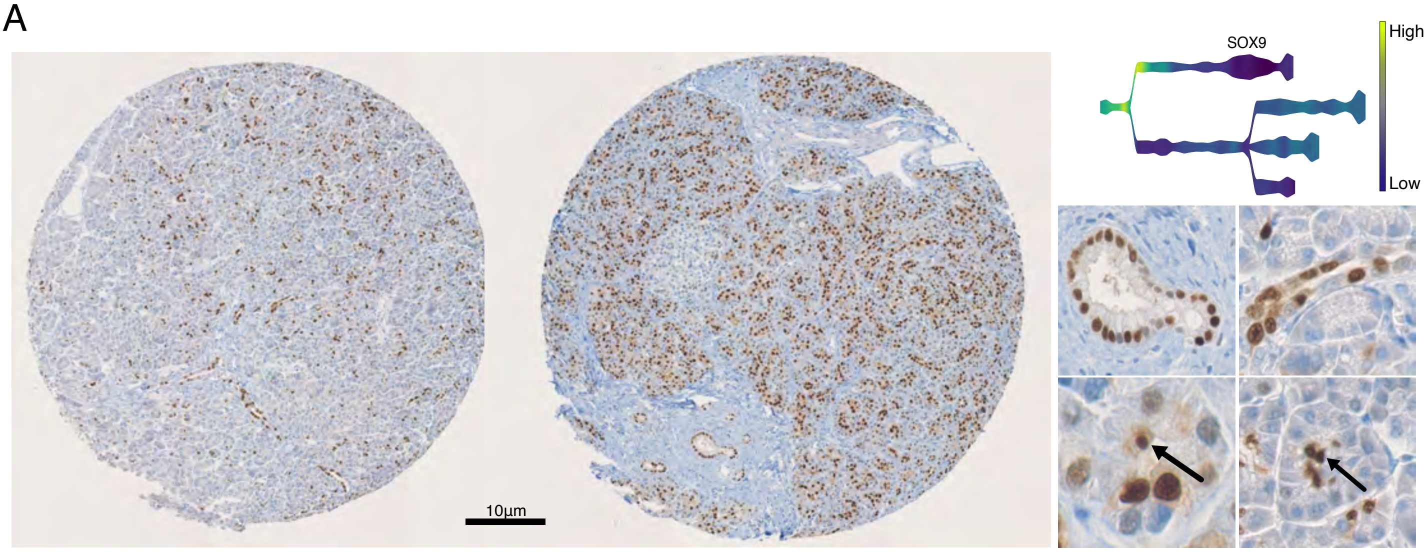


B

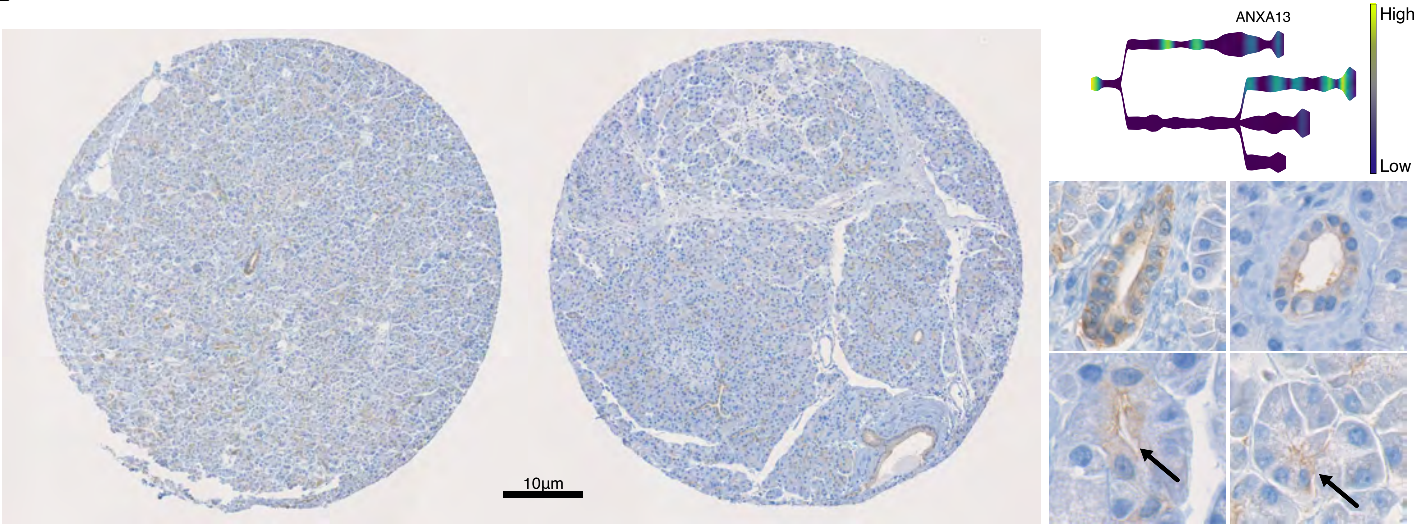


C

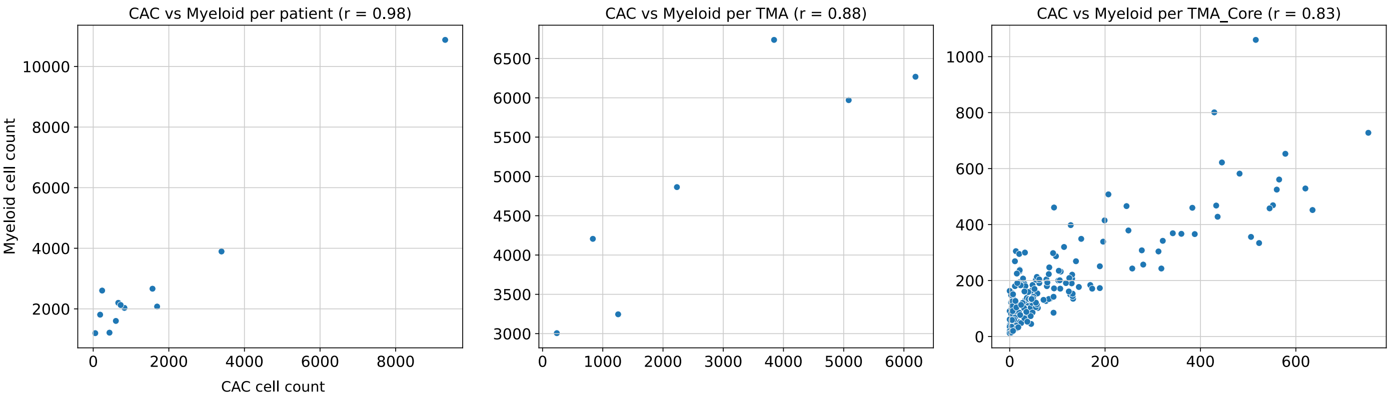

***Supplementary Fig. 5.***

**A-B.** *SOX9* and *ANXA13* immunostaining in TMA cores. Ductal cells were mainly stained although a high number of pCAC cells also were positively stained. Arrows point to putative pCACs. The adult snRNA STREAM trajectories of these two genes have been added.

**C.** CAC and Myeloid cell abundance correlation across different variables (TMA, TMA core and patient).

Supplementary Figure 6

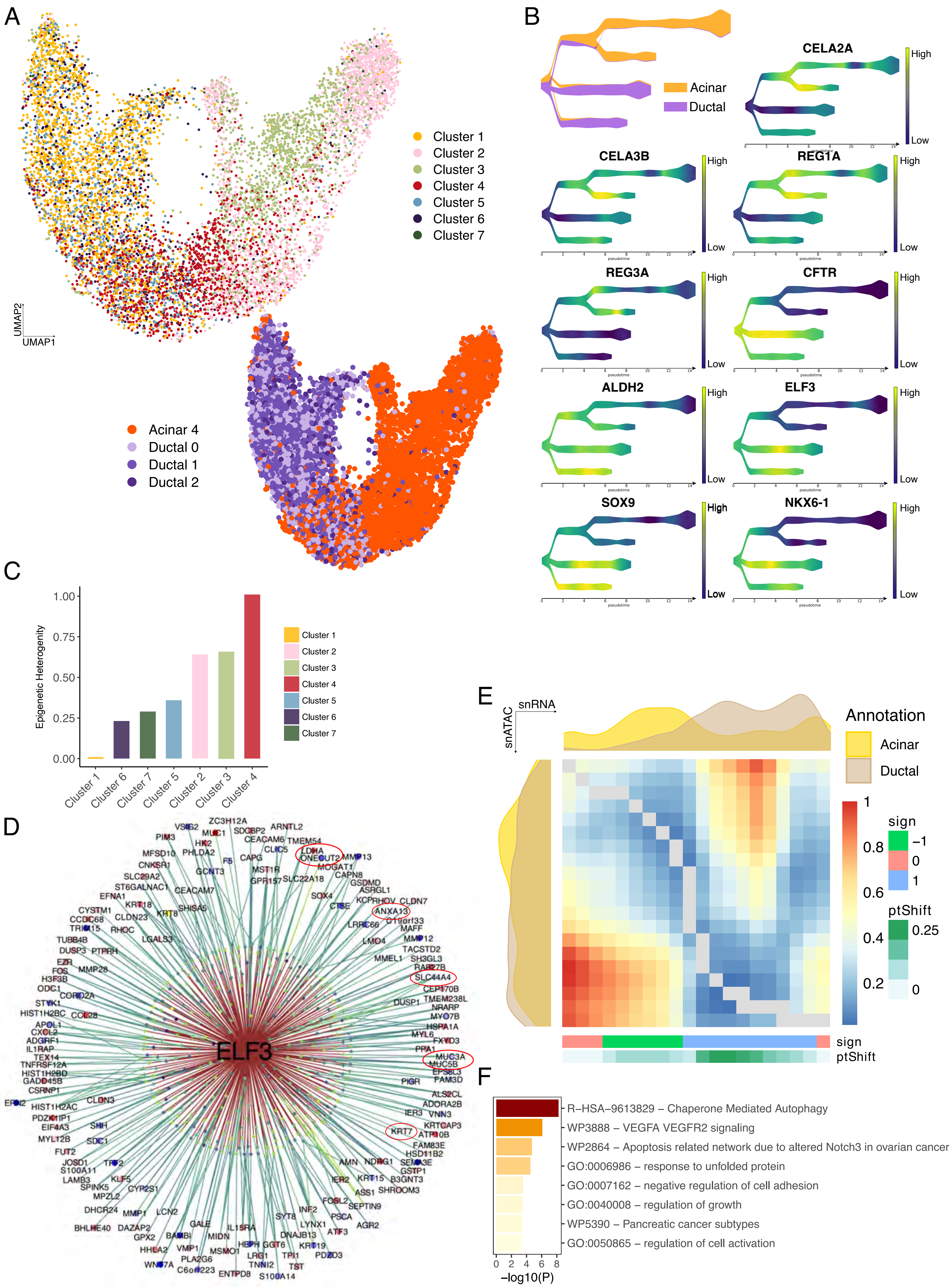

***Supplementary Fig. 6.***

**A.** UMAP visualization of the putative plastic cells identified in the adult snATAC dataset and their original annotation.

**B.** STREAM trajectory analysis of the putative plastic cells identified in adult snATAC. Both canonical acinar and ductal identity markers (REG3A - REG1A) as well plasticity markers (ELF3, SOX9 and NKX6-1) have been represented.

**C.** EpiCHAOS score for the putative plastics clusters identified in adult snATAC. The higher the score, the more epigenetic heterogeneity the cluster presents.

**D.** Pseudotime alignment of the snATAC and snRNA adult trajectory analyses through dynamic time warping algorithms. Acinar and ductal density plots have been added as marginal plots.

**E.** ELF3 regulon network as identified with SCENIC+ in a subset of 60k cells of the adult snATAC dataset. Diamond shapes represent regions and are colored according to the region log2FC in the acinar cells. Dots represent genes (labeled) and are also colored based on the log2FC of the gene expression in acinar cell.

**F.** Plasticity-associated terms identified through a GSEA in the snRNA population employed for the pseudotime alignment.

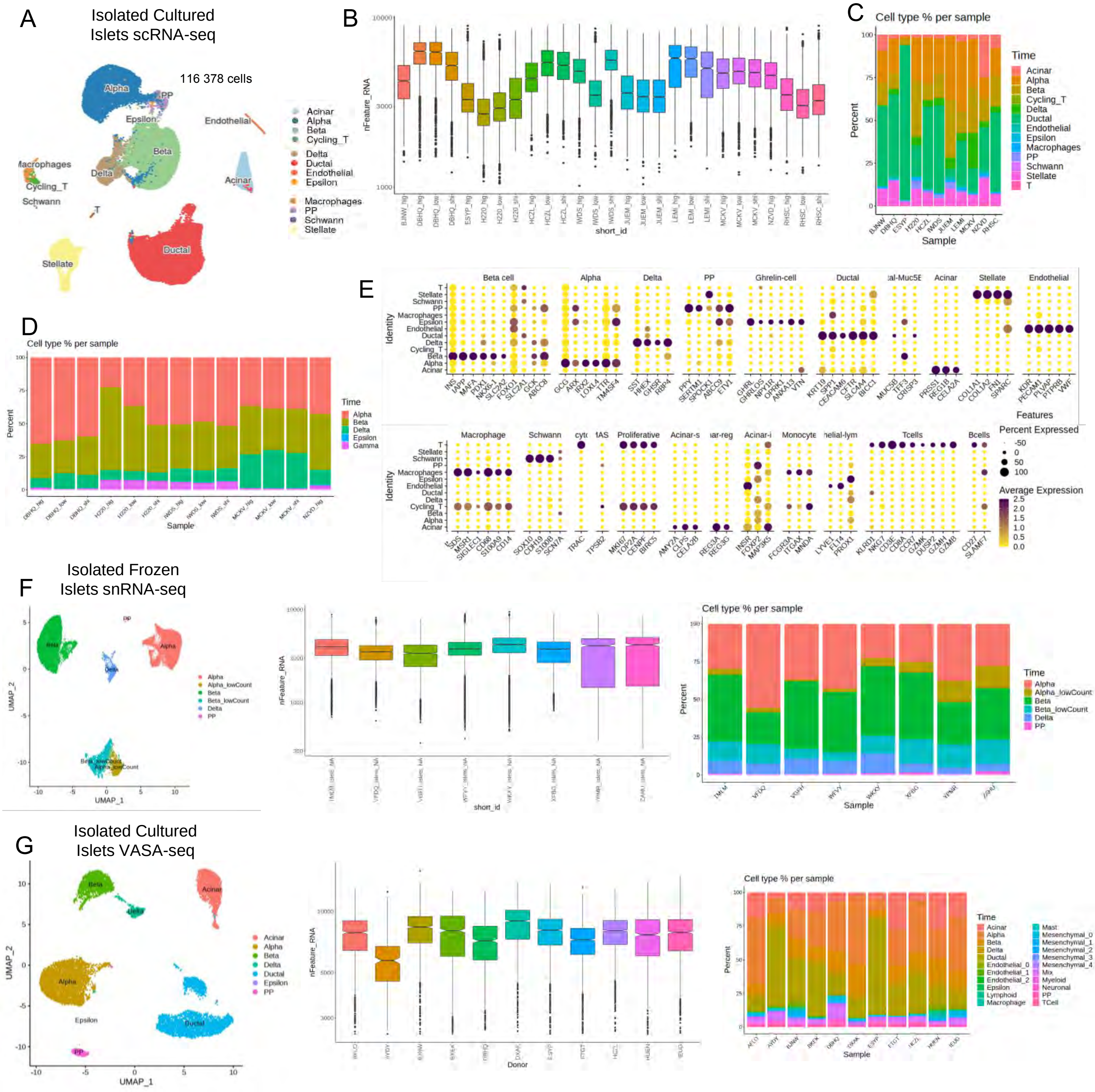

A
Isolated Cultured Islets scRNA-seq
116 378 cells
Alpha
PP
Epsilon
Endothelial
Beta
Macrophages
Delta
Cycling_T
Schwann
Acinar
T
Stellate
Ductal
Acinar
Alpha
Beta
Cycling_T
Delta
Ductal
Endothelial
Epsilon
Macrophages
PP
Schwann
Stellate
B
nFeature_RNA
short_id
C
Cell type % per sample
Percent
Sample
Time
Acinar
Alpha
Beta
Cycling_T
Delta
Ductal
Endothelial
Epsilon
Macrophages
PP
Schwann
Stellate
T
D
Cell type % per sample
Percent
Sample
Time
Alpha
Beta
Delta
Epsilon
Gamma
E
Identity
Features
Percent Expressed
Average Expression
F
Isolated Frozen Islets snRNA-seq
UMAP_2
UMAP_1
Alpha
Alpha_lowCount
Beta
Beta_lowCount
Delta
PP
nFeature_RNA
short_id
Cell type % per sample
Percent
Sample
Time
G
Isolated Cultured Islets VASA-seq
UMAP_2
UMAP_1
Acinar
Alpha
Beta
Delta
Ductal
Epsilon
PP
nFeature_RNA
Donor
Cell type % per sample
Percent
Sample
Time

***Supplementary Fig. 7.***

**A.** UMAP-base embedding projection of the isolated cultured islets scRNA-seq depicting cell clusters at cell-type resolution.
**B.** Number of genes detected in each individual scRNA-seq sample.
**C.** Distribution of cell types from each of the donors.
**D.** Distribution of endocrine cell types on each of the samples.
**E.** Gene markers for each of the cell types detected in A.
**F.** UMAP-base embedding projection of the isolated frozen islets endocrine cells snRNA-seq depicting cell clusters at cell-type resolution. Number of genes detected in each individual snRNA-seq encapsulation. Distribution of endocrine cell types on the samples from each of the 8 donors.
**G.** UMAP-base embedding projection of the isolated cultured islets VASA-seq depicting cell clusters at cell-type resolution. Number of genes detected in each individual VASA-seq sample. Distribution of endocrine cell types on the samples from each of the 11 donors.

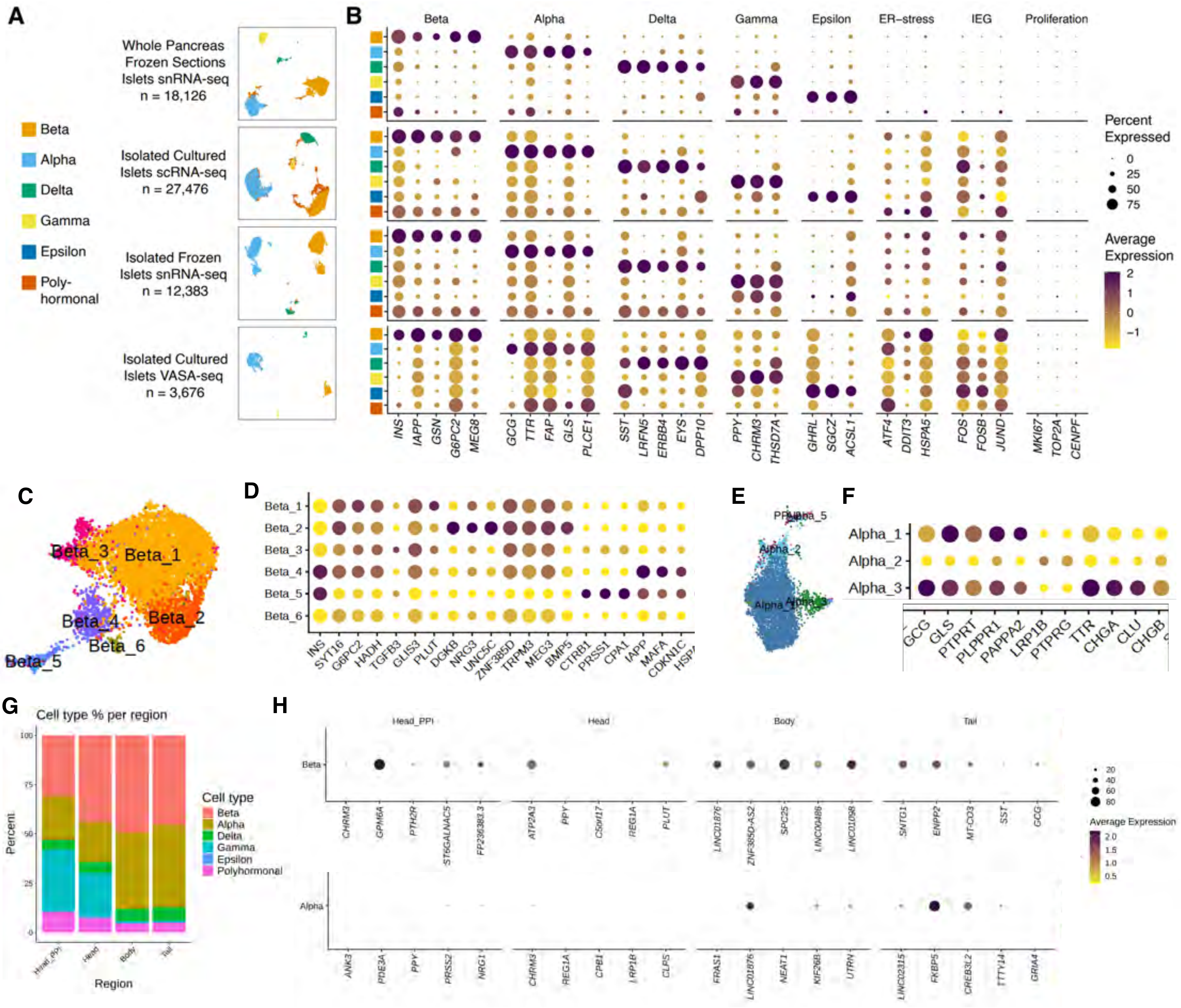
A
B
Whole Pancreas Frozen Sections Islets snRNA-seq n = 18,126
Isolated Cultured Islets scRNA-seq n = 27,476
Isolated Frozen Islets snRNA-seq n = 12,383
Isolated Cultured Islets VASA-seq n = 3,676
Beta
Alpha
Delta
Gamma
Epsilon
Poly-hormonal
Beta
Alpha
Delta
Gamma
Epsilon
ER−stress
IEG
Proliferation
INS
IAPP
GSN
G6PC2
MEG8
GCG
TTR
FAP
GLS
PLCE1
SST
LRFN5
ERBB4
EYS
DPP10
PPY
CHRM3
THSD7A
GHRL
SGCZ
ACSL1
ATF4
DDIT3
HSPA5
FOS
FOSB
JUND
MKI67
TOP2A
CENPF
Percent Expressed
0
25
50
75
Average Expression
2
1
0
−1
C
Beta_1
Beta_2
Beta_3
Beta_4
Beta_5
Beta_6
D
Beta_1
Beta_2
Beta_3
Beta_4
Beta_5
Beta_6
INS
SYT16
G6PC2
HADH
TGFB3
GLIS3
PLUT
DGKB
NRG3
UNC5C
ZNF385D
TRPM3
MEG3
BMP5
CTRB1
PRSS1
CPA1
IAPP
MAFA
CDKN1C
E
Alpha_2
Alpha_5
Alpha_1
Alpha_3
F
Alpha_1
Alpha_2
Alpha_3
GCG
GLS
PTPRT
PLPPR1
PAPPA2
LRP1B
PTPRG
TTR
CHGA
CLU
CHGB
G
Cell type % per region
Percent
100
75
50
25
0
Head_PPI
Head
Body
Tail
Region
Cell type
Beta
Alpha
Delta
Gamma
Epsilon
Polyhormonal
H
Head_PPI
Head
Body
Tail
Beta
CHRM3
GPM6A
PTH2R
ST6GALNAC5
FP236383.3
ATP2A3
PPY
C5orf17
REG1A
PLUT
LINC01876
ZNF385D-AS2
SPC25
LINC00486
LINC01098
SNTG1
ENPP2
MT-CO3
SST
GCG
Alpha
ANK3
PDE3A
PPY
PRSS2
NRG1
CHRM3
REG1A
CPB1
LRP1B
CLPS
FRAS1
LINC01876
NEAT1
KIF26B
UTRN
LINC02315
FKBP5
CREB3L2
TTTY14
GRIA4
20
40
60
80
Average Expression
2.0
1.5
1.0
0.5

***Supplementary Fig. 8.***

**A.** Human pancreatic islet endocrine cell datasets collected in ESPACE with different scRNA-seq modalities.

**B.** Consensus markers for endocrine cell types across the 4 scRNA-seq modalities.

**C.** UMAP-based embedding projection of the beta cell subpopulations observed in the whole pancreas frozen sections islets snRNA-seq dataset, depicting cell clusters at cell sub-type resolution.

**D.** Gene markers for each of the endocrine cell clusters in C.

**E.** UMAP-based embedding projection of the alpha cell subpopulations observed in the whole pancreas frozen sections islets snRNA-seq dataset, depicting cell clusters at cell sub-type resolution.

**F.** Gene markers for each of the beta cell clusters in E.

**G.** Distribution of the major islet endocrine cell types across the different pancreas anatomical regions (Head_PPI: uncinate process, Head, Body or Tail), as observed in the whole pancreas frozen sections islets snRNA-seq dataset.

**H.** Gene markers for beta and alpha cells originating from each pancreas anatomical region.

### Supp. Fig. 9 - Hormone expression outside of islets

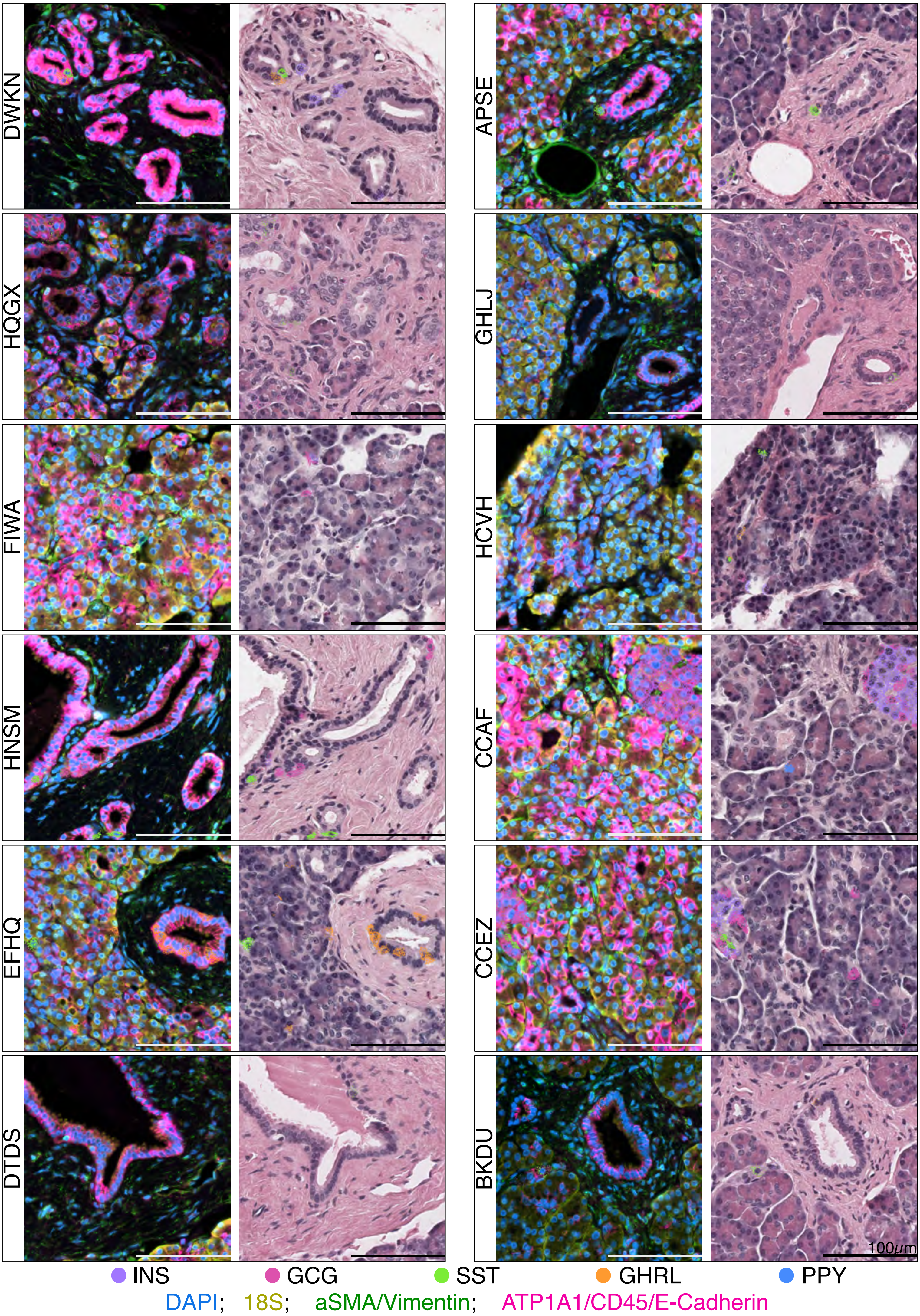

***Supplementary Fig. 9.***

**A.** Visualization of Xenium spatial transcriptomics data across different donors, depicting the presence of cells with endocrine hormone transcripts in ductal cells or outside pancreatic islets. Representative TMA cores containing endocrine cells, stained with hematoxilin-eosin and with immunofluorescence markers. Transcript counts for the main endocrine hormones are overlaid on the images. Immunofluorescence markers and endocrine hormones transcripts identities are color-coded and indicated in the legend at the bottom. (Visualization generated with 10x Genomics Xenium Explorer 3.2.0)

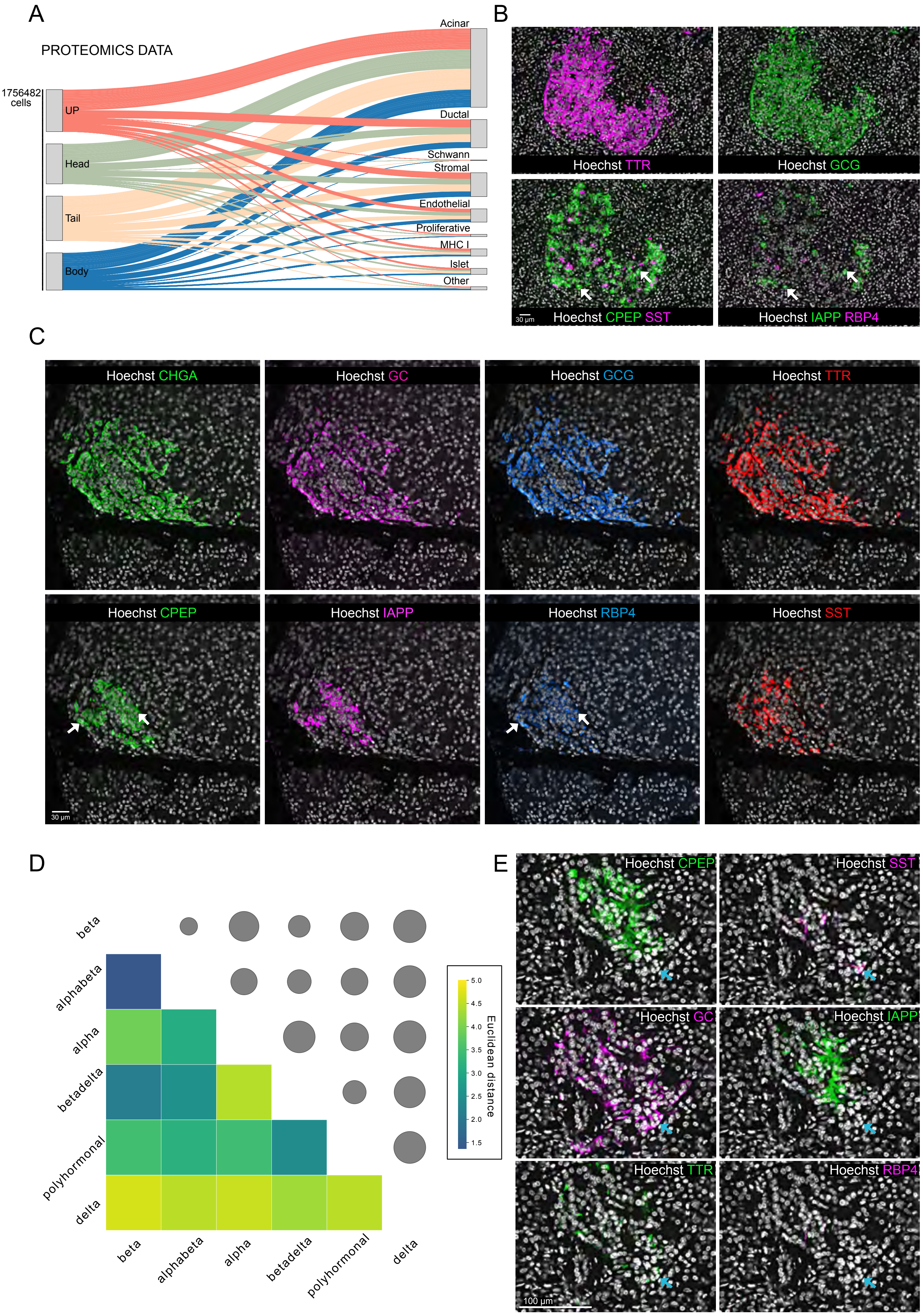

A
PROTEOMICS DATA
1756482 cells
UP
Head
Tail
Body
Acinar
Ductal
Schwann
Stromal
Endothelial
Proliferative
MHC I
Islet
Other
B
Hoechst TTR
Hoechst GCG
Hoechst CPEP SST
Hoechst IAPP RBP4
30 µm
C
Hoechst CHGA
Hoechst GC
Hoechst GCG
Hoechst TTR
Hoechst CPEP
Hoechst IAPP
Hoechst RBP4
Hoechst SST
30 µm
D
beta
alphabeta
alpha
betadelta
polyhormonal
delta
5.0
4.5
4.0
3.5
3.0
2.5
2.0
1.5
Euclidean distance
E
Hoechst CPEP
Hoechst SST
Hoechst GC
Hoechst IAPP
Hoechst TTR
Hoechst RBP4
100 µm

***Supplementary Fig. 10.***

**A.** A Sankey plot illustrating the proportion of annotated cell populations across various regions of non-diseased adult pancreatic tissue.

**B.** Representative multiplex immunofluorescence images of an adult pancreatic islet showing alpha cell markers (TTR, GCG), beta cell markers (CPEP, IAPP), and delta cell markers (SST, RBP4). Arrows indicate a classical beta cell population.

**C.** Representative immunostaining images displaying various islet subpopulations characterized by the expression of endocrine markers: chromogranin A (CHGA), glucagon (GCG), pepsinogen C (GC), transthyretin (TTR), C-peptide (CPEP), amylin (IAPP), retinol-binding protein 4 (RBP4), and somatostatin (SST) in pancreatic tissue sections. All markers are shown within the same islet. Arrows indicate cells co-expressing CPEP and RBP4.

**D.** A heatmap representing the average Euclidean distance between annotated endocrine cell populations, with the shortest distances to neighboring cells indicated in dark blue.

**E.** Example of hybrid islet states co-expressing alpha markers (GC, TTR) and a delta marker (SST) in the context of a whole islet (2D section). Images of remaining islet populations are shown for comparison. Arrows highlight hybrid polyhormonal cells positioned at the islet periphery, spatially separated from classical subtypes.

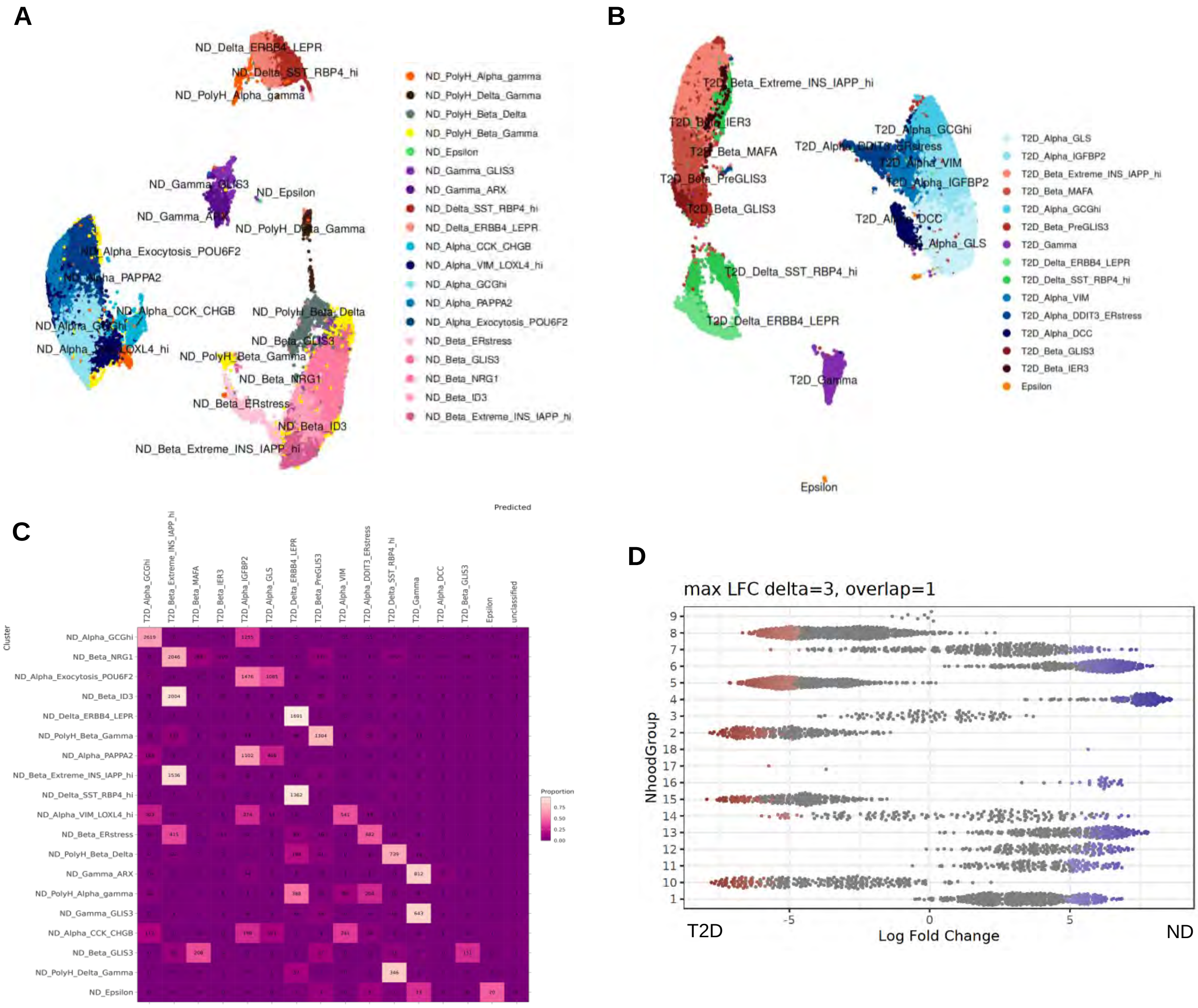
A
ND_PolyH_Alpha_gamma
ND_PolyH_Delta_Gamma
ND_PolyH_Beta_Delta
ND_PolyH_Beta_Gamma
ND_Epsilon
ND_Gamma_GLIS3
ND_Gamma_ARX
ND_Delta_SST_RBP4_hi
ND_Delta_ERBB4_LEPR
ND_Alpha_CCK_CHGB
ND_Alpha_VIM_LOXL4_hi
ND_Alpha_GCGhi
ND_Alpha_PAPPA2
ND_Alpha_Exocytosis_POU6F2
ND_Beta_ERstress
ND_Beta_GLIS3
ND_Beta_NRG1
ND_Beta_ID3
ND_Beta_Extreme_INS_IAPP_hi
B
T2D_Alpha_GLS
T2D_Alpha_IGFBP2
T2D_Beta_Extreme_INS_IAPP_hi
T2D_Beta_MAFA
T2D_Alpha_GCGhi
T2D_Beta_PreGLIS3
T2D_Gamma
T2D_Delta_ERBB4_LEPR
T2D_Delta_SST_RBP4_hi
T2D_Alpha_VIM
T2D_Alpha_DDIT3_ERstress
T2D_Alpha_DCC
T2D_Beta_GLIS3
T2D_Beta_IER3
Epsilon
C
Predicted
Cluster
Proportion
D
max LFC delta=3, overlap=1
NhoodGroup
T2D
Log Fold Change
ND

***Supplementary Fig. 11.***

**A**. UMAP-base embedding projection of the ND (non diabetes donor) isolated cultured islets endocrine cells scRNA-seq.

**B**. UMAP-base embedding projection of the T2D (type 2 diabetes donor) isolated cultured islets endocrine cells scRNA-seq.

**C**. Metrics of the scOMM-aided label transfer between ND and T2D scRNA-seq

**D**. Neighborhoods identified using differential abundance analysis between ND and T2D conditions.

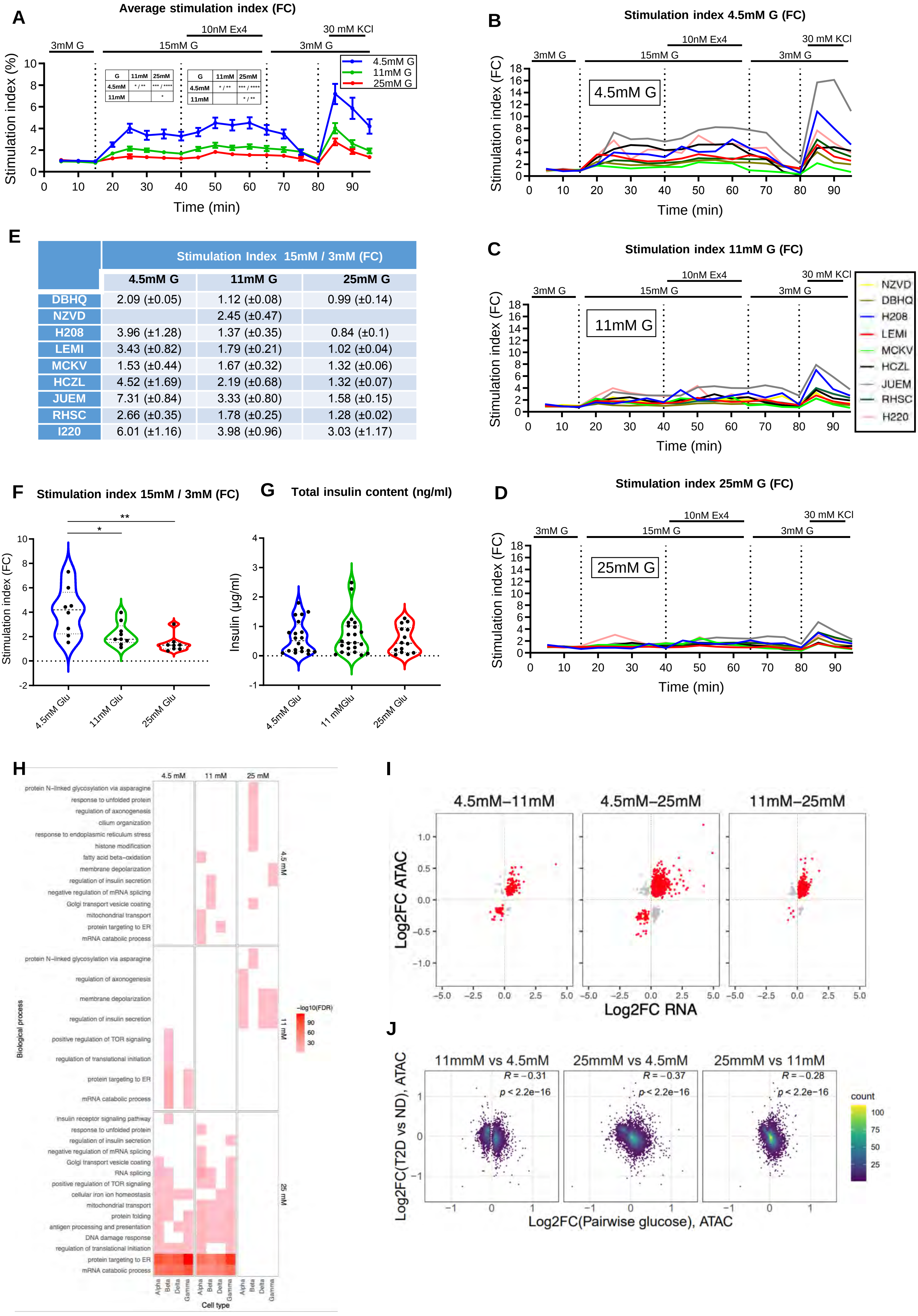


| | Stimulation Index 15mM / 3mM (FC) | | |
|---|---|---|---|
| | 4.5mM G | 11mM G | 25mM G |
| DBHQ | 2.09 (±0.05) | 1.12 (±0.08) | 0.99 (±0.14) |
| NZVD | | 2.45 (±0.47) | |
| H208 | 3.96 (±1.28) | 1.37 (±0.35) | 0.84 (±0.1) |
| LEMI | 3.43 (±0.82) | 1.79 (±0.21) | 1.02 (±0.04) |
| MCKV | 1.53 (±0.44) | 1.67 (±0.32) | 1.32 (±0.06) |
| HCZL | 4.52 (±1.69) | 2.19 (±0.68) | 1.32 (±0.07) |
| JUEM | 7.31 (±0.84) | 3.33 (±0.80) | 1.58 (±0.15) |
| RHSC | 2.66 (±0.35) | 1.78 (±0.25) | 1.28 (±0.02) |
| I220 | 6.01 (±1.16) | 3.98 (±0.96) | 3.03 (±1.17) |

***Supplementary Fig. 12.***

**A**. Average dynamic glucose stimulated insulin secretion of human islet preparations cultured for 72-96h at 4.5 mM, 11 mM or 25mM glucose showing glucose concentration-dependent insulin secretion impairment. n= 8-9 individual preparation run in triplicate (4.5 mM and 11 mM) or duplicate (25 mM).

**B-D**. Dynamic glucose stimulated insulin secretion of individual preparations of human islets cultured for 72-96h at 4.5 mM (B), 11 mM (C) and 25 mM (D) glucose. n= 8-9 preparations.

**E**. Summary table of Stimulation index values for each human islet preparation, derived from experiments presented in A.

**F.**Stimulation index (15 mM / 3mM glucose) of human islet preparations cultured for 72-96h in 4.5 mM, 11 mM, or 25 mM glucose, derived from experiments presented in A-D.

**G.** Total insulin content of samples presented in A-D.

Two-way ANOVA coupled to Bonferroni post-test was used in A-D and One-Way ANOVA coupled to Bonferroni post-test was used in E-F.

**H**. Biological processes enriched in the gene sets responsive to glucose in the different islet endocrine cell types.

**I**. Correlation between glucose regulated genes at RNA and ATAC level.

**J.** Correlation between glucose regulated genes and T2D regulated genes at ATAC level.

### Supplementary Figure 13 - Fetal cell populations

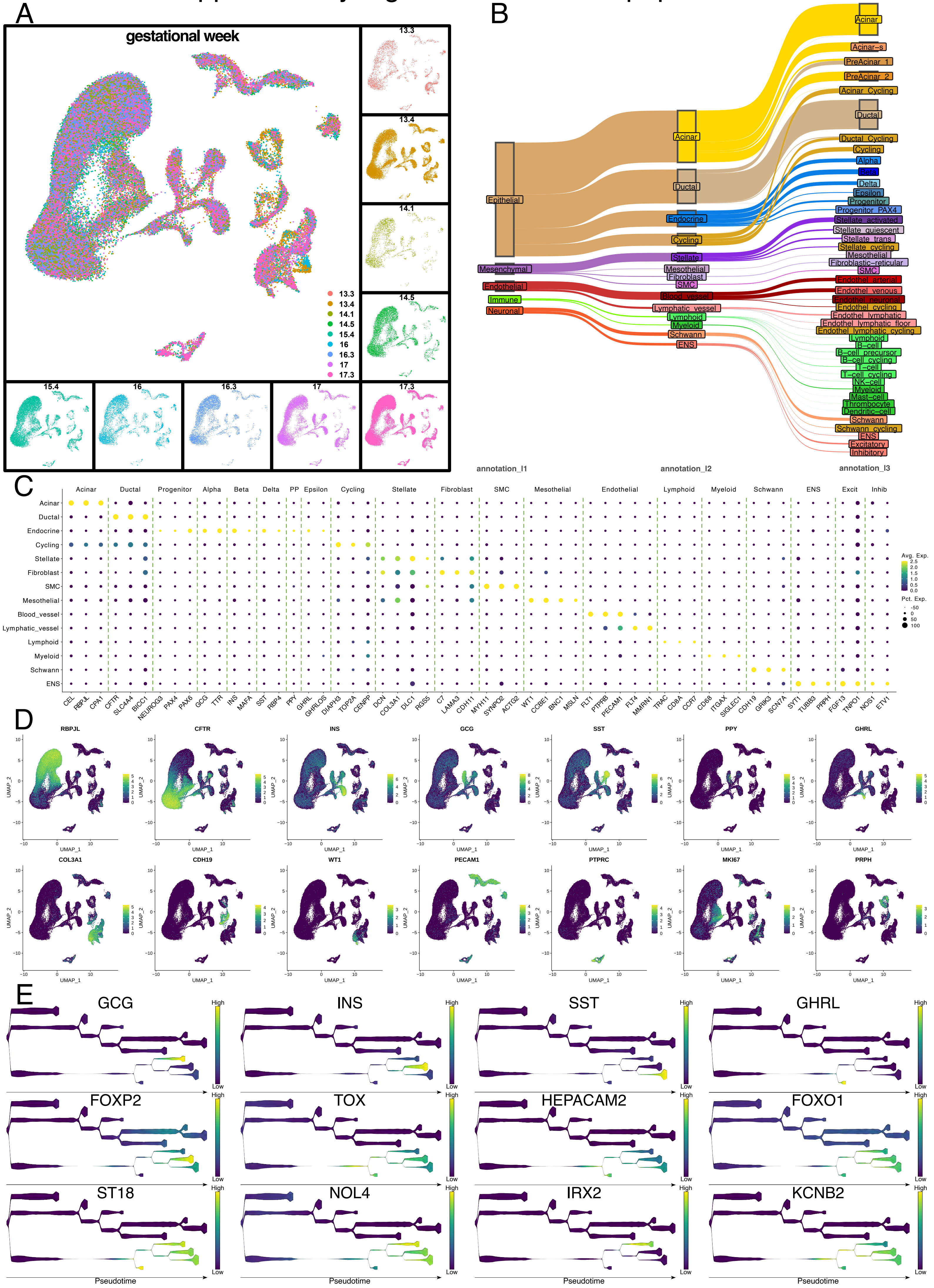

***Supplementary Fig. 13. - Fetal cell populations***

**A.** UMAP plot of 87324 nuclei from snRNA-seq from the fetal human pancreas, visualizing the weeks post conception (wpc) as well as the cell contribution from each donor.

**B.** Sankey plot visualizing the different granularities of cell (sub-)type annotation.

**C.** DotPlot of marker genes used to identify cell types.

**D.** Feature plot showing marker gene expression for each major compartment.

**E.** Stream plots visualizing expression over pseudotime. First row shows the expression of each major endocrine cell type (glucagon, insulin, somatostatin, ghrelin). Other rows show significant genes/transcription factors for endocrine differentiation identified by STREAM analysis

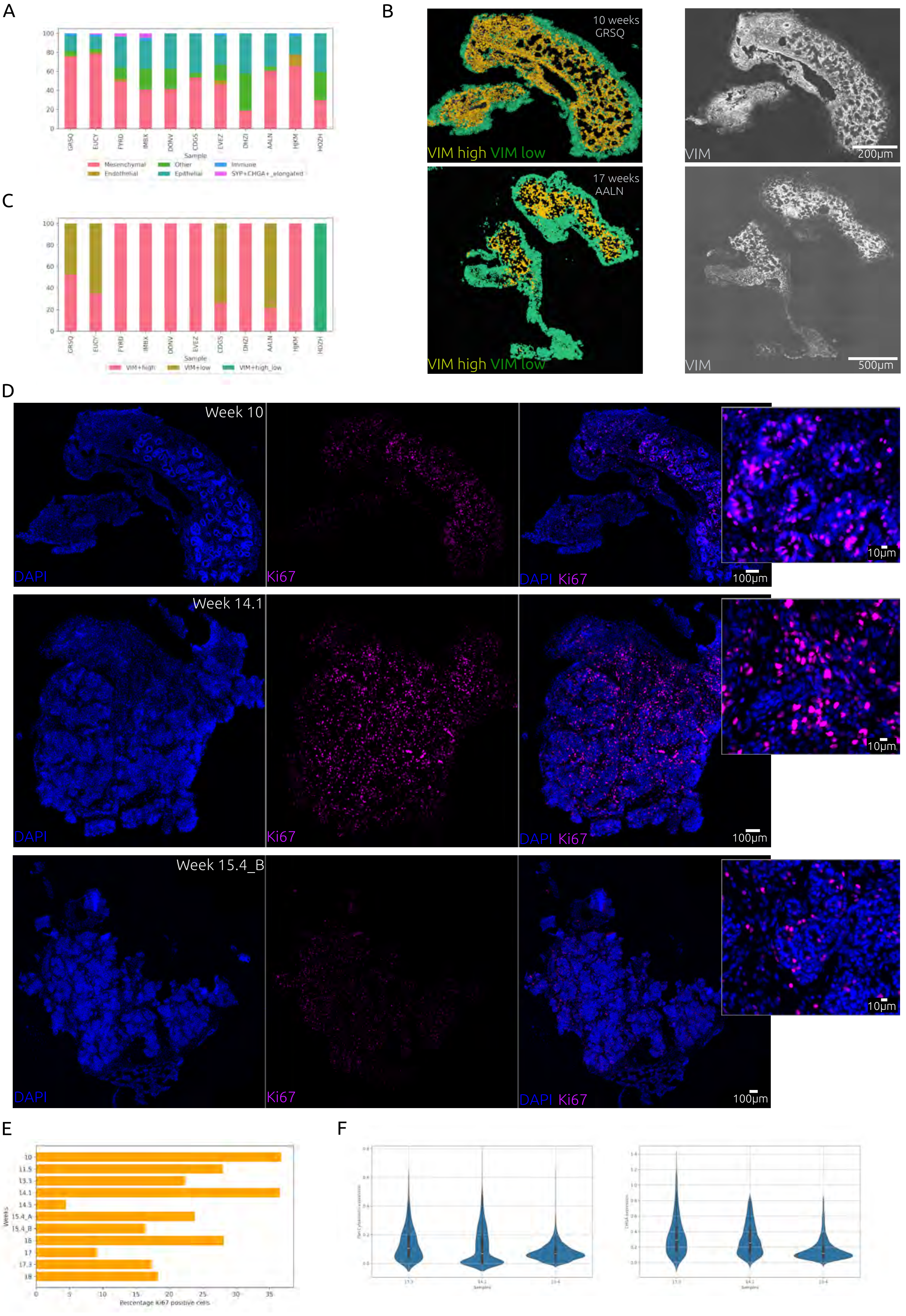

A
B
C
D
E
F
Sample
Mesenchymal
Endothelial
Other
Epithelial
Immune
SYP+CHGA+_elongated
GRSQ
EUCY
FYRD
IMBX
DDNV
CDGS
EVEZ
DHZI
AALN
HJKM
HOZH
VIM+high
VIM+low
VIM+high_low
10 weeks
GRSQ
17 weeks
AALN
VIM high VIM low
VIM
200µm
500µm
Week 10
Week 14.1
Week 15.4_B
DAPI
Ki67
DAPI Ki67
100µm
10µm
Weeks
Percentage Ki67 positive cells
10
11.5
13.3
14.1
14.5
15.4_A
15.4_B
16
17
17.3
18
Samples

***Supplementary Fig. 14. - Multiplexed imaging and spatial characterization of cell type populations in developing human pancreas tissue***

**A.** Barplot representing the percentage of annotated cell types in each sample, ranging from 10 to 18 weeks post conception (wpc).

**B.** Cell masks (left) showing the cluster with high VIM+ expressed cells (yellow) and low VIM+ expressed cells (green) in human pancreatic sections at 10 and 17 wpc. The grayscale images (right) from the same ages are immunofluorescence staining targeting VIM.

**C.** Barplot representing the percentage of high VIM+ expressed cells and low VIM+ expressed cells in each sample, ranging from 10 to 18 wpc.

**D.** Immunofluorescence images showing nuclear DAPI staining in blue, Ki67 expression in magenta, and overlay of both images at 10, 14.1 and 15.4 wpc.

**E.** Percentage of Ki67-positive cells in the developing human pancreas tissue.

**F.** Violin plots depicting the PanCytokeratin (left) and CHGA (right) protein expression distributions across gestational time points 13.3, 14.1 and 15.4 wpc. The white dots indicate the median intensity, black boxes indicate the interquartile range, the thin black lines indicate the upper and lower adjacent values.

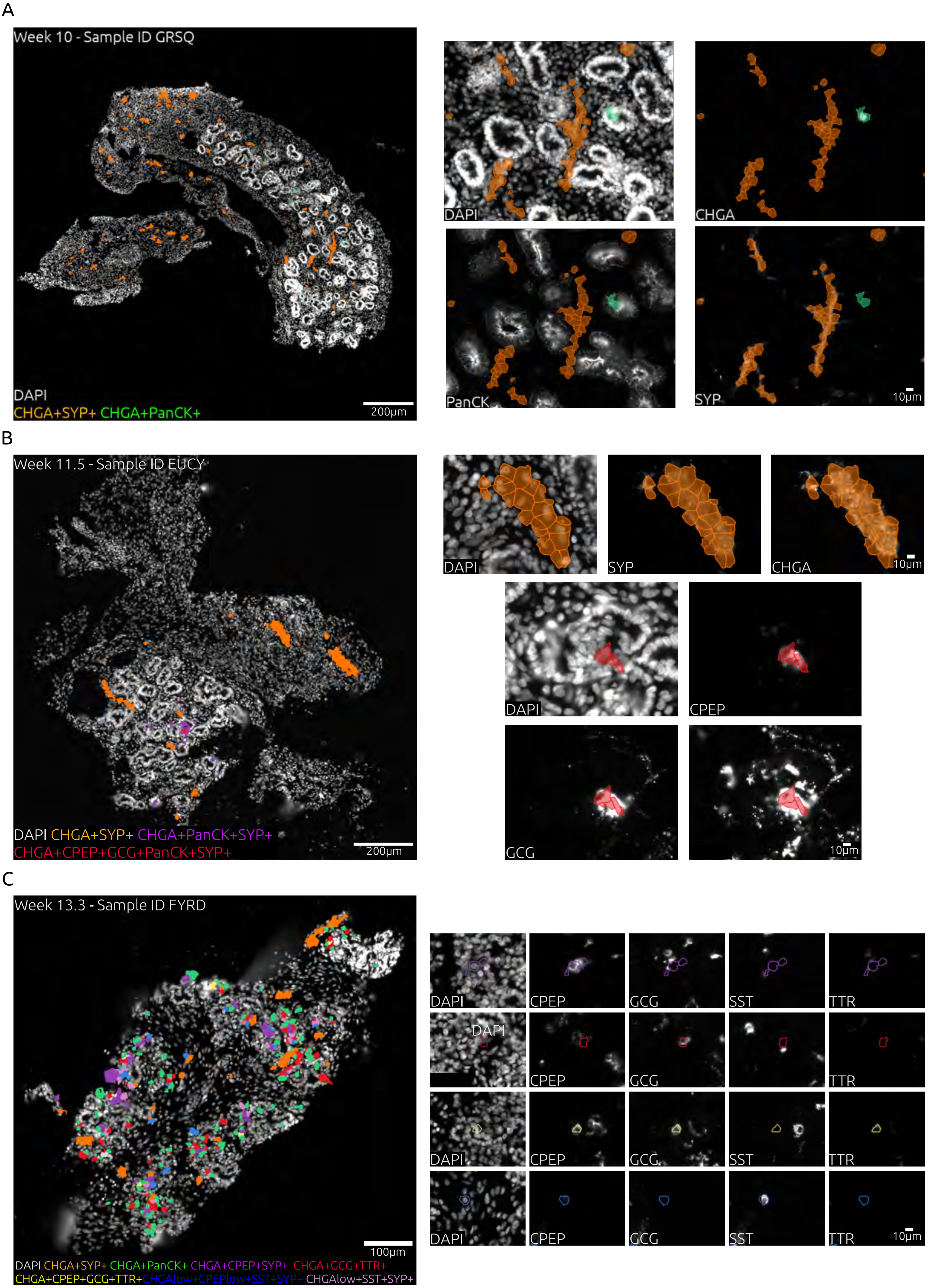
A
Week 10 - Sample ID GRSQ
DAPI
CHGA+SYP+ CHGA+PanCK+
200μm
DAPI
CHGA
PanCK
SYP
10μm
B
Week 11.5 - Sample ID EUCY
DAPI CHGA+SYP+ CHGA+PanCK+SYP+
CHGA+CPEP+GCG+PanCK+SYP+
200μm
DAPI
SYP
CHGA
10μm
DAPI
CPEP
GCG
10μm
C
Week 13.3 - Sample ID FYRD
100μm
DAPI CHGA+SYP+ CHGA+PanCK+ CHGA+CPEP+SYP+ CHGA+GCG+TTR+
CHGA+CPEP+GCG+TTR+ CHGAlow+CPEPlow+SST+SYP+ CHGAlow+SST+SYP+
DAPI
CPEP
GCG
SST
TTR
10μm

***Supplementary Fig. 15. - Multiplexed imaging of endocrine populations during human pancreas development from 10 to 13.5 weeks post conception (wpc), accompanied by representative images at higher magnifications***

**A.** Immunofluorescence staining of a 10 wpc human pancreatic tissue section, showing the spatial organization of CHGA+SYP+ cells (orange cell mask) and CHGA+PanCK+ (green cell mask) cells.

**B.** Immunofluorescence staining of an 11.5 wpc human pancreatic tissue section, displaying the expression of the first CPEP+GCG+ cells (red cell mask).

**C.** Immunofluorescence staining of a 13.3 wpc human pancreatic tissue section, demonstrating the formation of individual endocrine cells identified through unsupervised clustering.

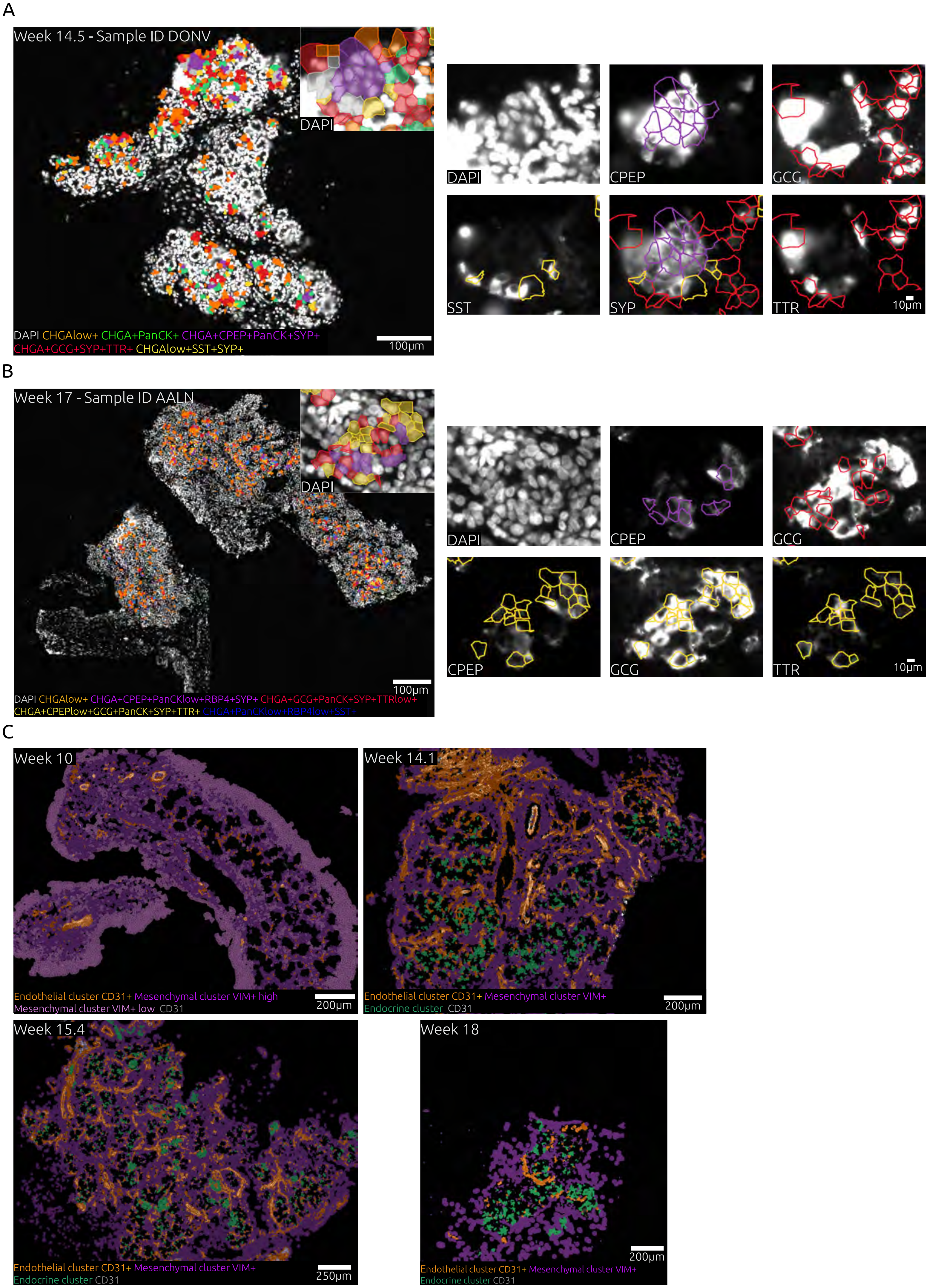

A
Week 14.5 - Sample ID DONV
DAPI
DAPI CHGAlow+ CHGA+PanCK+ CHGA+CPEP+PanCK+SYP+
CHGA+GCG+SYP+TTR+ CHGAlow+SST+SYP+
100μm
DAPI
CPEP
GCG
SST
SYP
TTR
10μm
B
Week 17 - Sample ID AALN
DAPI
DAPI CHGAlow+ CHGA+CPEP+PanCKlow+RBP4+SYP+ CHGA+GCG+PanCK+SYP+TTRlow+
CHGA+CPEPlow+GCG+PanCK+SYP+TTR+ CHGA+PanCKlow+RBP4low+SST+
100μm
DAPI
CPEP
GCG
CPEP
GCG
TTR
10μm
C
Week 10
Endothelial cluster CD31+ Mesenchymal cluster VIM+ high
Mesenchymal cluster VIM+ low CD31
200μm
Week 14.1
Endothelial cluster CD31+ Mesenchymal cluster VIM+
Endocrine cluster CD31
200μm
Week 15.4
Endothelial cluster CD31+ Mesenchymal cluster VIM+
Endocrine cluster CD31
250μm
Week 18
Endothelial cluster CD31+ Mesenchymal cluster VIM+
Endocrine cluster CD31
200μm

***Supplementary Fig. 16. - Multiplexed imaging and spatial characterization of endocrine populations and vascularization in developing human pancreas tissue***

**A.** Immunofluorescence staining of a 14.5 weeks post conception (wpc) human pancreatic tissue section, showing the spatial distribution of beta cells (purple cell mask), alpha cells (red cell mask) and delta cells (yellow cell mask), accompanied by representative images at higher magnifications.

**B.** Immunofluorescence staining of a 17 wpc human pancreatic tissue section, revealing the spatial locations of beta cells (purple cell mask), alpha cells (red cell mask), alpha-beta cells (yellow cell mask) and, delta cells (blue cell mask), accompanied by representative images at higher magnifications.

**C.** Immunofluorescence staining showing the progressive vascularization during the fetal development. CD31 signal (grey color) overlapping with cell masks corresponding to endothelial cells (orange), mesenchymal cells (magenta) and endocrine cells (green) of 10, 14.1, 15.4 and 18 weeks post conception.

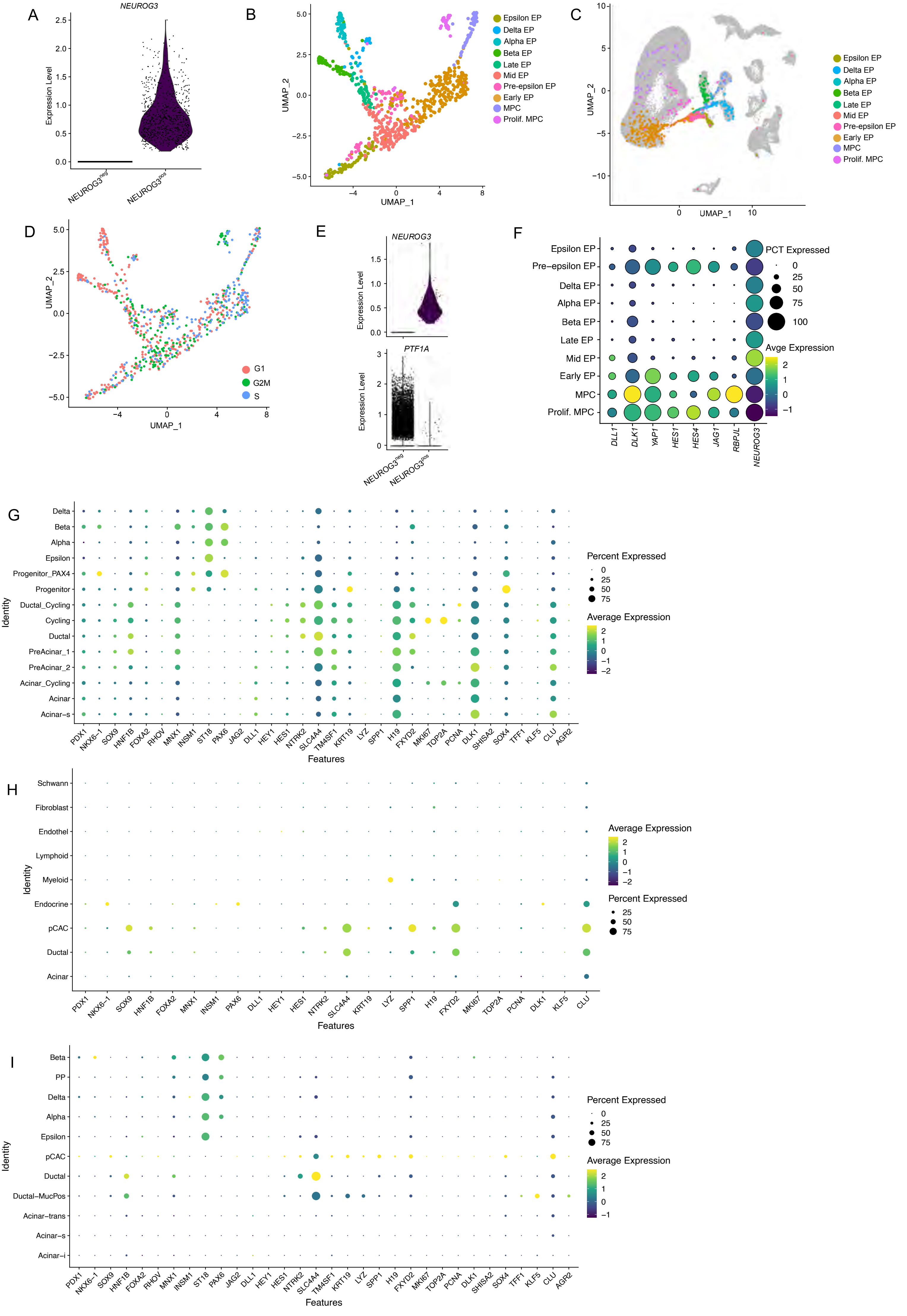
A
NEUROG3
Expression Level
NEUROG3neg
NEUROG3pos
B
UMAP_1
UMAP_2
Epsilon EP
Delta EP
Alpha EP
Beta EP
Late EP
Mid EP
Pre-epsilon EP
Early EP
MPC
Prolif. MPC
C
UMAP_1
UMAP_2
Epsilon EP
Delta EP
Alpha EP
Beta EP
Late EP
Mid EP
Pre-epsilon EP
Early EP
MPC
Prolif. MPC
D
UMAP_1
UMAP_2
G1
G2M
S
E
NEUROG3
PTF1A
Expression Level
NEUROG3neg
NEUROG3pos
F
Epsilon EP
Pre−epsilon EP
Delta EP
Alpha EP
Beta EP
Late EP
Mid EP
Early EP
MPC
Prolif. MPC
DLL1
DLK1
YAP1
HES1
HES4
JAG1
RBPJL
NEUROG3
PCT Expressed
Avge Expression
G
Delta
Beta
Alpha
Epsilon
Progenitor_PAX4
Progenitor
Ductal_Cycling
Cycling
Ductal
PreAcinar_1
PreAcinar_2
Acinar_Cycling
Acinar
Acinar−s
Identity
PDX1 NKX6−1 SOX9 HNF1B FOXA2 RHOV MNX1 INSM1 ST18 PAX6 JAG2 DLL1 HEY1 HES1 NTRK2 SLC4A4 TM4SF1 KRT19 LYZ SPP1 H19 FXYD2 MKI67 TOP2A PCNA DLK1 SHISA2 SOX4 TFF1 KLF5 CLU AGR2
Features
Percent Expressed
Average Expression
H
Schwann
Fibroblast
Endothel
Lymphoid
Myeloid
Endocrine
pCAC
Ductal
Acinar
Identity
PDX1 NKX6−1 SOX9 HNF1B FOXA2 MNX1 INSM1 PAX6 DLL1 HEY1 HES1 NTRK2 SLC4A4 KRT19 LYZ SPP1 H19 FXYD2 MKI67 TOP2A PCNA DLK1 KLF5 CLU
Features
Average Expression
Percent Expressed
I
Beta
PP
Delta
Alpha
Epsilon
pCAC
Ductal
Ductal−MucPos
Acinar−trans
Acinar−s
Acinar−i
Identity
PDX1 NKX6−1 SOX9 HNF1B FOXA2 RHOV MNX1 INSM1 ST18 PAX6 JAG2 DLL1 HEY1 HES1 NTRK2 SLC4A4 TM4SF1 KRT19 LYZ SPP1 H19 FXYD2 MKI67 TOP2A PCNA DLK1 SHISA2 SOX4 TFF1 KLF5 CLU AGR2
Features
Percent Expressed
Average Expression

***Supplementary Fig. 17. - Analysis of NEUROG3+ fetal subpopulations***

**A.** Violin plot indicating NEUROG3 expression levels in NEUROG3 negative and positive cells.

**B.** UMAP of reclustered NEUROG3 positive cells.

**C**. Allocation on a UMAP of reclustered NEUROG3 positive cells on all detected cell types in the fetal dataset.

**D.** Cell cycle analysis illustrated in UMAP space.

**E.** Violin plots comparing NEUROG3 and PTF1A expression levels in NEUROG3 negative and positive cells selected from acinar cells.

**F.** Dotplot representing gene markers for NOTCH signaling in selected NEUROG3 positive cell clusters.

**G.** Dotplot representing multipotent pancreatic progenitor markers in the fetal dataset.

**H.** Dotplot representing multipotent pancreatic progenitor markers in the adult dataset.

**I.** Dotplot representing multipotent pancreatic progenitor markers in the Xenium dataset

### Supplementary Figure 18 - Adult and fetal comparison

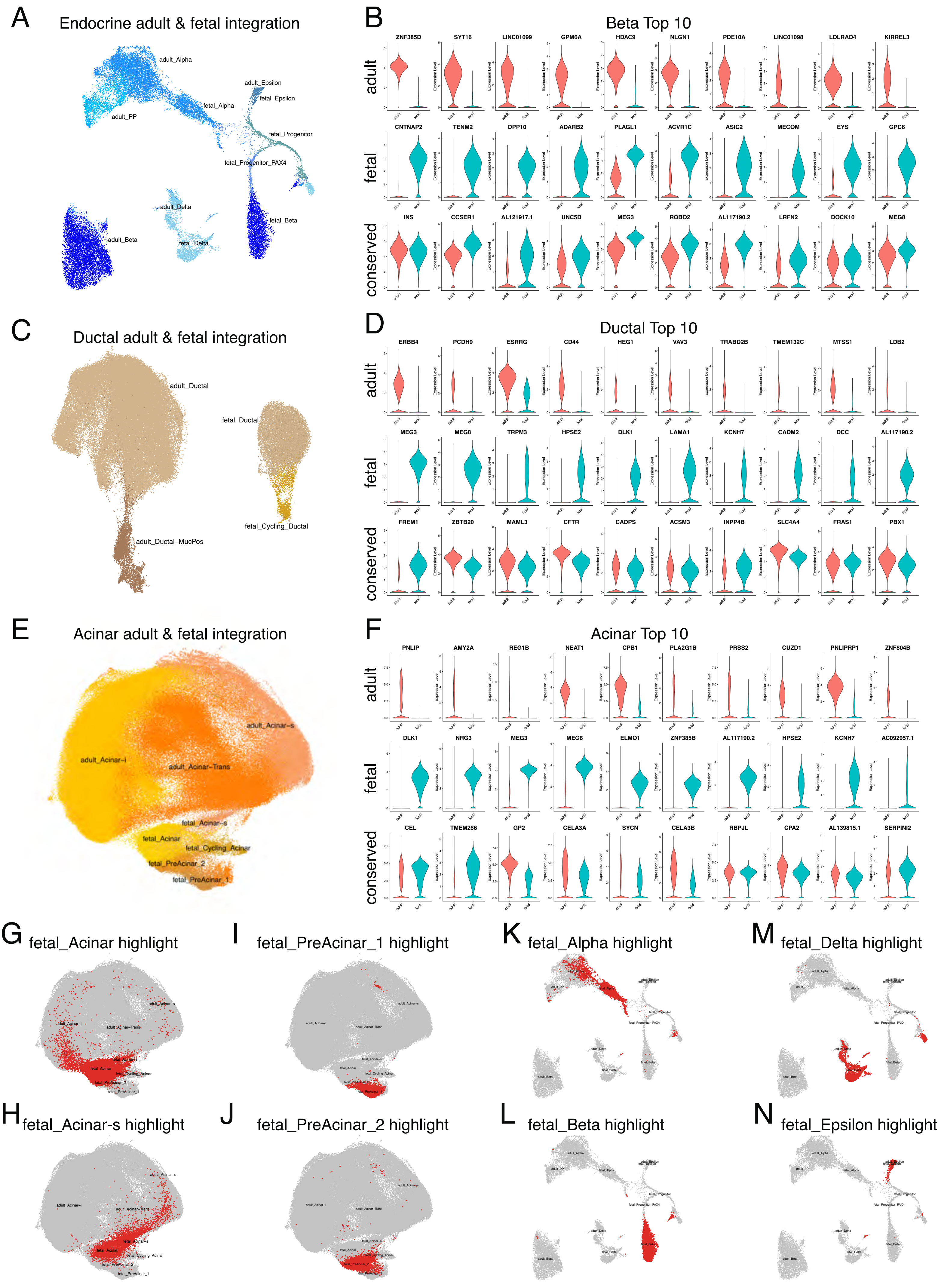

***Supplementary Fig. 18. - Adult and fetal comparison***

**A.** UMAP visualization of the integrated endocrine subset.

**B.** Violin plot comparing expression of indicated genes in adult and fetal subset of the beta cell compartment in the integrated endocrine object. First row are top 10 differentially expressed genes in the adult subset; second row are top 10 differentially expressed genes in the fetal subset; third row are top 10 conserved genes between adult and fetal.

**C.** UMAP visualization of the integrated ductal subset

**D.** Violin plot comparing expression of indicated genes in adult and fetal subset of the ductal cell compartment in the integrated ductal object. First row are top 10 differentially expressed genes in the adult subset; second row are top 10 differentially expressed genes in the fetal subset; third row are top 10 conserved genes between adult and fetal.

**E.** UMAP visualization of the integrated acinar subset

**F.** Violin plot comparing expression of indicated genes in adult and fetal subset of the acinar cell compartment in the integrated acinar object. First row are top 10 differentially expressed genes in the adult subset; second row are top 10 differentially expressed genes in the fetal subset; third row are top 10 conserved genes between adult and fetal.

**G-J.** Integrated UMAP plot of 353260 nulcei from snRNA-seq from adult and fetal human acinar subsets showing the fetal sub-populations Acinar (**G**), Acinar-s (**H**), PreAcinar_1 (**I**) and PreAcinar_2 (**J**) in red.

**K-N.** Integrated UMAP plot of 27847 nulcei from snRNA-seq from adult and fetal human endocrine subsets showing the fetal sub-populations Alpha (**K**), Beta (**L**), Delta (**M**) and Epsilon (**N**) in red.

### Supplementary Figure 19 - GO analysis of human fetal Pancreas

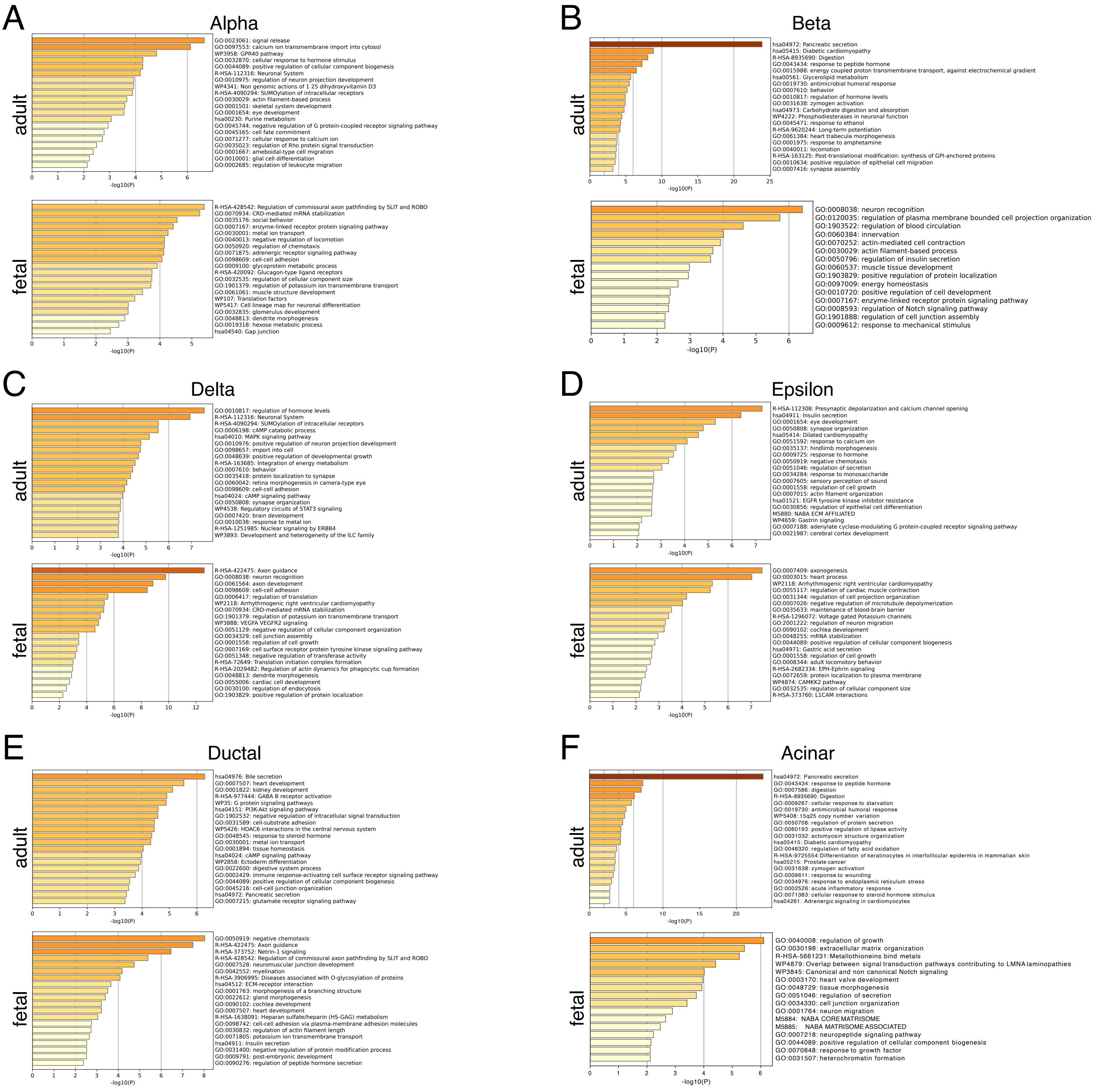

***Supplementary Fig. 19. - GO-term analysis for human fetal pancreas***

**A-F:** GO-term analysis using Metascape based on the top 100 differentially expressed genes of adult and fetal subsets in the Alpha (**A**), Beta (**B**), Delta (**C**), Epsilon (**D**), Ductal (**E**) and Acinar (**F**) compartment based on the integrated objects.

### Supplementary Figure 20 - Conserved genes in human fetal pancreas

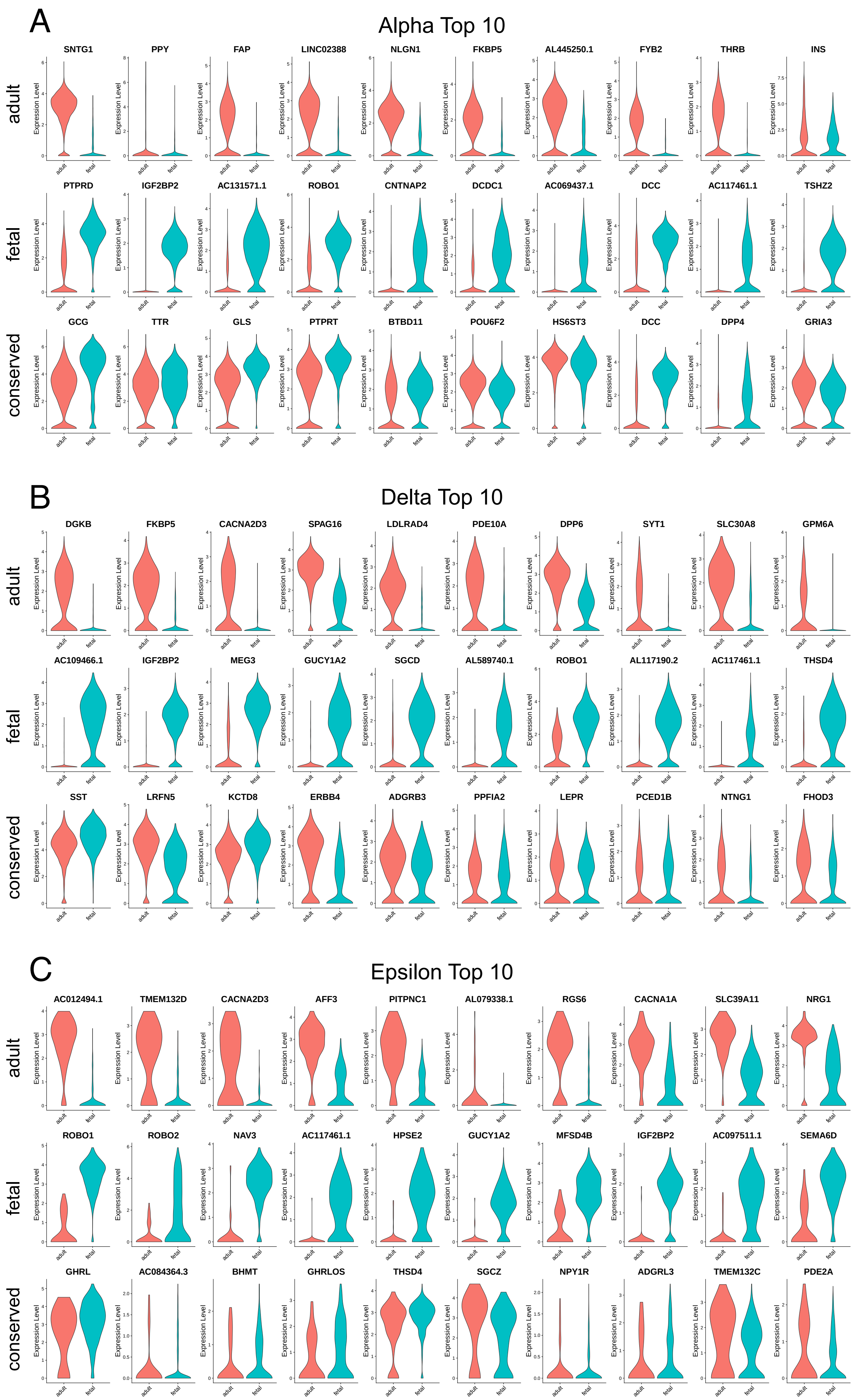

***Supplementary Fig. 20. - Endocrine human adult and fetal gene expression comparison***

**A.** Violin plot comparing expression of indicated genes in adult and fetal subset of the alpha cell compartment in the integrated endocrine object. First row are top 10 differentially expressed genes in the adult subset; second row are top 10 differentially expressed genes in the fetal subset; third row are top 10 conserved genes between adult and fetal.

**B.** Violin plot comparing expression of indicated genes in adult and fetal subset of the delta cell compartment in the integrated endocrine object. First row are top 10 differentially expressed genes in the adult subset; second row are top 10 differentially expressed genes in the fetal subset; third row are top 10 conserved genes between adult and fetal.

**C.** Violin plot comparing expression of indicated genes in adult and fetal subset of the epsilon cell compartment in the integrated endocrine object. First row are top 10 differentially expressed genes in the adult subset; second row are top 10 differentially expressed genes in the fetal subset; third row are top 10 conserved genes between adult and fetal.

**F.** Violin plot showing the number of UMIs per sample.

**G.** Violin plot showing the number of genes per sample.

**H.** Bar plot showing the contribution to each major cell type from each sample

#### Supplementary Notes

##### *Supp. note 1: Transcriptional heterogeneity of ductal cells*

Through the analysis of ductal cells, we have identified two main subtypes: *CFTR+* and *MUC+* cells (referred to as *Ductal* and *Ductal-MucPos*, respectively, in Fig. 1D). The majority of ductal cells (~93%) belong to the *CFTR+* subtype (D0-D5, D7, D9 clusters, Supp. Fig. 2F), characterized by the expression of genes associated with various important functions of ductal cells. These include ion transport (*SLC4A4, SLC17A4*), metabolic regulation (*FOXO1, PDE3A*), and mitochondrial activity (*MT-CO2, MT-CO1, MT-ATP6*), indicating active metabolism essential for ductal function. Additionally, transcriptional regulation by genes such as *PRDM16*, and *CREB3L1* underscores the versatile roles these cells play in maintaining ductal cell functions and cellular homeostasis.

Interestingly, within the *CFTR+* cells, the cell cluster D4 displayed high expression of genes that are specifically associated with digestive functions typically related to acinar cells in the pancreas (e.g. *AMY2A, CPA1, and CEL3A, Supp. Fig. 2F*). The expression of such a wide array of markers indicated the potential functional versatility and plasticity of these ductal acinar-like cells.

In contrast, the second subtype of ductal cells (clusters D6 and D8) exhibits a distinct gene expression profile. These cells are characterized by the upregulation of genes associated with secretory processes (*MUC5B, GOLM1, PRKCA, PGC, PLD1*), ion channel activity (*TMC5, TRIM5*), as well as genes involved in cell morphology and tissue morphogenesis (*SHROOM3, THSD4, WDR72*). Notably, the high expression of the transcription factor gene *ONECUT2* in the *KRT19+*, *MUC+* cells suggests a significant role in the regulation and maintenance of ductal cell identity and function.

Furthermore, within these cells, we identified a very rare but distinct subpopulation of approximately 21 cells expressing a signature of ionocyte-like markers, including *HEPACAM2*, *FOXI1*, *KIT*, *TFCP2L1*, *PLCG2*, *CLNK*, *ATP6V0D2*, and *ATP6V1C2* (Supp. Fig. 4A–D). Ionocytes are specialized epithelial cells involved in ion transport and homeostasis, and have recently been described in lung and other tissues using similar marker combinations (*117, 118*). Although rare, these ionocyte-like cells were also detected in our spatial transcriptomic (Xenium) dataset using the snRNA-seq–derived gene signature (see Methods), enabling further exploration of their localization and potential function in the pancreas (Supp. Fig. 4E)).

***Supp. note 2: Transcriptional heterogeneity of acinar cells***

The analysis of the acinar compartment revealed 12 clusters (Supp. Fig. 2B), highlighting the complexity of these cells beyond their well-known role in enzyme production, secretion, and digestion.

Acinar-s cells, characterized by the expression of genes such as *CPA1*, *PRSS1*, and the *CEL* genes (*CELA2B*, *CELA3A*, *CELA3B*), align predominantly with clusters A4, A6, A8, A9 and A11 (Supp. Fig. 2E). These clusters are marked by high expression of genes responsible for the synthesis and secretion of digestive enzymes, including *CLPS*, *AMY2A*, *PNLIP*, *CTRC*, *CPA1* and *PRSS3*.

In contrast, Acinar-i cells, which express genes involved in regulatory functions and stress responses (e.g., *RBPJL*, *CHRM3*, *FOXP2*), are associated with clusters A0, A1 and A5. These clusters include cells marked by genes such as *TPM1*, *FOXO1*, *FOXO3*, and other markers like *CD44*, *C3* and *SOD2*, indicating a potential role in cellular stress response and metabolic regulation.

Acinar-trans cells exhibit a mixed expression profile, indicating a transitional state, and overlap with clusters A2, A3, A7, and A10. These clusters show combined features of digestive enzyme synthesis (e.g., *PRSS2*, *CPA1*, *CTRB2*) and regulatory and mitochondrial functions (e.g., *CFTR*, *BICC1*, *SLC4A4*, *PDK4*, *BNIP3, MT-CYB, MT-ND1*), reflecting functional plasticity. Notably, the expression of *CFTR*, *BICC1*, and *SLC4A4*, genes typically associated with ductal cells, suggests that these acinar-trans cells might be in a process of transdifferentiation from acinar to ductal cells. This dual expression pattern supports the hypothesis that acinar-trans cells possess the ability to shift their phenotype, demonstrating a capacity for significant functional adaptation within the tissue.

Moreover, these cells express REG genes in a gradient, with clusters 3 and 7 showing the highest expression, indicating an underlying regenerative process (Supp. Fig. 2E). Although REG+ cells have been previously associated with a specific acinar state (*13*), here we observe that REG expression is not confined to a specific state but is co-expressed with other genes, reinforcing the hypothesis of a transitional state.

***Supp. note 3: Pseudotime-based epigenetic reconstruction of the acinar-to-ductal transition and integration with snRNA-seq data***

Similarly to the snRNA-seq data, we performed a STREAM-based pseudotime analysis on the snATAC-seq dataset. We first pre-selected cells from the Acinar 4 cluster and ductal cells exhibiting a clear hybrid phenotype (see Methods) and analyzed them together (Supp. Fig. 6A). To interpret the underlying regulatory programs, we leveraged gene activity scores inferred from chromatin accessibility. This analysis revealed a trajectory with a similar, but more branched structure compared to the RNA-based pseudotime, (Fig. 2E) delineating distinct acinar and ductal arms (Supp. Fig. 6B). These were marked by canonical genes, *CFTR* and *ALDH2* for the ductal branch, and *CELA2A, CELA3B, REG1A, REG3A* for the acinar branch. Interestingly, we observed the enrichment of known centroacinar-associated and progenitor-like markers, including *SOX9, NKX6-1*, and *ELF3* (*119*), within the ductal arm, suggesting the presence of a plastic or transitional cell state within this lineage. This interpretation is further supported by epiCHAOS (*120*), a chromatin entropy metric that quantifies epigenetic heterogeneity, which identified the root cluster (i.e., cluster 4, Fig. 6A) as the most heterogeneous region in the trajectory (Supp. Fig. 6C), suggesting that these cells occupy a highly plastic state, consistent with a transitional or progenitor-like identity. Additionally, enhancer driven gene regulatory network analysis identified *ELF3* as a central hub within the ductal lineage (Supp. Fig. 6D), regulating a broad set of epithelial and plasticity-associated genes, including *KRT19*, *CLDN7*, *CFTR*, and *ANXA13,* which was also found in the snRNA-seq data. This highlights *ELF3* as a potential transcriptional regulator of the plastic centroacinar-like state.

To explore the temporal relationship between transcriptional and chromatin dynamics, we aligned the pseudotime trajectories from snRNA-seq and snATAC-seq using cellAlign (*121*), which applies a dynamic time-warping algorithm to identify matched points across modalities. We focused on the shared portion of the trajectory preceding the acinar–ductal bifurcation (see Methods), where structural consistency was observed across datasets (Supp. Fig. 6E). The alignment revealed a strong correspondence between modalities, while also identifying local shifts where chromatin accessibility either preceded transcriptional changes, reinforcing the notion of regulatory priming, where chromatin remodeling anticipates gene expression during acinar-to-ductal transition.

To explore the biological programs associated with the observed differentiation trajectories, we performed pathway enrichment analysis on marker genes along the STREAM-defined branches. This revealed enrichment for pathways related to cellular stress and plasticity, including chaperone-mediated autophagy, VEGF signaling, and Notch-associated networks, as well as processes involved in protein folding, adhesion, and growth (Supp. Fig. 6F).

***Supp. note 4: Glucose-dependent signal overlaps with T2D-associated signal***

Pancreatic islets maintain glucose homeostasis, and the disruption of their action can have a critical effect on health, and be responsible for T2D progression. Therefore, we next investigated the relationship between beta cell T2D-associated genes and glucose-responsive genes. We observed that gene accessibility at higher glucose concentration was negatively correlated with gene accessibility in T2D beta cells (Supp. Fig. 12J). This relationship was not recapitulated using scRNA-seq. We also detected that 85 consistent glucose-dependent genes in beta cells have previously been linked to T2D. Ultimately, we identified 2 genes associated to both T2D and glucose, consistently across modalities and with genetic evidence, VPS13C -involved in the regulation of mitochondrial function- and SLC2A2 (or GLUT2) -a glucose transporter-, pointing out key regulators in human health.

***Supp. note 5: Description of fetal populations***

We used canonical cell type marker genes to annotate the main epithelial and non-epithelial cell types: acinar (*RBPJL, CPA1*), ductal (*CFTR, BICC1*), endocrine (*INS, GCG, SST, GHRL, PPY*), mesenchymal (*COL3A1, CDH19, WT1*), endothelial (*PECAM1, FLT1, FLT4*), immune cells (*PTPRC*), proliferative cells (*MKI67*) and neuronal cells (*PRPH*), (Fig. 1E, Supp. Fig. 13 B-D, Supp. Table 15).

Detailed analysis enabled the identification of 44 distinct cell types/states (Supp. Fig 13B). Among these were transitional populations bridging the ductal and acinar compartments (PreAcinar_1 and PreAcinar_2), which progressively upregulated acinar identity transcription factors such as *GATA4*, *RBPJ*, and *RBPJL*, which are key regulators of exocrine differentiation. We also identified a small population of Acinar-s cells, which were previously characterized by high expression of digestive enzymes including *CPA1*, *PRSS1* and *CEL* (*13*).

In the endocrine compartment we identified four distinct clusters of endocrine cells: alpha (*GCG*), beta (*INS*), delta (*SST*) and epsilon cells (*GHRL*). Gamma cells, expressing pancreatic polypeptide (*PPY*), did not form a distinct cluster but instead grouped with alpha cells, consistent with prior observations that gamma cells are transcriptionally similar to other endocrine subtypes during development (*122*). We also observed progenitor populations transitioning from ductal epithelium toward endocrine lineage. These were marked by a wave of *NEUROG3* expression, a key driver of endocrine differentiation (*60, 123*). With further differentiation, these cells showed increasing expression of lineage-specifying transcription factors such as ARX, PAX4, and MAFA, consistent with their roles in endocrine subtype specification (*124–126*).

The non-epithelial compartment of the fetal pancreas comprised mesenchymal, endothelial, neural, and immune cell populations. High-resolution single-cell analysis enabled us to identify Schwann cells (*CDH19, SCN7A* and *GRIK3*) and a large population of fibroblasts (*COL3A1, DLC1, LAMA3*), which can be further subdivided into different states of stellate cells. Smaller populations of mesothelial cells (*WT1*, *MSLN*) and vascular smooth muscle cells (*MYH11*, *SYNPO2*) were also detected (Fig. 1E, Supp. Fig 13B-D). Endothelial clusters were divided into arterial, venous and lymphatic cells (*FLT4, MMRN1*). Similarly, we could stratify the immune compartment cells into myeloid (*CD68, ITGAX*) and lymphoid (*CD8A, TRAC*). Finally, we annotated cells of the enteric nervous system expressing *PRPH* and *TUBB3*, and further distinguished inhibitory (*NOS1*, *GFRA1*) and excitatory (*PLXNA4*, *SCUBE1*) neuronal subtypes (*13, 127, 128*).

***Supp. note 6: Proteomics description of the fetal population***

These fetal populations were also observed at the proteomic level. A curated, highly multiplexed dataset (24 markers) generated with CODEX/PhenoCycler Fusion and comprising 224,713 cells was annotated into distinct cell types: mesenchymal, epithelial, endothelial, immune, SYP+CHGA+ elongated, and other groups (Supp. Fig. 14A).The spatial proteomics data added findings indicating that early developmental stages exhibited a higher proportion of mesenchymal cells compared to other cell types. The mesenchymal composition was categorized into two different distinct types (*129*) based on spatial distribution (Supp. Fig. 14B): the peri-pancreatic mesenchyme (PPM) surrounding the pancreas and the intra-pancreatic mesenchyme (IPM) embedded with the other cell types. These two types presented lower and higher *VIM* positive expressions, respectively, in our dataset (Supp. Fig. 14C). Our results showed that proliferative *Ki67+* cells were spatially distributed in the inner part of the pancreatic tissue (Supp. Fig. 14D), with very few positive cells present in the surrounding PPM tissue. There was a higher proportion of proliferative *Ki67*+ cells at early time points, such as 10 wpc and 14.1 wpc (Supp. Fig. 14E).

The fetal pancreatic epithelial compartment, expressing the *PanCK+* antibody (clone A1/A3, which recognizes *KRT10*, *KRT14*, *KRT15*, *KRT16*, and *KRT19*) was detected along the different weeks, where the presence of positive *CHGA+* cells increased throughout the developmental period (Supp. Fig. 14F). By 10 wpc, the *PanCK+* epithelium was arranged in numerous ductal structures, where further investigation of the endocrine compartment revealed a few scarce interspersed cells co-expressing *CHGA*+ and *PanCK*+ markers (Supp. Fig. 15A). The signature endocrine markers (*CPEP*+ canonical beta cells, *GCG*+ canonical alpha cells) were first expressed at time point 11.5 wpc (Supp. Fig. 15B), with the appearance of *SST*+ (delta canonical cells) at 13.3 wpc (Supp. Fig. 15C). Polyhormonal cells were observed from 11.5 wpc to 18 wpc where they co-expressed a large combination of markers, showing a high prevalence of *CPEP*+ and *CGC*+, *CPEP*+ and *SST*+ (*130*) and *GCG*+ and *TTR*+, along with *CHGA*+. Multiplexed images from week 14.1 and week 15.4, which merged 4 endocrine markers and 2 additional neuroendocrine protein signatures (*CHGA*+ and *SYN*+), revealed ample heterogeneity regarding their protein expression levels, with larger polyhormonal populations in the latest week (Fig. 5A-B). By 14.5 wpc, endocrine-positive cells appeared to start clustering together, with *CPEP*+ cells organized in the core and surrounded by *GCG*+ cells and *SST*+ cells, displaying heterogeneous expression (Supp. Fig. 16A). We identified, at 17 wpc, a low *CHGA*+ cell population that was negative for the signature markers *CPEP*, *GCG*, and *SST*, indicating the presence of another endocrine cell group not covered by our current multiplexing antibody panel (Supp. Fig. 16B). By weeks 17.3 and 18, the pancreatic islets were more defined in shape, and the different cell populations (beta, alpha, and delta) began to exhibit canonical markers specific to each population (data not shown). We observed an increase in vascularization, as indicated by the detection of CD31+ cells, and noted their spatial distribution across different developmental time points. By 10 wpc, endothelial cells were confined to the IPM region, with no positive detection in the PPM. When endocrine cell populations were clearly detected due to their abundance, the expression of endothelial cells was also more pronounced, demonstrating a clear embedding between these cell types
